# Biologically inspired warning patterns deter birds from wind turbines

**DOI:** 10.1101/2025.04.14.648692

**Authors:** George R.A. Hancock, Heta Lehtonen, Theo Brown, Amine Ejjite, Ossi Nokelainen, Johanna Mappes, Sandra Winters

## Abstract

Wind power has been at the forefront of investment and innovation in renewable energy. However, bird fatalities from collisions with wind turbines present an ecological and social challenge to the growing deployment of wind power. Increasingly, research has focused on utilising the sensory ecology of animals to provide passive or active cues that minimise collision risk by increasing the detectability and/or aversiveness of the turbine blades and towers. In nature, numerous aposematic species use contrasting colours and striped patterns to warn birds of their unprofitability. These common signal elements are effective due to their salience within a wide range of natural scenes, memorability, generalisability across taxa due to mimicry, and exploitation of the innate colour preferences of birds. This begs the question: might biologically inspired turbine blades that mimic aposematic patterns help protect birds by increasing their avoidance of turbine blades? Here, we used a screen-based ‘game’ experimental setup to test the behavioural responses of wild-caught great tits (*Parus major)* to three existing wind turbine patterns (white, red striped, and single black blade) as well as a novel biologically inspired aposematic pattern. Birds were less likely to approach and, when they did approach, took significantly longer to approach patterned compared to uniform white blades. This effect was strongest for our aposematic pattern compared to all other patterns tested, highlighting the utility of our bio-inspired approach. Our work suggests that adding red, black and yellow warning patterns to wind turbine blades could reduce bird collisions with wind turbines.

## INTRODUCTION

Rising international concern over the ecological and economic impacts of climate change and the resulting race to limit global warming to less than 2°C above pre-industrial levels has led to increasing investment and research into low-carbon energy sources to help transition away from fossil fuels (Agreement, 2015; Mitigation, 2011). Of the energy alternatives, wind power has been identified as one of the most promising energy sources due to its wide geographic applicability and relative efficiency (Lu and McElroy, 2023). However, the construction of wind power plants can displace species (Tolvanen et al., 2023; Zwart et al., 2016) and lead to fatalities either from collisions between flying animals and turbines (Johnson et al., 2003; Marques et al., 2014; Schuster et al., 2015) or by forming sensory traps from attraction to the turbine’s colour, as is the case for invertebrates (Long et al., 2011; Voigt, 2021). Bird fatalities from wind turbines occur from collisions with the moving turbine rotor blades or the static tower (Loss et al., 2015; Stokke et al., 2020). Blade collision risk is higher for birds that soar at the same altitude as the blades, such as eagles and vultures (Perrow, 2017; Rushworth and Krüger, 2014; Watson et al., 2018), and birds that migrate through wind farms in large numbers and/or at night (Richardson, 1998). Tower collisions on the other hand have almost exclusively been observed from low-flying ground-nesting birds, such as willow ptarmigan (*Lagopus lagopus*), black grouse (*Lyrurus tetrix*), and capercaillie (*Tetrao urogallus*) (Coppes et al., 2020; Gonzalez et al., 2016; Stokke et al., 2020; Zeiler and Gruenschachner-Berger, 2009). Despite efforts to strategically place wind turbines within lower-risk sites, accelerating expansion of wind turbine deployment will only increase the conflict between energy production and the environment (Balotari-Chiebáo and Byholm, 2024), inciting the need to develop effective mitigation strategies for collisions to preserve protected species.

The question of why birds collide with objects that are highly conspicuous to the human eye challenges our understanding of how birds identify potential hazards, how their visual systems differ from those of humans, and importantly how to resolve this issue. Several factors have been cited as potential causes for bird collisions with wind turbine blades, including failure to detect the wind turbine blades due to low-light conditions (Hüppop et al., 2006), low visual acuity (Martin and Banks, 2023), motion smear from the high blade tip speeds, and limited field of view (Hodos et al., 2001; May et al., 2020). Additionally, birds could fail to recognise turbines as a threat (Simmons and Martins, 2024) or may be unable to manoeuvre safely away from them when within the blade-swept zone (De Lucas et al., 2008; Marques et al., 2014). However, the infrequency of behavioural observations of birds at the time of collision with turbines has made it difficult to identify the cause. Proposals to reduce the risk of bird collisions with wind turbines have increasingly relied upon knowledge of the sensory and behavioural ecology of the species at risk. These include using auditory deterrents for bats or birds (Arnett et al., 2013; Smith et al., 2011) and visual deterrents such as modifications to the wind turbines themselves with reflectors, flashing lights, or painted patterns to increase their conspicuousness (Kerlinger et al., 2010; May et al., 2020; Stokke et al., 2020).

Painting turbines has been highlighted as one of the best possible options for a passive visual deterrent due to its lower dependence on lighting than reflectors, lower likelihood of attracting nocturnal birds compared with flashing lights (Kerlinger et al., 2010), and lower running cost compared to alternative ancillary detection technology from radar or camera-based systems (Marques et al., 2014; Singh et al., 2015). Within the natural world, warning signals – which can be visual, acoustic or olfactory – are commonly used by animals to signal themselves as being unprofitable to attack (Caro and Ruxton, 2019; Mappes et al., 2005). This phenomenon, known as aposematism, is thought to explain the prevalence of conspicuous colourations within numerous unpalatable or ‘dangerous’ taxa. Signals are often more effective when they are more easily recognised and remembered by the predator (Endler and Rojas, 2009; Ruxton et al., 2019), thus aposematic species have frequently evolved colours and patterns that are salient against the variable conditions of natural backgrounds (Aronsson and Gamberale-Stille, 2009; Sillén-Tullberg, 1985; Stevens and Ruxton, 2012) and are similar to those of other local unpalatable species (Mullerian mimicry) (Beatty et al., 2004; Sherratt, 2008). Most terrestrial aposematic colours are likely to be directed at birds given their prevalence as predators (Kassarov, 2003) and the commonly converged evolution of bright red and yellow colours, which are indistinguishable from many natural backgrounds to dichromatic mammalian observers (Lawrence and Noonan, 2018). Behavioural experiments with birds have shown that they can quickly and socially learn to avoid warning colours (Aronsson and Gamberale-Stille, 2008; Landová et al., 2017; Lawrence and Noonan, 2018; Roper and Wistow, 1986; Thorogood et al., 2018) and possess unlearned aversion towards red, orange, and yellow colourations and highly contrasting black patterning, common in aposematic species (Halpin et al., 2020; Lindström et al., 1999; Pegram and Rutowski, 2014; Protti-Sánchez et al., 2023; Schuler and Hesse, 1985). Species particularly vulnerable to collisions with turbines, such as raptors, have also been shown to avoid aposematic patterns of sympatric prey species (Brodie III, 1993; Valkonen et al., 2011).

Despite the prevalence of aposematic colouration, using warning signals on wind turbines has seen limited experimental investigation and no study has used patterns directly inspired by aposematic animals. By using biomimicry (the act of copying nature to solve problems) we might be able to design wind turbine patterns that reduce the likelihood of collisions with the blades. As most birds are tetrachromats and are able to detect varying degrees of UV light, increasing the UV reflectance of wind turbines from their typical 10% UV reflectance (UV dark) to 70% (white) has been field trialled as a cryptic warning (invisible to humans), with inconclusive results (Erickson et al., 2003). Painting both the base of the tower and a single blade of the turbine black has been shown to reduce collision risk for grouse and raptors respectively (May et al., 2020; Stokke et al., 2020). In both cases, black was thought to be more conspicuous against the background than white and in the latter case painting a single blade helps to reduce motion smear based on electroretinogram responses to blades within a laboratory setting (Hodos, 2003). Meanwhile, contrasting red stripes are already applied to wind turbines in some countries, such as Germany, to signify their proximity to airstrips. Given the prevalence of black and red in aposematic signals (Robinson et al., 2023), these patterns may also provide a warning function increasing the aversion of birds to the turbines, but this remains untested. Here, we provide the first behavioural comparison of avian hazard perception in response to turbines with existing blade patterns (white, red striped, and single black blade) and a novel biomimetic pattern inspired by aposematic animals, which includes red, yellow, and black stripes. To do this we used screen-based behavioural assays where birds were tasked with pecking 2-dimensional grey dot-shaped targets inside and outside the radius of the wind turbine blades rotating at the average speed of turbines (20rpm) in exchange for food. We predicted that the biomimetic pattern should decrease the likelihood and increase the latency of birds pecking the targets, particularly those inside the radius of the blades where the perceived risk should be higher.

## METHODS

### Experimental setup

For our experiments, we used wild Eurasian great tits (*Parus major*, n=22) captured and housed at Lammi Biological Research Station (61.0543N, 25.04086E) as a model avian observer. The nearest wind turbine site was over 80km away, so the birds were unlikely to have regularly encountered wind turbines before and thus, having any pre-learned response on our experimental set up. Birds were captured using a trapping system, housed individually throughout our experiments, and provided with food and water ad libitum, except during training and testing where food was deprived prior and returned after, to ensure motivation. During initial training, birds were housed in temporary home aviaries (46 x 68 x 58 cm) before being transferred to one of three larger aviaries (90 x 90 x 79 cm) which also acted as our touchscreen operant chambers (TOCs) (Figure 1) (Seitz et al., 2021; Winters et al., In Prep). All aviaries were custom-built from plywood and contained a perch for the birds to sit and sleep on. The TOCs used high-performance GPU-enabled PCs (Lenovo Legion T5 Ryzen 7 with 16GB RTX 3070) and gaming monitors (Asus ROG Swift PG329Q 32” at 165 Hz, 2560 x 1440 resolution) to present moving stimuli with a frame rate higher than the critical flicker fusion frequency (CFF) of the most closely related species for which the CFF is known, the blue tit (*Cyanistes caeruleus*) which sees at up to a maximum of 131 Hz (Boström et al., 2016). Each screen was calibrated using a Calibrite ColorChecker Studio (XRITE, Grand Rapids, MI, USA). To allow the birds’ beaks to interact with the screens we used an infrared touch frame (G6 Integration Kit Touch Frame 6TP 32”, 250 fps maximum). As the birds were unable to navigate the screen by flight alone, a steel grid mesh was positioned in front of the IR frame to allow the bird to access the entire screen area. A 3mm thick acrylic sheet cut to the same dimensions as the monitor was placed between the infrared frame and the screen to protect it from damage. All digital stimuli were rendered using the Psychtoolbox-3 for MATLAB (Kleiner et al., 2007). All MATLAB code and images used to construct the digital stimuli are available within our supplementary material. A full breakdown of our TOC apparatus can be found in (Winters et al., In Prep) win. Bird training and experiments were carried out between January – April 2024.

**Figure 1.**
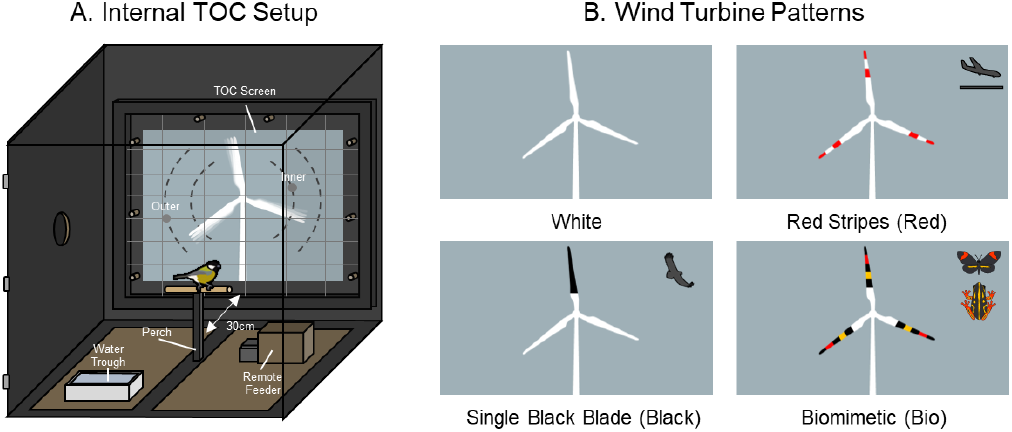
(A) Illustration of internal touch operant chamber (TOC) setup. Wind turbine stimuli during the final experiment were presented on the screen with two target dots, one inside the rotor-swept area of the turbine blades (inner) and the other outside (outer). The dashed lines show the possible locations where the inner and outer dots could occur. (B) Wind turbine stimuli are shown to scale with their backgrounds and the inspiration for their function shown on the right for the three patterned wind turbines (red = aviation warning, black = raptor deterrent, and bio = inspired by warning signals).

### Turbine Design

Each turbine consisted of a live 2D render against a blue-grey background (159, 176, 183 RGB; chosen to approximate the color of an overcast Finnish sky) with a 20cm tower and 16cm blades (Figure 1). The blade shapes were constructed by tracing real-world images of wind turbines and for each turbine the white regions were (255,255,255 RGB). We used four turbine designs: all white (255, 255, 255 RGB; ‘white’), two red (255, 0, 0 RGB) stripes on the outer portions of all blades (‘red’), a single black (0, 0, 0 RGB) blade (‘black’), and a biomimetic design (‘bio’). For the biomimetic blade pattern, we used a red (255, 0, 0 RGB), yellow (255, 192, 0 RGB), and black striped pattern along each blade, inspired by the patterns frequently observed in aposematic species (Figure 1)(Brodie III, 1993; Exnerová et al., 2003; Stevens and Ruxton, 2012; Valkonen et al., 2011). For our biomimetic pattern, we picked a spatial scale of stripes that was both similar equal to the red stripe pattern of the ‘red’ turbine (1.91cm) and which would be visible from the maximum viewing distance of the TOC (100cm) when modelled with the acuity of similarly sized members of the family Paridae (5 cp/d) (Caves et al., 2024; Moore et al., 2013). As neither the screens nor lights used for the aviaries emitted UV light, the white blades would have been similarly UV dark to actual wind turbines, although the background would have lacked the high proportion of UV light within a natural sky. The colours of aposematic species are also frequently UV dark (Barnett et al., 2018; Crowell et al., 2024; Lyytinen et al., 2001).

### Initial Training

Before being exposed to wind turbines, the birds were trained to peck 1.75 cm dark grey dot stimuli in exchange for a food reward (Silvasti et al., 2021). Training birds to reliably peck a target stimulus allowed us to then identify how turbine appearance and relative location influenced their willingness to do so. Grey dots were chosen as a generic stimulus unlikely to interact with particular turbine patterns. First, birds were trained to associate a grey printed paper dot with food by placing a sunflower seed in a slit inside the dot to be retrieved. After the birds successfully fed from these dots twelve times, the seeds were then glued underneath the dots for a further twelve times to force the birds to peck and rip through the printed dot to access their food. Birds were then trained to feed from an open food reward box containing half-cut sunflower seeds before being moved to one of the TOC aviaries. They were then trained to associate pecking at a printed dot with the food reward box opening remotely and retrieving food from it. These physical dots were then gradually moved further from the feeder until the bird was able to peck them on the screen against the blue-grey background. As some birds struggled to peck at dots on the screen a Perspex cover was placed over the printed dot for some of the trials to ensure they would still try to peck the dot even if they could not physically touch it. Lastly, birds were trained to peck at a digital dot (114, 114, 114 RGB) on the touchscreen instead of a physical dot. Birds were also trained to associate landing on the central perch and facing the screen with activation of the trials. Once the birds had successfully pecked dots in all four quarters of the screen, five times each and in less than two minutes per dot, they were allowed to move on to the experiments. 10 of the 32 birds captured failed to finish the initial training within five days and were released and ringed to prevent recapture.

### Turbine Training

Pilot testing showed that naive birds were unwilling to approach a novel turbine stimulus with blades spinning at full speed, necessitating a training stage to facilitate acceptance of this novel stimulus. To prevent potential confounds that could arise from training on alternative turbine appearances, each bird was randomly assigned one of the four wind turbine patterns for training (white N = 5; red = 5; black N = 6; bio N = 6); in later experimental trials, this training turbine pattern was therefore ‘familiar’, and the other three patterns were ‘novel’. For each turbine training trial, birds were tasked with pecking a single gray dot in the presence of turbines rotating at various speeds (0, 2.5, 5, 10 rpm), starting with a stationary turbine (0 rpm) then proceeding through additional speed increases in order. Birds had to peck the dot within two minutes of it appearing. The dot could appear along an arc to the right or left of center of the rotor, either inside (radius = 14 cm from center, inner) or outside (radius = 20 cm from center, outer) of the rotor-swept area of the turbine blades. Inside dots were rendered behind the blades (i.e., the blades covered the dot as they rotated). Each bird had to successfully peck 5 inner and 5 outer dots for each speed before advancing. Once the dot was clicked, the screen turned grey (127, 127, 127 RGB) and a food reward was given. Dots were randomly ordered such that an inner or outer dot could not occur more than twice in a row. The starting orientation of the blades was randomised for each trial and the blades always spun clockwise. For each trial we recorded the bird ID, time, date, order (trial number for that bird), time taken for the bird to peck the dot, whether the dot was in/out, left/right, and whether the bird timed out. Trials were monitored by the experimenter using a viewing window and recorded using a camera mounted inside the TOC aviary (GoPro Hero11 Black, San Mateo, California, United States). In some trials, an issue with the touch-frame encoding for the Psychtoolbox (later resolved, see supplementary material) prevented a peck on the dot from registering; in these cases, a key press override was instead used by the experimenter to trigger the end of the trial. The time between the onset of the trial and the peck on the dot was recorded by MATLAB and with a stopwatch as backup. Trials were typically shown in blocks of 4-6 trials back-to-back depending on how long the birds maintained motivation, with 20-30 min breaks in between each block. Birds that completed trials but did not take any food were considered to no longer be food-motivated and were given a break. Birds that completed the criterion at 10 rpm proceeded to the blade colour comparison experiment.

### Blade Colour Comparison Experiment

In the blade colour comparison experiment, we assessed the responses of each bird to each of the four turbine patterns. Before the experiment commenced, each bird was presented with a series of control trials using the same wind turbine pattern it was trained with (‘familiar’ pattern). Unlike the training experiment, for the control trials both an inner and an outer dot were shown simultaneously, and the speed of the blades was set to 20 rpm. The birds could now peck either the inner or the outer dot, resulting in the same grey screen and food reward once one of the dots was pecked. We recorded the dot they selected (inner or outer), then continued the trial after the bird had eaten and resumed its position on the perch, with the same stimulus being shown again but with only the un-pecked dot remaining. Each comparison trial with a given turbine appearance therefore consisted of two (potential) pecks at dots, with a brief break in between. Birds had to complete a minimum of four control trials (8 dots), with three successful trials in a row before moving on to the final comparison experiment. A trial was considered successful if at least one of the two dots was pecked. Once the control trials were completed, the birds were exposed to all four blade patterns, including the familiar training pattern and three novel ones. Trials were conducted in five blocks of four, with each block consisting of the four turbine patterns, randomly ordered using pre-generated Latin squares. After completing the experiment birds were released and colour-ringed to prevent recapture.

### Statistical Analyses

For our analyses, we compared the effect of the wind turbine patterns and target positions (outer = low risk & inner = high risk) on bird behaviour with R version 4.3.2 (R Core Team, 2023) using linear mixed models (LMMs) and binomial generalised linear mixed models (GLMMs) with the “ lme4” package (Bates et al., 2015). For the training experiment, we tested whether the turbine pattern and target position as well as the (factorial) speed of the blades and the trial number influenced the proportion of timeouts (when birds did not peck a dot within two minutes; binomial model) and the capture time for successful target pecks (linear model). This approach first assesses what affects birds’ willingness to approach the turbine, then for birds that do approach, what affects their peck latency. For the comparison experiment, we likewise tested the effects of pattern and target position as well as novelty (was the blade novel or familiar), trial number (1-20), and whether the dot peck was first or second on capture time (linear model) and the proportion of timeouts (binomial model). We also tested the effect of these predictor variables on the proportion of outer dots chosen first (binomial model) using a subset of the data that only included the first part of each trial (i.e., the first out of two dot presentations). To compare differences between factor levels, e.g. wind turbine colour or speed, we used emmeans Tukey post hoc tests (https://CRAN.R-project.org/package=emmeans).

For each test, we constructed a base model containing all fixed effects and then used likelihood ratio tests to remove non-significant terms from the model in a stepwise fashion. Retained variables were then tested for significant interactions by comparing the fit of models with different levels of interactions to a model without interactions. For each model, the ID of the bird and the TOC number were included as random effects, and trial ID (trial number + bird ID) was used as a random effect for the comparison experiment as each trial contained two dot pecks. For comparisons of capture time, the log time was used as it was closer to a normal distribution than the raw time. For all models, numeric variables were rescaled to a mean of zero and a standard deviation of 1. Residual plots were used to verify that all models met assumptions.

## RESULTS

### Turbine Training Exercises

For the training results only the first 10 trials for each speed were used in our analyses as some birds required additional trials to meet the criteria of 5 successes for inner dots and 5 successes for outer dots. For the binomial model assessing timeouts, only speed and trial number were retained as effects, meanwhile for log capture time, speed, dot type and trial number were retained. Blade pattern had no significant effect on timeouts or capture time during training. However, birds on average did take longer to respond to the appearance of dots for patterned blades (red, black and bio) compared to white when training commenced (speed = 0, see supplementary material). Later trials were significantly less likely to timeout (β =−0.407, t= −3.064, p=0.0022) and had shorter capture times (β = −0.091, t_749.9_ = −2.757, p = 0.006) as the birds became more habituated to the blades. Birds took significantly longer to peck outer dots compared to inner dots (β = −0.2115, t_752.54_ = −2.757, p = 0.00124) The number of timeouts did not increase across speeds as was expected. Post Hoc comparisons between speeds showed that birds were significantly more likely to timeout at 2.5rpm compared to 0 rpm (Table 1) and took significantly longer to capture dots for 2.5 rpm compared to all other speeds (Figure 3A, Table 1). This was likely in response to the novelty of motion (2.5 is the first speed jump after 0) as capture time declined with repeated exposure to moving blades (Figure 2B).

**Table 1.** Results for emmeans Tukey posthoc comparisons of wind turbine speed’s effect on the likelihood of a timeout and capture time during the turbine training exercise. Negative estimates indicate that speed A has a lower likelihood of a time out / capture time than B, while positive values indicate that A is higher than B.

| Speed A | Speed B | Timeout |  |  |  | Capture Time |  |  |  |
| --- | --- | --- | --- | --- | --- | --- | --- | --- | --- |
|  |  | Estimate | SE | Z value | p value | Estimate | SE | t value | p value |
| 0.0 rpm | 2.5rpm | -1.372 | 0.395 | -3.47 | 0.0029 | -0.3844 | 0.0928 | -4.140 | 0.0002 |
| 0.0 rpm | 5.0 rpm | -0.923 | 0.406 | -2.28 | 0.1308 | 0.0882 | 0.0915 | 0.964 | 0.7701 |
| 0.0 rpm | 10.0 rpm | -0.593 | 0.418 | -1.42 | 0.4867 | 0.0104 | 0.0908 | 0.115 | 0.9995 |
| 2.5 rpm | 5.0 rpm | 0.449 | 0.333 | 1.35 | 0.5308 | 0.4726 | 0.0942 | 5.018 | <.0001 |
| 2.5 rpm | 10.0 rpm | 0.779 | 0.351 | 2.22 | 0.1178 | 0.3948 | 0.0932 | 4.234 | 0.0002 |
| 5.0 rpm | 10.0 rpm | 0.330 | 0.64 | 0.90 | 0.8026 | -0.0778 | 0.0923 | -0.843 | 0.8340 |

**Figure 2.**
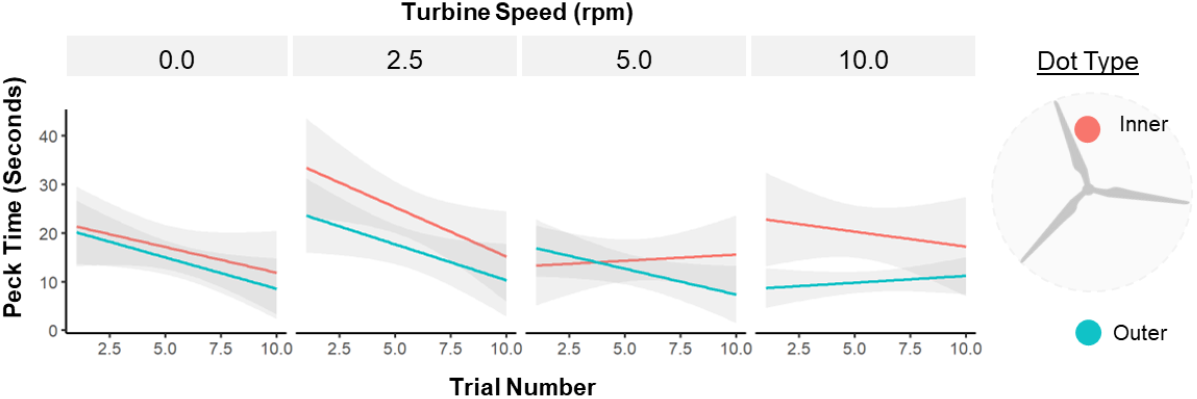
Effect of trial number, turbine speed and dot type, on the time taken for birds to peck the dot in seconds. Lines show the fitted linear model for the inner ‘higher risk’ dot (red) and the outer ‘lower risk’ dot (blue). The grey bars show the standard error.

**Figure 3.**
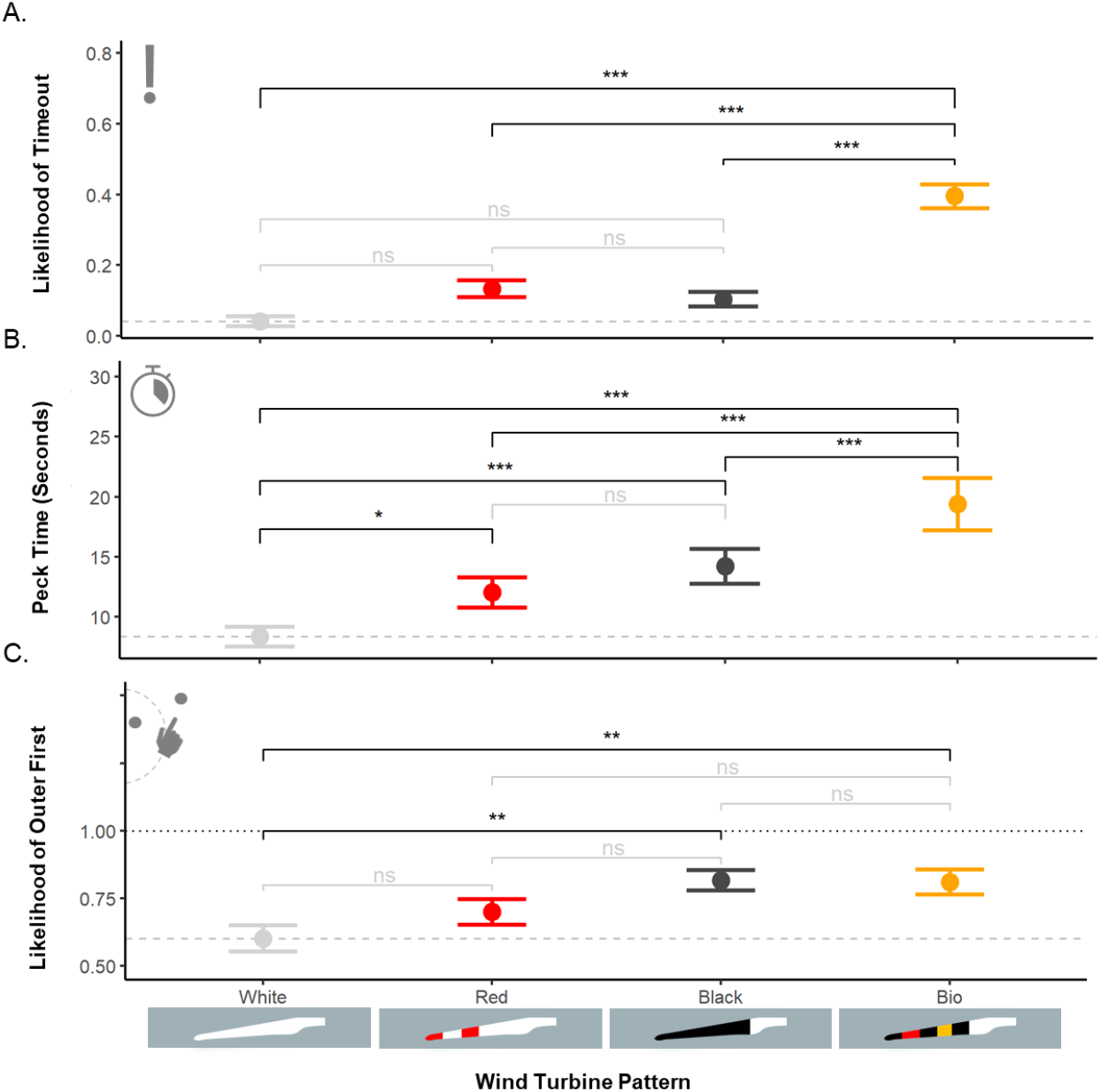
(A) The effect of turbine blade pattern on the likelihood of a bird timing out during a trial, greater values indicate a greater likelihood of a timeout. (B) the effect of turbine blade pattern and pattern novelty (familiar = trained with, novel = not trained with) on the log time for the target dot to be pecked. (C) The effect of turbine blade pattern on the binomial for dot choice, greater values indicate a greater likelihood of choosing the outer dot first (lower risk). For all plots, points show the mean and the error bars show the SE. Brackets indicate the significance of Tukey post hoc comparisons between blade patterns, where ns = not significant, * = p<0.05, ** = p<0.01, and *** = p<0.001. The dashed grey lines indicate the mean for the white un-patterned turbines.

### Blade Colour Comparison Experiment

Of the 440 trials conducted, 14 were excluded due to the bird not engaging in the trial (e.g., bathing, preening, or other signs of inattention) or technical errors. For our final model of timeouts, only turbine pattern, novelty and trial number were retained. Dot type (inside or outside) and whether the target was the first to be pecked were dropped from the final model. The biomimetic pattern was more likely to timeout than any of the other designs; the other three designs were not statistically significantly different from each other (Table 1; percent of trials that timed out: white = 4.19%, red = 13.42%, black = 10.45%, bio = 39.53%). Similarly, familiar blade patterns (β =−7.136, Z= −3.274, p = 0.0011) and increasing trial number (β =−0.221, Z= −3.158, p = 0.0016) lowered the likelihood of a timeout. The biomimetic pattern had a significantly higher likelihood of a timeout than any other pattern and was the only pattern with a significantly higher likelihood than the white control (Figure 3A, Table 2).

**Table 2.** Results for emmeans Tukey posthoc comparison of the wind turbine pattern’s effect on the likelihood of a timeout and capture time during the blade colour experiment. Negative estimates indicate that wind blade pattern A has a lower likelihood of a time out / capture time then B, while positive values indicate that A is higher than B.

| Pattern A | Pattern B | Timeout |  |  |  | Capture Time |  |  |  |
| --- | --- | --- | --- | --- | --- | --- | --- | --- | --- |
|  |  | Estimate | SE | Z value | p value | Estimate | SE | t value | p value |
| White | Red | -3.570 | 1.76 | -2.022 | 0.180 | -0.274 | 0.096 | -2.862 | 0.0231 |
| White | Black | -2.540 | 1.76 | -1.443 | 0.473 | -0.369 | 0.094 | -3.912 | 0.0006 |
| White | Bio | -11.60 | 3.40 | -4.824 | <.0001 | -0.907 | 0.108 | -8.400 | <.0001 |
| Red | Black | 1.020 | 1.43 | 0.714 | 0.8917 | -0.095 | 0.097 | -0.985 | 0.7582 |
| Red | Bio | -8.030 | 1.88 | -4.270 | <.0001 | -0.633 | 0.109 | -5.819 | <.0001 |
| Black | Bio | -9.050 | 2.06 | -4.396 | <.0001 | -0.538 | 0.108 | -4.985 | <.0001 |

As with timeouts for our final model of capture time, the wind turbine pattern shown, the novelty of the pattern, and the trial number all had a significant effect on capture time. While there were significant interactions between novelty and blade pattern, all models with interactions were a poorer fit than the one without interactions, as determined by likelihood ratio tests, and so the interaction term was dropped. We observed shorter delays to peck the targets for both familiar turbine patterns (i.e., the pattern the bird used for training; β= −0.400, t_342.8_ = −5.03, p < .0001) and as trial number increased (β= −0.221, t_336.5_ = −6.127, p < .0001). The biomimetic pattern had a significantly higher capture time than all other blade patterns (mean times: white=8.36s, red=12.05s, black=14.21s and bio =19.41s), while the red and black treatment had a significantly higher capture time than the white but not from each other (Figure 3B, Table2).

Although the dot location (inside or outside of the blade-swept zone) did not statistically impact the likelihood of a bird timing out or their peck time, the outer dot tended to be chosen first, especially for the black and biomimetic blade (outer chosen first: white=60.19%, red=70.10%, black=81.73% and bio =81.08%). In the model analysing the location of the first dot choice, only the turbine pattern was retained as an effect, with both the black and biomimetic patterns significantly increasing the likelihood of the outer dot being chosen first compared to the white (Figure 3C), but not when compared to each other or the red pattern(White-Black: β =−1.157, Z=−3.439, p=0.0033; White-Bio: β =−1.2015, Z=−3.185, p=0.0079; see supplementary material for full table).

## DISCUSSION

Here we provide the first behavioural evidence that incorporating patterns onto wind turbines can increase their aversiveness to birds compared with plain white turbines. This work follows a long history of evidence that contrasting patterns and colours of aposematic animals, in this case combinations of red, black, and yellow, can increase the aversion of birds towards foreign objects (Aronsson and Gamberale-Stille, 2009; Halpin et al., 2020; Schuler and Hesse, 1985) and supports proposals to add striped patterns to wind turbines (Martin and Banks, 2023; Simmons and Martins, 2024). Using patterns inspired by aposematism, we produced a biomimetic wind turbine that birds treated as more aversive than any of the other patterns tested. While our biomimetic pattern had the greatest effect in our behavioral assay – generating more hesitancy to approach the turbines and a longer latency to peck targets near them – the red-striped and singular black blade patterns also increased aversion, but to a lesser extent. Both the biomimetic and singular black blade patterns directed birds to attack targets outside the swept area of the blades when given a choice. Novelty alone is unlikely to explain our observed results for the biomimetic pattern (Barnett, 1958; Marples and Kelly, 1999), as the biomimetic pattern still had a higher capture time when it was familiar(i.e., when the birds had previously trained on this blade appearance) compared to the white and red treatments. While birds were able to habituate to the blades, resulting in less turbine avoidance and faster pack times across sequential trials, the speed of habituation was likely influenced by the presence of a food reward associated with approaching the screen that the turbines were displayed on.

Black patterns have commonly been proposed to reduce the likelihood of collision by increasing the turbine’s contrast against the background, better matching the appearance of natural hazards (e.g. trees) and reducing motion smearing when using offset stripes or singularly painted blades (Hodos, 2003; Martin and Banks, 2023; May et al., 2020; Stokke et al., 2020). The increased aversion to our patterned turbines is unlikely to have been the result of motion smear prevention. At the spatial scale of our experiments, the turbines at 20rpm would have had a visual angle speed of 64.0 va/s when viewed by our birds on the perch (30cm away) and 384.0 va/s when viewed on the wire frame (<5cm away). These visual angle speeds are equivalent to a viewing distance of 187.5 m and 31.3 m respectively when observing a turbine with a 100m blade travelling at the same rpm and falling within the value range where differences in pattern electroretinogram (PERG) response have been observed for different motion smear prevention strategies (striped and mono black blades), between 100 va/s and 240 va/s (Hodos, 2003). PERG measures of American kestrels (*Falco sparverius*) in response to rotating turbines have shown that black and red colours produce higher amplitude responses against blank backgrounds but not natural backgrounds where white produces higher amplitudes (Hodos, 2003). The reduced collision risk of raptors in sites where turbines have a single blade painted black may be driven by an increased aversion to dark moving objects compared to light as opposed to being influenced by absolute contrast (May et al., 2020). Previous experiments have shown various vertebrate taxa are more likely to perform escape behaviours in response to dark-on-light, rather than light-on-dark, moving objects (Temizer et al., 2015; Yilmaz and Meister, 2013). Intermixing dark-striped patterns also has the potential to break up the appearance of wind turbines at long distances reducing their visual impact on humans (Maffei et al., 2013) and increasing their novelty to approaching birds, similar to distant dependent camouflage (Barnett et al., 2018, 2017).

One of the largest barriers to painting wind turbines and testing the effectiveness of different patterns in situ is the difficulty of re-painting wind turbines that are already operational (May et al., 2020). The methods used here to assess the behavioural responses of birds to wind turbine patterns have the potential to be used to test a wide range of additional turbine parameters (speed, pattern, number, background, etc) far more easily and cheaply. As with other computer renderings of threats shown to animals, such as looming objects (Carlile et al., 2006; Temizer et al., 2015; Yilmaz and Meister, 2013), the birds in our experiment appeared to treat the moving turbines as a physical threat risk, despite there being no physical risk of collision. When exposed to a novel blade pattern during the control experiment or rotating blades for the first time during training, the birds frequently fled from their perch and attempted to move as far away from the blades as they could, often with an alarm call. Birds were also observed fleeing and dodging away from the blades when attempting to peck the targets (See Supplementary Videos). The initial spike in target capture time when the blades started moving during training supports the observation that many bird species actively avoid wind turbines, particularly migrating birds which may not have prior exposure to turbines, modifying their flight height and orientation when they encounter wind farms (Johnston et al., 2014; Santos et al., 2022; Schiff, 1965). However, while our experiment’s results align with previous studies with actual wind turbines, further experimental validation in the field is required to ensure the increased aversiveness of our biomimetic aposematic pattern is maintained at larger scales and when observed by high-risk species such as white-tailed sea eagles (*Haliaeetus albicilla*) (Balotari-Chiebáo and Byholm, 2024; May et al., 2013). While large soaring birds, such as raptors, are often the target of wind turbine collision mitigation strategies, passerines, which comprise the majority of bird species, are also vulnerable to collisions with wind turbines and are often under-sampled in wind turbine studies as they are difficult to find and are quickly scavenged (Erickson et al., 2014; Nilsson et al., 2023).

The main reasoning for wind turbines being painted shades of white is usually to minimise visual impact to humans, increase conspicuousness from the air, and to reduce thermal stress on the blades (Eyerly, 2019). Furthermore, some countries also have legislative constraints for the appearance of wind turbine blades that can restrict alterations to the appearance of wind turbine blades. E.g. In Finland where only white turbines are permitted for onshore wind farms. Visual impact on humans and aviation are important considerations for the design and positioning of wind turbines. Despite red-striped patterns already being wildly incorporated and beneficial within the context of aviation, their ecological effect on collision risk has yet to be investigated where they are implemented. Some countries, such as South Africa, have considered using red stripes as a method for reducing raptor collisions (Simmons and Martins, 2024). Yellow bands are already used for offshore wind turbines to increase conspicuousness to ships (Department of Trade and Industry (DTI), 2005) and the addition of stripes to virtual turbines has been shown to reduce the visual impact of turbines for humans when viewed from a distance (Maffei et al., 2013). The distance-dependence of black and yellow stripes as a warning signal (Barnett et al., 2018) might also have the added bonus of reducing habituation to the turbines.

## CONCLUSION

Our findings highlight the value of using biomimicry to develop nature-inspired solutions for nature. Just as blade shapes inspired by humpback whale fins and silent owl feathers have improved turbine efficiency and reduced noise pollution, respectively (Fish et al., 2011; Lilley, 1998), colours inspired by natural warning signals could provide an effective means of lowering the environmental impacts of wind turbines on bird mortality. While the biomimetic pattern we propose functions as a generic example of aposematic colouration there are many components of the pattern that have yet to be investigated such as the optimal spatial frequency of stripes (Barnett et al., 2018, 2017; Hodos, 2003) and the choice and position of the biomimetic colours (e.g., does it need to be yellow and red?). Other impact factors of wind turbines, such as the aesthetic preferences of humans, should also be evaluated when testing future designs (Bishop, 2002; Maffei et al., 2013; Sullivan et al., 2013). Likewise, countries with restrictions on wind turbine painting should consider introducing legislation to allow for wind turbines to be painted with warning patterns, even if it is only at sites close to migration flyways of vulnerable birds.We hope our work encourages future exploration of the influence of aposematic colouration on wind turbine collision mitigation and that our experimental framework provides a means of piloting different turbine pattern regimes and testing their effect on different bird species’ behaviour.

## Supporting information

Supplemental Video - Turbine Speeds

Supplemental Video - Trial Clips

## Ethics

This work was conducted with approval from the Central Finland Centre for Economic Development, Transport and Environment (permit no. 6870/2022). All birds were released into the area where they were caught once they had completed the experiment and had no indication of ill effects from handling or training.

## Acknowledgements

We would like to thank Lammi Research Station for facilities and technical support, Eric Lehtonen and Fiorina Muster for help maintaining and training our birds, and John Niemi for ringing assistance. We thank Renewables Finland for helpful discussions. The study was funded by the Research Council of Finland (grant 349613 to SW) and 1345091 to JM) and The Finnish Society of Sciences and Letters (to JM and SW). The TOC system was developed with support from the Wild Animal Phenotyping (WildAP) infrastructure funding.

