## Supplemental Information for "Biologically inspired warning patterns deter birds from wind turbines"

### Supplementary Material for "Biologically inspired warning patterns deter birds from wind turbines".

#### Acuity Rendering and Turbine Pattern

To ensure that our striped patterns would be resolvable to birds within the TOC we used the acuity view function within the micaToolbox (Caves and Johnsen 2018; Van Den Berg et al. 2020). Both the red-striped patterns copied from real wind turbines and our biomimetic stripes would have been visible from any viewing position (max distance = 100 cm) within the TOC and even far beyond it (Figure S1). This should be unsurprising as for the length of the stripes (1.91cm), a viewing distance of greater than 5.47 m is required unresolvable to an animal with an acuity of 5 cycles/degree.

*Figure S1. Acuity view rendering of striped turbines (Red and Bio) at different viewing distances, 30cm, 100cm, and 500cm, for a spatial acuity of 5 cycles/degree. The calculated angular width of the screen is included.*

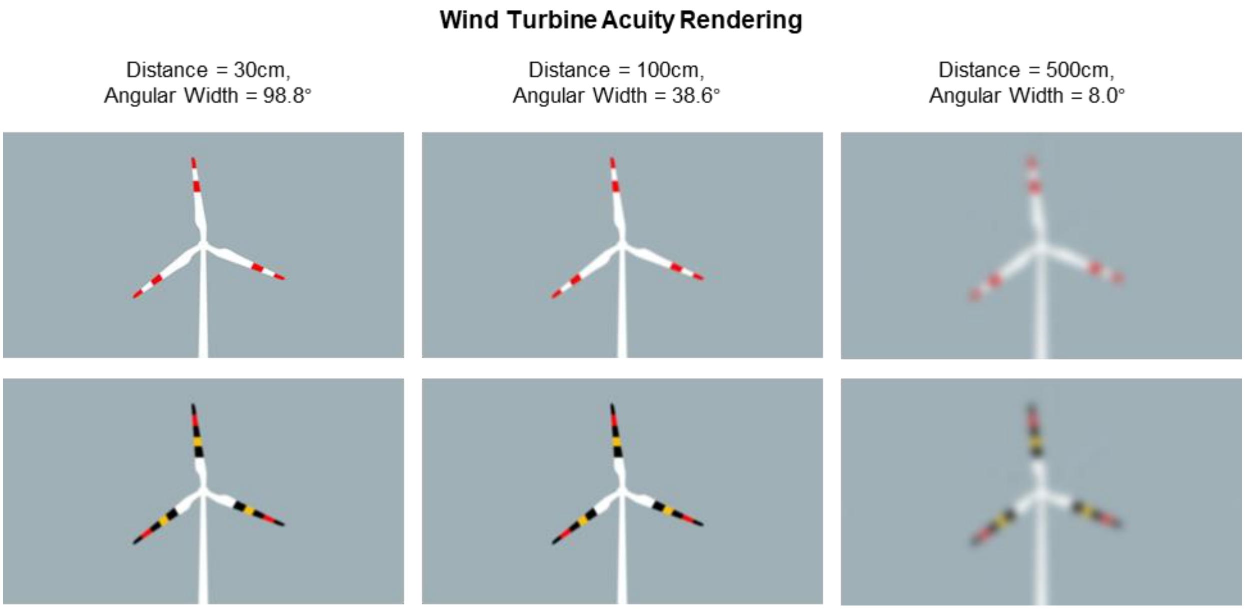

#### Turbine Luminance Contrast

*Table S1. Weber contrasts of the different wind turbine elements against the background. NB: Values are approximated using the screen's calibrated CIELAB space and are unlikely to match the exact values. Luminance values were calculated by measuring excitation of the blue tit double cone (DBL) in the micaToolbox. Weber's contrast was calculated using ( $L_{TurbineColour} - L_{Background}$ ) / $L_{Background}$ . Michelson's contrast was calculated using ( $L_{Max} - L_{Min}$ ) / ( $L_{Max} + L_{Min}$ ).*

*NB the screen minimum and maximum values are assumed to be 0.05 and 0.95, rather than 0 and 1.*

| ***Item*** | ***Colour*** | ***sRGB*** | | | ***Blue Tit DBL*** | ***Weber's Contrast*** | ***Michelson's Contrast*** |
|---|---|---|---|---|---|---|---|
| | | *R* | *G* | *B* | | | |
| *Background* | | *159* | *176* | *183* | *0.47* | | |
| *White* | | *255* | *255* | *255* | *0.95* | *1.02* | *0.34* |
| *Red* | | *255* | *0* | *0* | *0.28* | *0.40* | *0.25* |
| *Black* | | *0* | *0* | *0* | *0.05* | *0.89* | *0.81* |
| *Yellow* | | *255* | *192* | *0* | *0.61* | *0.30* | *0.13* |

#### Turbine Training and Turbine Pattern

While we did not find a significant difference in peck time or the number of timeouts between training patterns, on average, the time taken to peck was greater for the patterned blades (red = 19.77 s, black = 14.75 s, and bio = 17.93 s) compared to the white (9.25 s) un-patterned blades at 0.0 rpm (Figure S2). Had we measured the time taken to peck dots without a turbine as a control, we would likely have observed a significant effect the effect was otherwise overpowered by the bird ID (Table S2).

*Table S2. Results for emmeans Tukey posthoc comparison of the wind turbine pattern's effect on log dot peck time during training and when the speed = 0.0rpm, i.e., the start of training. For this model BirdID has been excluded as a random effect. Negative estimates indicate that wind blade pattern A had a lower dot peck time than B, while positive values indicate that A is higher than B.*

| Pattern A | Pattern B | Estimate | SE | t. value | p value |
|---|---|---|---|---|---|
| White | Red | -0.653 | 0.198 | -3.302 | 0.0062 |
| White | Black | -0.465 | 0.195 | -2.388 | 0.0827 |
| White | Bio | -0.517 | 0.187 | -2.762 | 0.0316 |
| Red | Black | 0.188 | 0.197 | 0.955 | 0.7748 |
| Red | Bio | 0.136 | 0.189 | 0.718 | 0.89 |
| Black | Bio | -0.052 | 0.186 | -0.28 | 0.9923 |

*Figure S2. Influence of wind turbine colour pattern on log peck time during turbine training exercises. Each point shows the mean peck time in seconds, and the error bars show the standard error. The dashed line indicates the mean for the white turbine-trained birds at 0.0 rpm.*

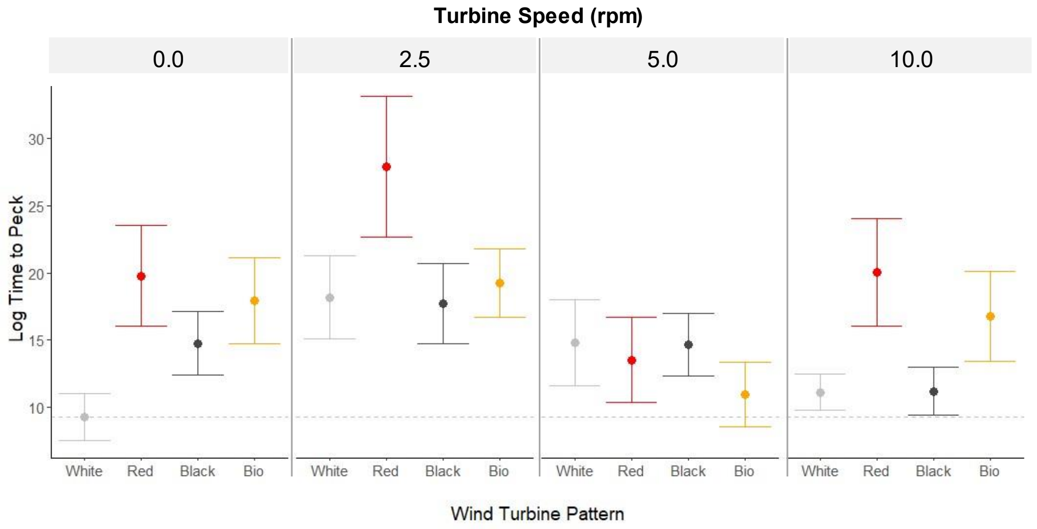


#### Dot Choice Emmeans Table

*Table S3. Results for emmeans Tukey posthoc comparison of the wind turbine pattern's effect on first dot choice. Negative estimates indicate that wind blade pattern A has a lower likelihood of choosing the outer dot first than B, while positive values indicate that A is higher than B.*

| Pattern A | Pattern B | Estimate | SE | Z value | p value |
|---|---|---|---|---|---|
| White | Red | -0.4841 | 0.313 | -1.545 | 0.4103 |
| White | Black | -1.1574 | 0.337 | -3.439 | 0.0033 |
| White | Bio | -1.2015 | 0.377 | -3.185 | 0.0079 |
| Red | Black | -0.6733 | 0.348 | -1.933 | 0.214 |
| Red | Bio | -0.7174 | 0.385 | -1.863 | 0.2442 |
| Black | Bio | -0.0441 | 0.403 | -0.109 | 0.9995 |

#### Touch Screen Registration

During our experiment, we were unable to get the Psychtoolbox-3's touch screen function to work properly. After the first trial, the click registration for the touch screen. This was because we failed to clear the touch screen queue between each trial. Flushing the touch event after each trial loop helped to fix this issue for later projects as did creating a new function for obtaining touch events.

```
TouchQueueStop(game.touchDev);

TouchEventFlush(game.touchDev);
```

##### Dot Spawn Positions

Inner dots spawned 512 px from the centre and outer dots spawned 731 px from the centre. For both dots the angle range was between -35 and +35 degrees from the centre right (0 degrees) or the centre left (180 degrees).

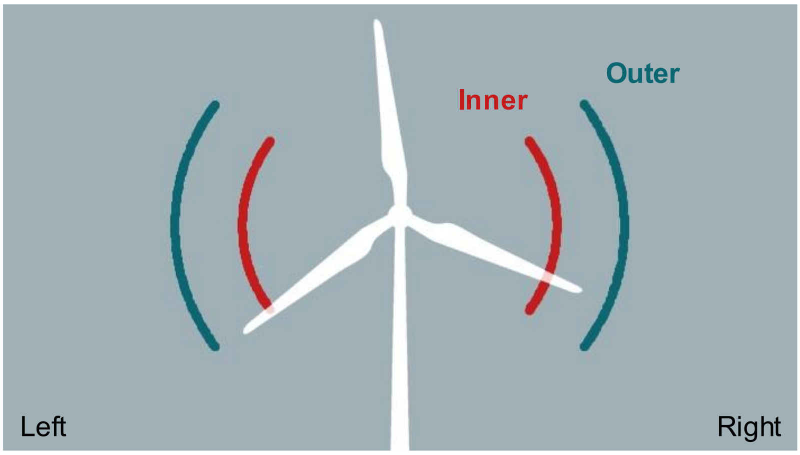


*Figure S3. Range of spawn positions for the inner 'higher risk' dots (red) and the outer 'lower risk' dots (blue). Dots were rendered below the wind turbine blades .png allowing them to pass over them as if they were an object behind the wind turbine.*

##### TOC (Touchscreen Operant Chamber) Images

*Interior Image*

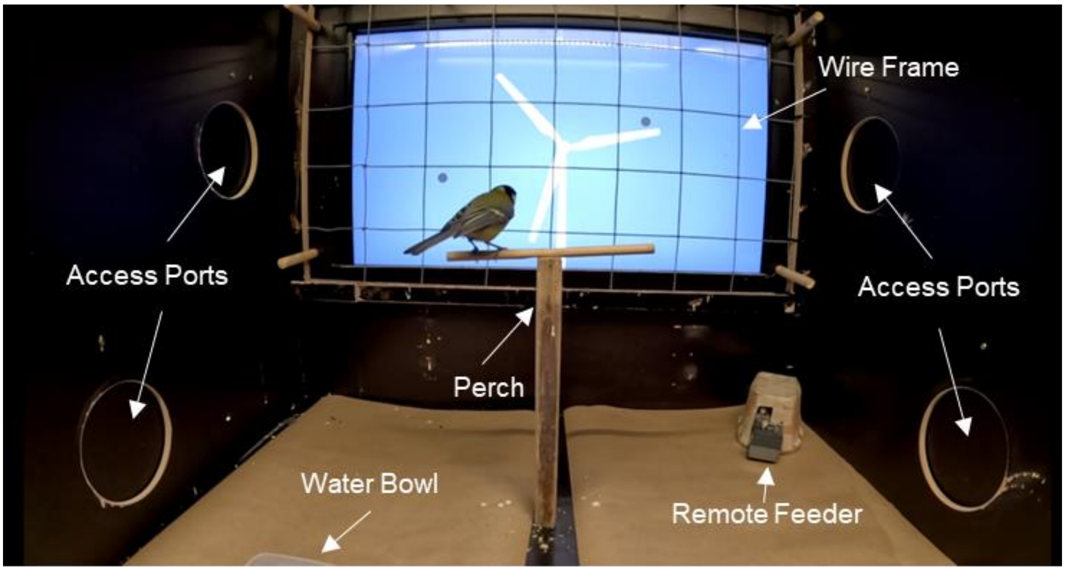

*Exterior Image*

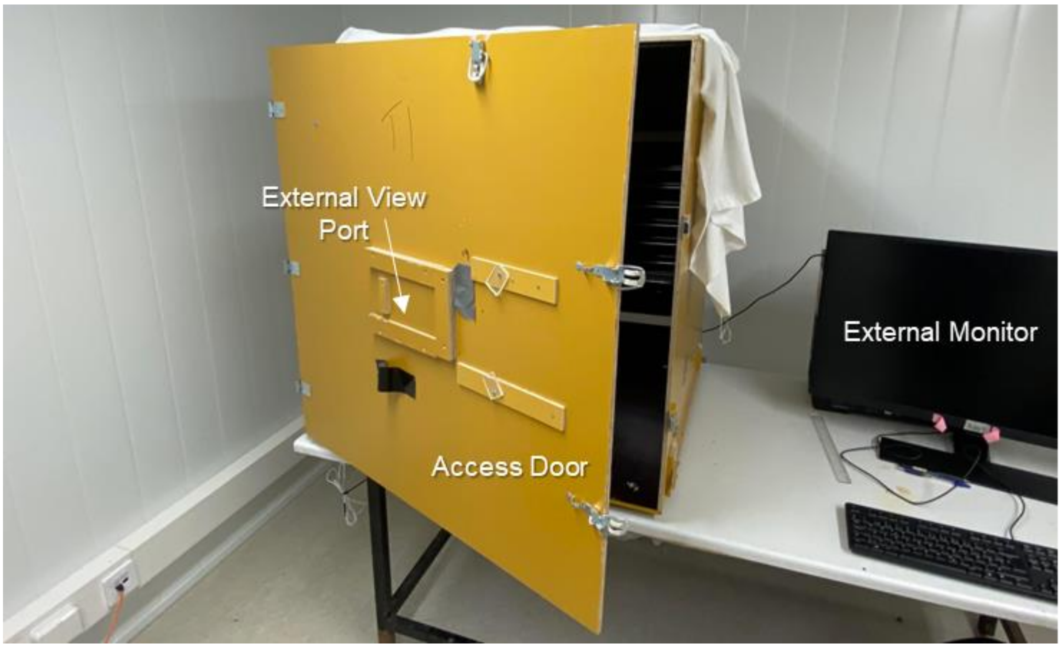
