## Supplementary material for "Biologically inspired warning patterns deter birds from wind turbines": README_file.md

This README.txt file was generated on 2025.04.02 [yyy,mm,dd] by George RA Hancock

# 1) GENERAL INFORMATION
---------------------------------------------------------------------------------------------------------------------------------------------------------

1. Title of Dataset: Biologically inspired warning patterns deter birds from wind turbines.

2. Author Information
	A. Principal Investigator Contact Information
		Name: George Hancock
		Institution: University of Exeter / University of Helsinki
	

3. Date of data collection. January 2024 - April 2024 

4. Data collection location: Lammi Biological Research Station, Pääjärventie 320, 16900 Lammi, Finland

5. Funding Body: Research Council of Finland and The Finnish Society of Sciences and Letters

6. Software recommendations. 

	Statistical Analyses

	- R version 4.3.2+ [https://www.r-project.org/]
	

	Wind Turbine Bird Game
	(NB, MATLAB requires a software lisence, unfortunatley our code is not compatible with Octave).

	- MATLAB [https://www.mathworks.com/products/matlab.html]

	- Psychtoolbox-3 [https://psychtoolbox.org/]
	
	- Windows 10 / Windows 11
	
	- Two monitors, one of which can run at 165 fps.
	
	 NB, screen switching may not work properly with a laptp.
	


# 2) DATA & FILE OVERVIEW
---------------------------------------------------------------------------------------------------------------------------------------------------------

## 1. File List (In Alphabetical order):	

    **/Stats_&_Data_Output/

	*WT_Data_Exp.csv
	
	- Contains the data collected from the wind turbine blade comparison experiment.

	*WT_Data_TOC.csv
	
	- Contains information for which TOC each bird belonged to.

	*WT_Data_Training.csv
	
	- Contains the data collected from the wind turbine training exercises for 0 rpm to 10 rpm.

	*WT_R_Code.csv
	
	- Contains the R code for performing the stats present in the main text and supplementary material.


    **/WindTurbine_Code/

	*/Backgrounds/
	
	- Stores background images used during training and for the experiment.

	*/Birds/
	
	- Stores data for birds, included is an example folder for bird ID 24G03 which was trained with a white pattern.
	- Contains:
		o.Folders with the output for different experimental stages
		o.Text file storing the training pattern (White).
		
	*/Blades/
	
	- Stores blade images for the wind turbines
	
	*/nircmd/
	
	- Used to switch primary monitors during the experiment
	
	
	MATLAB CODE
	
	*a_AddBird.m
	
	- Used to create a folder for a bird and to set its training pattern.
	
	*a_SetupScreens.m
	
	- Makes sure the screens are in the correct configuration before starting the experiment.
	
	*b_WindmillRandDotTrain.m
	
	- Execute this script to start training a bird to peck a dot on the screen, peck times will be output in command line.
	
	*c_WindmillTrainingExperiment.m
	
	- Execute this script to perform the wind turbine training exercises, the user can select which bird they are using and the training stage. 
	- Inner and outer dots are manually set with key commands.
	- Data is output as .csv files in a seperate folder for each speed/stage (stages 0-3).

	*d_WindmillExperiment.m

	- Execute this script to perform the multi dot trials at 20 rpm and the blade comparison trials.
	- Wind turbine pattern is manually chosen with a number key for the comparative trials, numbers were selected using latin square tables.
	- Data is output as .csv files in a seperate folder for each stage ( 4 multidot training and 5 comparative).
	- Pressing the i key registers a hit for the inner dot, pressing the o key registers a hit for the outer dot.
	
	*mergeCSV_x
	
	- These scripts were created to help merge all of the wind turbine training and experiment data into one .csv folder.

## 2. Relationship between files, if important: 


    **/Stats_&_Data_Output/
     To run the R code, the .csv files and R code must be in the same folder.


    **/WindTurbine_Code/
     To run the R code all of the code and files must be kept in the same folder and the file structure shown here.


## 3. Additional related data collected that was not included in the current data package: N/A

## 4. Are there multiple versions of the dataset? NO
	A. If yes, name of file(s) that was updated: N/A
		i. Why was the file updated? N/A
		ii. When was the file updated? N/A


# 3) METHODOLOGICAL INFORMATION
---------------------------------------------------------------------------------------------------------------------------------------------------------

## 1. Description of methods used for collection/generation of data: 

    ** Study Species
	
	Eurasian great tits *Parus major*
	- n = 22 
	- location = Lammi Biological Research Station (61.0543N, 25.04086E) 
	- Capture method = baited trap
	- Housing: home aviary (46 x 68 x 58 cm), TOC aviary ( 90 x 90 x 79 cm).

    ** TOC (touchscreen operant chamer) Setup 
	- High-performance GPU-enabled PCs (Lenovo Legion T5 Ryzen 7 with 16GB RTX 3070) 
	- Gaming monitors (Asus ROG Swift PG329Q 32" at 165 Hz, 2560 x 1440) 
	- Callibrated with a Calibrite ColorChecker Studio (XRITE, Grand Rapids, MI, USA).
	- Infrared touch frame (G6 Integration Kit Touch Frame 6TP 32”, 250 fps maximum). 
	- 3mm thick acrylic sheet cut to the same dimensions as the monitor 
	- Plywood hull ( 90 x 90 x 79 cm).
	- Water bowl.
	- RC food reward box.
	- Strip lights
	- TOCs were housed in temperature controlled rooms.
	
	 ** Bird Games
	Games used to train our birds were created using custom code for Psychtoolbox-3 for MATLAB. Each game included one or two dots which the bird was tasked with pecking in exchange for 
	a food reward. Upon pecking one of the dots the trial would end and the experimenter would open remotely a food reward box.

    ** Data Collection
	All data collected was generated from our MATLAB code for our bird training and comparative experiment to investigate how wind turbine pattern influences capture time. Additional data
	such as which TOC (1 of 3) was appended to our data set.


2. Methods for processing the data: 

      .csv data can be analysed in R.
	  
	  Games can be remade using MATLAB


3. Instrument- or software-specific information needed to interpret the data: 

	- R version 4.3.2+ [https://www.r-project.org/]


4. Standards and calibration information, if appropriate: 

	- Calibrite ColorChecker Studio (XRITE, Grand Rapids, MI, USA).

5. Environmental/experimental conditions: 

	- Temperature controlled rooms


# 4) DATA-SPECIFIC INFORMATION FOR: 
---------------------------------------------------------------------------------------------------------------------------------------------------------
	(In Alphabetical order)	


## WT_Data_Exp.csv
Wind Turbine Data from Blade Comparison Experiment

1. Number of variables: 45

2. Number of cases/rows: 866

3. Variable List: 

	[1] BirdID. The unique ID for thje bird

	[2] TOC. The TOC the bird was assigned to.

	[3] TrialType. Always 5_ComparativeExperiment.

	[4] trainWindImage. The wind turbine pattern the bird was trained with (white, red, black, bio).

	[5] shownWindImage. The wind turbine pattern the bird was shown (white, red, black, bio).

	[6] Speed. The wind turbine speed, always 20 rpm.

	[7] number. The trial number, e.g. 1 is trial 1, 1.5 is the second part of trial 1 (one dot only).

	[8] number2. The continuous trial number where the second part of each trial is considered its own trial.

	[9] group. Which block of 4 trials does it belong to (0,1,2,3,4), remember each blade was shown 5 times, once per block.

	[10] Date. the date for when the trial took place.

	[11] Time. the time the trial took place.

	[12] FirstPeck. was the peck first or second.

	[13] TimeToPeck. the time taken for the bird to peck the dot in seconds, NA indicates no peck.

	[14] Clicks. stores coordinates for where the bird clicked (note most are NAs as we switch to manually recording clicks with key presses).

	[15] SuccFail. did the bird successfully peck (succ) or did it fail (fail)

	[16] DotType. was the dot pecked an inner dot or an outer dot.

	[17] leftRight_i. was the inner dot on the left or the right?

	[18] leftRight_o. was the outer dot on the left or the right?

	[19] TargetX_i. the x coordinate of the inner dot.

	[20] TargetX_o. the x coordinate of the outer dot.

	[21] TargetY_i. the y coordinate of the inner dot.

	[22] TargetY_o. the y coordinate of the outer dot.

	[23] TargetSize. the target size in pixels.

	[24] TargetAngle_i. the angle of the inner dot from the centre.

	[25] TargetAngle_o. the angle of the outer dot from the centre.

	[26] RadiusCentre_i. the radius of the inner dot from the centre (always 512px).

	[27] RadiusCentre_o. the radius of the outer dot from the centre (always 731px).

	[28] TurbineAngle. the angle of the turbine image at the time the dot was pecked.

	[29] Blade1Angle. the angle of blade 1 at the time the dot was pecked.

	[30] Blade2Angle. the angle of blade 2 at the time the dot was pecked.

	[31] Blade3Angle. the angle of blade 3 at the time the dot was pecked.

	[32] dotCol. the colour of the dot in sRGB (0-1).

	[33] timeOutTime. the time limit for the experiment.

	[34] frames. the number of frames shown.
	
	[35] screenWidth. the width of the screen in pixels.

	[36] screenHeight. the height of the screen in pixels.
	
	[37] TargetPeckedX. the x coordinate of the target that was pecked.

	[38] TargetPeckedY. the y coordinate of the target that was pecked.

	[39] TaretPeckedAngle. the angle of the target that was pecked.

	[40] TargetPeckedLR. was the target pecked on the left or on the right?

	[41] NovelBlade. was the blade pattern shown the one it was trained with (trained) or novel (novel).

	[42] TratmentOrder. duplicate of group but from 1-5.

	[43] Weird. binary was there anything off about the data?

	[44] Exlucde. binary, should the data be excluded.

	[45] Comments. why should the data be excluded if at all.


## WT_Data_TOC.csv
Wind Turbine Data from Blade Comparison Experiment

1. Number of variables: 2

2. Number of cases/rows: 22

3. Variable List: 

	[1] BirdID. The unique ID for thje bird

	[2] TOC. The TOC the bird was assigned to.


## WT_Data_Training.csv
Wind Turbine Data from Blade Comparison Experiment

1. Number of variables: 45

2. Number of cases/rows: 1062

3. Variable List: 

	[1] BirdID. The unique ID for thje bird

	[2] TOC. The TOC the bird was assigned to.

	[3] TrialType. The speed setting (0_Static, 1_Slow, 2_Medium, 3_Fast).

	[4] trainWindImage. The wind turbine pattern the bird was trained with (white, red, black, bio).

	[5] Speed. The wind turbine speed, ( 0 , 2.5 , 5 , 10 rpm).

	[6] Number. The trial number for the speed.

	[7] Date. the date for when the trial took place.

	[8] Time. the time the trial took place.

	[9] InOut. was the dot shown in (inner) or out (outer).

	[10] leftRight. was the dot shown on the left or the right.

	[11] TimeToPeck. the time taken for the bird to peck the dot in seconds, NA indicates no peck.

	[12] SuccFail. did the bird successfully peck (succ) or did it fail (fail)

	[13] TargetX. the x coordinate of the target.

	[14] TargetY. the y coordinate of the target.
	
	[15] TargetSize. the target size in pixels.
	
	[16] TargetAngle. the angle of the target from the centre.

	[17] RadiusCentre. the radius of the target from the centre (512 = inner, 731 = outer).

	[18] TurbineAngle. the angle of the turbine image at the time the dot was pecked.

	[19] Blade1Angle. the angle of blade 1 at the time the dot was pecked.

	[20] Blade2Angle. the angle of blade 2 at the time the dot was pecked.

	[21] Blade3Angle. the angle of blade 3 at the time the dot was pecked.

	[22] dotCol. the colour of the dot in sRGB (0-1).

	[23] timeOutTime. the time limit for the experiment.

	[24] frames. the number of frames shown.

	[25] screenWidth. the width of the screen in pixels.

	[26] screenHeight. the height of the screen in pixels.
