## Supplementary material for "Biologically inspired warning patterns deter birds from wind turbines": README_file.txt

This README.txt file was generated on 2025.04.02 [yyy,mm,dd] by George RA Hancock

1) GENERAL INFORMATION

---------------------------------------------------------------------------------------------------------------------------------------------------------

1. Title of Dataset: Biologically inspired warning patterns deter birds from wind turbines.

2. Author Information
	A. Principal Investigator Contact Information
		Name: George Hancock
		Institution: University of Exeter / University of Helsinki
	

3. Date of data collection. January 2024 - April 2024 

4. Data collection location: Lammi Biological Research Station, Pääjärventie 320, 16900 Lammi, Finland

5. Funding Body: Research Council of Finland and The Finnish Society of Sciences and Letters

6. Software recommendations. 

	Statistical Analyses

	- R version 4.3.2+ [https://www.r-project.org/]
	

	Wind Turbine Bird Game
	(NB, MATLAB requires a software lisence, unfortunatley our code is not compatible with Octave).

	- MATLAB [https://www.mathworks.com/products/matlab.html]

	- Psychtoolbox-3 [https://psychtoolbox.org/]
	


2) DATA & FILE OVERVIEW

---------------------------------------------------------------------------------------------------------------------------------------------------------


1. File List: 

   o. /Datasets/
	(In Alphabetical order)	

	# BG_Geometry.txt
	
	- Contains the mean and standard deviation of Z (depth) for an area x15 the diameter of the target (30mm).
	- Additionally the dataset includes information necessary to merge with other datasets and the number for the original scan.
	- In instances where the scan failed the values are replaced by the average for the habitat it belonged to.

	# BG_Info.txt
	
	- Contains useful information for each backgorund photograph, most of which was unused within the final manuscript.
		 Information about the shadows (are cast shadows visible, is the target entirely in shadow).
		 Information about the backgorund, what is the dominant substrate, are certain features present e.g. stones, is the habitat
			anthropogenic?

	# BG_SiteInfo.txt
	
	- Contains useful information for the photography session of each site.
		 Year,month,day the photographs were taken.
		 What was the location UK county, lat, long.
		 What was the sun elevation and sun azimuth for that time of year.	
	- Some of this data is included in Site Info Table in the supplementary material.


	# BG_Vis_Energy.txt
	
	- Contains the gabor output metrics for the background or an area x15 the diameter of the target (30mm).


	# TG_Vis_Energy.txt
	
	- Contains the gabor output metrics for the target (30mm).


   o. /Sample_Scene_Data/

	# /Adjusted Scans/

	- Each folder contains a example scan for scene 1 of the habitat. 

	- Depth Map.tif a 2D 32-bit map where the pixel value indicates the height (Z value), can be opened in ImageJ and previewed in 3D 
	  with /plugins/3D/Interactive 3D Surface Plot.

	- fix_cloud.ply, the adjusted .ply file containing X,Y, and Z coordinates. Can be opened in meshlab or a similar ply model viewer.
	 		 Can also be opened in imageJ using the supplementary Import Normalised PLY plugin.

	# /Calibrated Photos/

	- Each folder contains example habitat photos for a diffuse background (D_backgorund) and natural/direct lighting background (N_background). 

	- Backgrounds are saved as .tifs with the ROIs stored.

	- ROIs can be loaded in ImageJ by opening the image and executing: run("To ROI Manager");


   o. /Supplementary_ImageJ_Plugins/
	
	# /3D_Light_Scene_Analysis_Scripts/

	- Contains the batch gabor measurement scripts. These can be run on the sample calibrated photos as a demo.
		To do so run the script and select the /Calibrated_Photos/ folder when prompted to.
	
	- Step 1, run the Batch_ scripts by running them in ImageJ and selecting the folder /Sample_Scene_Data/

	- Step 2, run the _Combine_ scripts by running them in ImageJ and selecting the folder /Sample_Scene_Data/

	NB1. ignore the scripts in LightCamoProject these are for a seperate paper that also uses the backgrounds from this paper.

	NB2. for a more accessible version of these scripts that operate for the micaToolbox please see:

			https://github.com/GeorgeHancock471/Batch_RNL_Chromaticity.git


	# /3D_Measures/

	- Import_Normalised_PLY.txt, allows the user to select a ply that has been standardised in meshlab into ImageJ.

	- PLY Labeller, allows the user to select a ply and add the rois and adjust the height values to be relative to the selected target.

	# /Spatial_Analyses/

	- Includes Jolyon Troscianko's Difference of Gaussians (DoG) and Gabor filter spatial analysis scripts.


   o. RCode_Habitat_&_Light.R
     
        Contains the necessary R code for recreating the plots and statistical analyses used in the paper.


2. Relationship between files, if important: 


   o. /Sample_Scene_Data/Adjusted Scans/
       	To run the R code, it must be in a folder that contains /Datasets/ with all of the stored .txt files
	This is because the R code automatically detects the directory of the R code and sets it as the working directory.


   o. /Supplementary_ImageJ_Plugins/
       	To install copy and paste the content of the plugins folder into the plugins folder of ImageJ v1.52 with MICA installed.


   o. RCode_Habitat_&_Light and Datasets

       	To run the R code, it must be in a folder that contains /Datasets/ with all of the stored .txt files
	This is because the R code automatically detects the directory of the R code and sets it as the working directory.


3. Additional related data collected that was not included in the current data package: N/A

4. Are there multiple versions of the dataset? NO
	A. If yes, name of file(s) that was updated: N/A
		i. Why was the file updated? N/A
		ii. When was the file updated? N/A


3) METHODOLOGICAL INFORMATION

---------------------------------------------------------------------------------------------------------------------------------------------------------

1. Description of methods used for collection/generation of data: 

   o. Photography

	Photographs of 28 different habitats were taken from a ~1.2m using an ASUS A002 smartphone calibrated with imageJ using a 8% reflectance standard. 
	Direct lighting photos were taken on sunny days with <10% cloud cover and the sun unobstructed by passing cloud. Diffuse lighting photos were taken
	from the same location and on the same day using a Neewer 1.5m^3 diffuser tent placed over the top. 

        Uncalibrated copies of the photos are available on request.


   o. 3D Scans

	Scans were taken using the Matterport Scenes app for Android IOS with Tango enabled google depth sensors (a feature of the ASUS A002 smartphone). Each
	scan was taken from the same location as the scene photograph. As the scans worked better under diffuse lighting, but the app could not be triggered via
	bluetooth the photography tent was folded and used as a single diffuser screen when taking the scans. 3D scans were standardised in MeshLab v.2022.02 and 
	exported as .ply files containing only the X,Y, and Z coordinates. .ply files were loaded in ImageJ with a custom script to render an X,Y, Z image where 
	1px equated to 1mm. Blank z values were replaced locally using nearest-neighbour filling, and the grey-targets were labelled using ROIs. 


   o. Analysis Measures.
     
      	For each scene the the photographs and height maps were rescaled to the minimum number of pixels per mm. The photographs were then converted into the
	human CIE L*a*b* colour space. For the photographs and the scans the mean and standard deviation were measured. Then for the pattern scale and orientation
	analyses gabor filters were used to extract the pattern contrast at 8 different spatial scales [0.878mm, 1.758mm, 3.516mm , 7.031mm, 14.062mm, 28.125mm, 
	56.25mm, 112.5mm] and 4 different angles [0°,45°,90°,135°]. For each scale the max energy at a given angle and the energy at the orthogonal angle were
	also reported. 


2. Methods for processing the data: 

      Reconstructing the data used requires the calibrated background photographs and the supplementary ImageJ plugins


3. Instrument- or software-specific information needed to interpret the data: 

	- R version 4.3.2+ [https://www.r-project.org/]


4. Standards and calibration information, if appropriate: 

	- 8% reflectance standard

5. Environmental/experimental conditions: 

	- artificially diffuse lighting conditions, dry with <10% cloud cover


4) DATA-SPECIFIC INFORMATION FOR: 
	(In Alphabetical order)	

4.1) BG_Geometry.txt
---------------------------------------------------------------------------------------------------------------------------------------------------------
Background Geometry Data

1. Number of variables: 6

2. Number of cases/rows: 672

3. Variable List: 

	[1] Habitat. the unique habitat ID e.g. 01_grassWildFlowers or 22_estuaryHabitat
	[2] Unique-Location. the unique code for each scene = Habitat + Scene Number (starts from zero, 0-23)
	[3] Scan-Original. The code number of the original scan (starts from 1, 1-24)
	[4] Unique-Scan. The unique code of the original scan  = Habitat + Scan-Original
	[5] BG_E15_Z_Mean. The Mean Z(height) for an area 15x the diameter of the target.
	[6] BG_E15_Z_Dev. The StdDev of Z(height) for an area 15x the diameter of the target.


4.2) BG_Info.txt
---------------------------------------------------------------------------------------------------------------------------------------------------------
Scene Information

1. Number of variables: 16

2. Number of cases/rows: 1344 (NB double geometry data length due to 2x lighting)

3. Variable List: 

	[1] Habitat. the unique habitat ID e.g. 01_grassWildFlowers or 22_estuaryHabitat.
	[2]	Background.ID. the photograph code (N=natural, D=diffuse), number equals scene number (0-23) 
					e.g. N_background_0.tif = the natural (direct lighting photo) for the first scene.
	[3]	Unique.Background. the unique code for each photograph = Habitat + PhotoName.
	[4]	Unique.Location. the unique code for each scene = Habitat + Scene Number (starts from zero, 0-23).
	[5]	RecievedShadows. does the target have recieved shadows on it 0/1, no/yes (binary) .
	[6]	SmotheredShadow. is the target entirely in shadow 0/1, no/yes (binary).
	[7]	Dappled. is there dappled lighting within the scene 0/1, no/yes (binary).
	[8]	CastShadow. is the cast shadow of the target visible 0/1, no/yes (binary).
	[9]	ExposureIssue. is the image over exposed in dappled regions 0/1, no/yes (binary).
	[10] Anthropogenic. is the scene anthropogenic or does it include manmade objects 0/1, no/yes (binary).
	[11] DominantSubstrate. which substrate makes up the majority of the scene? grass,leaflitter,bare,veg,gravel?
	[12] stones. does the scene have stones 0/1, no/yes (binary).
	[13] leaves. does the scene have leaflitter 0/1, no/yes (binary).
	[14] veg. does the scene have non-grass vegetation 0/1, no/yes (binary).
	[15] grass. does the scene have grass 0/1, no/yes (binary).
	[16] bare. does the scene have bare ground 0/1, no/yes (binary).


4.3) Site_Info.txt
---------------------------------------------------------------------------------------------------------------------------------------------------------
Site Information

1. Number of variables: 12

2. Number of cases/rows: 28 (one for each habitat)

3. Variable List: 

	[1] Habitat. the unique habitat ID e.g. 01_grassWildFlowers or 22_estuaryHabitat.
	[2] Year. the year the photographs were collected.
	[3] Month. the month the photographs were collected.
	[4] Day. the day the photographs were collected.
	[5] Time. the time of day (UK time) that the collection started (collection took ~2 hours)
	[6] TimeUTC. the UTC time.
	[7] Country. the country that the photographs were taken (always UK).
	[8] County. the county the photographs were taken.
	[9] Lat. the latitude of the photography site.
	[10] Long. the longitude of the photography site.
	[11] SunElevation. the elevation of the sun at the start of collection.
	[12] SunAzimuth. the azimuth of the sun at the start of collection.

4.4) BG_Vis_Energy.txt
---------------------------------------------------------------------------------------------------------------------------------------------------------
Background Energy Metrics

1. Number of variables: 196

2. Number of cases/rows: 1344 (two for each background, direct and diffuse)

3. Variable List: 

	[1] Habitat. the unique habitat ID e.g. 01_grassWildFlowers or 22_estuaryHabitat.
	[2]	Background.ID. the photograph code (N=natural, D=diffuse), number equals scene number (0-23) 
					e.g. N_background_0.tif = the natural (direct lighting photo) for the first scene.
	[3]	Unique.Background. the unique code for each photograph = Habitat + PhotoName.
	[4]	Unique.Location. the unique code for each scene = Habitat + Scene Number (starts from zero, 0-23).
	[5-196] Due to the large number of repeats the variables are broken down by their codes. 
			Output here represents the energy output metrics for different scales, orientations and channels.
			
			BG_E15, denotes that it is the background expanded by 15x (all variables are such).
			
			Next is whether it is luminance (L*, [5-68]) , red-green (A, [69-132]) or blue-yellow (B,[133-196])
			
			Next scale S smaller numbers indicate smaller spatial scales (higher frequency)
				[1,2,3,4,5,6,7,8] = [0.878mm, 1.758mm, 3.516mm , 7.031mm, 14.062mm, 28.125mm, 56.25mm, 112.5mm]
				
			Next is Gabor, this simply indicates that these are metrics from the gabor output.
			
			Then lastly is the specific metric these are:
			contrast, the average energy across all angles for that scale.
			deviation, the standard deviation of contrast for the angles at that scale.
			peak, the contrast at the angle with the highest contrast.
			orthogonal, the contrast at the angle orthogonal to the highest contrast.
			D[0,90,45,135], the contrast at the different angles these are used to calculate the directionality and the V (0) - H (90) measures in the R code.


4.5) TG_Vis_Energy.txt
---------------------------------------------------------------------------------------------------------------------------------------------------------
Target Energy Metrics

1. Number of variables: 196

2. Number of cases/rows: 1344 (two for each background, direct and diffuse)

3. Variable List: 

	[1] Habitat. the unique habitat ID e.g. 01_grassWildFlowers or 22_estuaryHabitat.
	[2]	Background.ID. the photograph code (N=natural, D=diffuse), number equals scene number (0-23) 
					e.g. N_background_0.tif = the natural (direct lighting photo) for the first scene.
	[3]	Unique.Background. the unique code for each photograph = Habitat + PhotoName.
	[4]	Unique.Location. the unique code for each scene = Habitat + Scene Number (starts from zero, 0-23).
	[5-196] Due to the large number of repeats the variables are broken down by their codes. 
			Output here represents the energy output metrics for different scales, orientations and channels.
			
			TG_Grey, denotes that the metric is for the grey-target.
			
			Next is whether it is luminance (L*, [5-68]) , red-green (A, [69-132]) or blue-yellow (B,[133-196])
			
			Next scale S smaller numbers indicate smaller spatial scales (higher frequency)
				[1,2,3,4,5,6,7,8] = [0.878mm, 1.758mm, 3.516mm , 7.031mm, 14.062mm, 28.125mm, 56.25mm, 112.5mm]
				
			Next is Gabor, this simply indicates that these are metrics from the gabor output.
			
			Then lastly is the specific metric these are:
			contrast, the average energy across all angles for that scale.
			deviation, the standard deviation of contrast for the angles at that scale.
			peak, the contrast at the angle with the highest contrast.
			orthogonal, the contrast at the angle orthogonal to the highest contrast.
			D[0,90,45,135], the contrast at the different angles these are used to calculate the directionality and the V (0) - H (90) measures in the R code.
