## Supplementary material for "Biologically inspired warning patterns deter birds from wind turbines": WT_R_Code.R

```
#..................................................................
#----Setup----
#..................................................................
#..................................................................


# GRAH's Laptop Only, the usual R directory is blocked, so a custom directory is specified
# Please delete this code before running.

.libPaths()
new_library_path <- "C:\\Users\\Localadmin_hangeorg\\Documents\\RStudio\\library\\"
.libPaths(new_library_path)


# Set path automatically to R location
#   please make sure the datasets are in the same folder as R.
setwd(dirname(rstudioapi::getSourceEditorContext()$path))


#load packages
#   please make sure they are installed.
library(lme4)
library(lmerTest) 
library(performance)
library(emmeans)
library(see)
library(ggplot2) #
library(ggsignif)
library(car) 
library(gtools)
library(coxme)
library(sjPlot)
library(multcomp)
library(scales)


# Themes ----
#.................................


# creates colour palettes used for plots
my_colors <- c("gray", "red", "#474747", "orange")
my_colors2 <- c( "orange",  "#474747","red","gray") #Inverse of above


# IMPORT ----- 
#.................................

##i) Training Data----
#'''''''''''''''''''

  #Loads and adjusts the dataset
  
  #Read the data
  df_train = read.csv('WT_Data_Training.csv')
  head(df_train) # check headings
  nrow(df_train) 
  
  t = read.csv('WT_Data_TOC.csv') # Add in TOC numbers for each bird.
  df_train=merge(t,df_train)
  nrow(df_train) #good, there is the same number of rows!
  
  #Convert trainWindImage into a factor
  df_train$trainWindImage=factor(df_train$trainWindImage)
  
  #Add patterned vs un-patterned variable
  df_train$Patterned = ifelse(df_train$trainWindImage=="White","Un-Patterned","Patterned")
  
  #Set factor order for turbine blades (inverse alphabet)
  turbineOrder = c("White","Red","Black","Bio")
  df_train$trainWindImage <- factor(df_train$trainWindImage, levels = turbineOrder) 
  df_train$trainWindImage=factor(df_train$trainWindImage)

  
  #Create timeout binomial variable.
  df_train$TimeOut=ifelse(df_train$SuccFail=="Succ",0,1)
  df_train$TimeOut=as.factor(df_train$TimeOut)
  

  #Create variable where a timeout is treated as the timelimit rather than NA.
  df_train$TimeToPeck_TimeLimit = ifelse(is.na(df_train$TimeToPeck), df_train$timeOutTime, df_train$TimeToPeck)
  df_train$TimeToPeck_TimeLimit
  
  #Rename InOut to DotType.
  df_train$DotType = df_train$InOut
  

##ii) Experiment Data----
#'''''''''''''''''''

  
  #Loads and adjusts the dataset
  
  #read data
  df = read.csv('WT_Data_Exp.csv')
  
  #remove excluded trials
  df = df[df$Exclude==0,] #this keeps only the rows where exclude is 0
  
  #make TOC a factor (categorical variable) instead of an integer
  df$TOC = factor(df$TOC)
  
  #create a variable that denotes each unique trial for each bird 
  #(so we can account for which in/out dots were presented together)
  df$TrialNumber = floor(df$Number)
  df$TrialID = paste0(df$BirdID, '_', floor(df$Number)) 
  df$TrialID2 = paste0(df$BirdID, '_',(df$Number)) 
  
  #TimeOut Binomial
  df$TimeOut=ifelse(df$SuccFail=="Succ",0,1)
  df$TimeOut=as.factor(df$TimeOut)
  
  #Reverse factor levels
  df$shownWindImage=factor(df$shownWindImage)
  df$shownWindImage <- factor(df$shownWindImage, levels = rev(levels(df$shownWindImage))) 
  
  df$trainWindImage=factor(df$trainWindImage)
  df$trainWindImage <- factor(df$trainWindImage, levels = rev(levels(df$trainWindImage)))
  
  df$TimeToPeck = as.numeric(df$TimeToPeck)
  
  #Create variable where a timeout is treated as the timelimit rather than NA.
  df$TimeToPeck_TimeLimit = ifelse(is.na(df$TimeToPeck), df$timeOutTime, df$TimeToPeck)
  df$TimeToPeck_TimeLimit
  
  # Create Block Number variable
  df$BlockNumber = df$Group
  
  #Patterned
  df$Patterned = ifelse(df$shownWindImage=="White","Un-Patterned","Patterned")
  

# CHECK ----- 
#.................................
  
  # Confirm the number of birds

  # Number of birds
  length(unique(df$BirdID)) #22 birds total
  
  t=subset(df,trainWindImage=="White")
  length(unique(t$BirdID)) # 5 birds were trained with white
  
  t=subset(df,trainWindImage=="Red")
  length(unique(t$BirdID))  # 5 birds were trained with red
  
  t=subset(df,trainWindImage=="Black")
  length(unique(t$BirdID)) # 6 birds were trained with black
  
  t=subset(df,trainWindImage=="Bio")
  length(unique(t$BirdID)) # 6  birds were trained with bio
  

# ANALYSES ----- 
#.................................
  
## Scale Data ----- 
##'''''''''''''''''''        

  ###i) Training Data ----
  df_train$Log_TimeToPeck = log( df_train$TimeToPeck)
  df_train$FactorSpeed = as.factor(df_train$Speed)
  df_train$Log_Number = log(df_train$Number)
  
  df_train_sub = subset(df_train,Number<=10)
  tS = subset(df_train_sub,TimeOut=="1")
  tS
  
  df_train_sub = subset(df_train_sub,TimeToPeck>0.3)
  df_train_sub = rbind(df_train_sub,tS)
  
  df_scaled_train <-df_train_sub %>%
    mutate(across(where(is.numeric), scale))
  
  
  ###ii) Comparative Data ----
  df$Log_TimeToPeck = log( df$TimeToPeck)
  df$Log_TimeToPeck_TimeLimit = log( df$TimeToPeck_TimeLimit)
  df$TrialID
  
  df_scaled <-df %>%
    mutate(across(where(is.numeric), scale))
  
  
  
  
##1) Training Analysis ----- 
##'''''''''''''''''''''''      

  
  ###i) Train Timeouts ----
  ###~~~~~~~~~~~~~~~~~~~~
  
  ####.Percentage of Failures ----
  
    nrow(subset(df_scaled_train,TimeOut=="1"))/nrow(df_scaled_train)*100
    #Only 10%
  
  ####.Base Models ----
  
    test_base_model1 <- glmer(TimeOut ~trainWindImage*FactorSpeed*Number*DotType+(1|BirdID), data=subset(df_scaled_train), family=binomial)
    summary(test_base_model1)
    #Models fail to converge!
    
    
    test_base_model2 <- glmer(TimeOut ~trainWindImage+FactorSpeed+Number+DotType+(1|BirdID), data=subset(df_scaled_train), family=binomial)
    summary(test_base_model2)
    #Model converged!
    
    #Base model is defined as
    base_model <- glmer(TimeOut ~ trainWindImage+FactorSpeed+Number+DotType+(1|BirdID), data=subset(df_scaled_train), family=binomial)
    summary(base_model)
    summary(base_model)
    qqPlot(residuals(base_model))
    hist(residuals(base_model))  
  
    
  ####.Likelihood Ratio Tests ----
    
    mod_null1 <- glmer(TimeOut ~ FactorSpeed+Number+DotType+(1|BirdID), data=subset(df_scaled_train), family=binomial)
    mod_null2 <- glmer(TimeOut ~ trainWindImage+Number+DotType+(1|BirdID), data=subset(df_scaled_train), family=binomial)
    mod_null3 <- glmer(TimeOut ~ trainWindImage+FactorSpeed+DotType+(1|BirdID), data=subset(df_scaled_train), family=binomial)
    mod_null4 <- glmer(TimeOut ~ trainWindImage+FactorSpeed+Number+(1|BirdID), data=subset(df_scaled_train), family=binomial)
    
    print("null_1")
    anova(base_model, mod_null1)  # Train image is not significant
    print("null_2") 
    anova(base_model, mod_null2)  # Speed is significant
    print("null_3") 
    anova(base_model, mod_null3)  # Number is significant
    print("null_4") 
    anova(base_model, mod_null4)  # DotType is not significant
    
    
  ####.Reduced model ----
    
    reduced_model <-  glmer(TimeOut ~FactorSpeed+Number+(1|BirdID), data=subset(df_scaled_train), family=binomial)
    summary(reduced_model)
    qqPlot(residuals(reduced_model))
    hist(residuals(reduced_model))  
    
    # Post-hoc comparisons for FactorSpeed
    emmeans(reduced_model, pairwise ~ FactorSpeed, adjust = "tukey")
    
    

    
    
    
    
  
  ###ii) Train Capture Time ----
  ###~~~~~~~~~~~~~~~~~~~~
  
  ####. Base model----
  
  # Base model is the model with all variables.
  
  # TimeToPeck = Capture time
  base_model <- lmer(Log_TimeToPeck ~trainWindImage+FactorSpeed+Number+DotType+(1|BirdID), data=subset(df_scaled_train ))
  summary(base_model)
  qqPlot(residuals(base_model))
  hist(residuals(base_model))  
  
  
  ####. Likelihood ratio test ----
  
  #Single Variable deletion
  mod_null1 <- lmer(Log_TimeToPeck ~FactorSpeed+Number+DotType+(1|BirdID), data=subset(df_scaled_train))
  mod_null2 <- lmer(Log_TimeToPeck ~trainWindImage+Number+DotType+(1|BirdID), data=subset(df_scaled_train))
  mod_null3 <- lmer(Log_TimeToPeck ~trainWindImage+FactorSpeed+DotType+(1|BirdID), data=subset(df_scaled_train))
  mod_null4 <- lmer(Log_TimeToPeck ~trainWindImage+FactorSpeed+Number+(1|BirdID), data=subset(df_scaled_train))
  
  print("null_1")
  anova(base_model, mod_null1)  # Wind turbine type is not significant
  print("null_2") 
  anova(base_model, mod_null2)  # Speed is significant
  print("null_3") 
  anova(base_model, mod_null3)  # Trial Number is significant
  print("null_4") 
  anova(base_model, mod_null4)  # DotType is signficiant!
  
  
  ####. Reduced model ----
  
  reduced_model <- lmer(Log_TimeToPeck ~FactorSpeed+DotType+Number+(1|BirdID), data=subset(df_scaled_train ))
  summary(reduced_model)
  qqPlot(residuals(reduced_model))
  hist(residuals(reduced_model))  
  
  emmeans(reduced_model, pairwise ~ FactorSpeed, adjust = "tukey")
  

  ####. Figure 2 A ----
  
  p <- ggplot(subset(df_train_sub ), aes(x=Number, y=TimeToPeck, col=DotType)) +
    geom_smooth (aes(),method=glm,formula=y~poly(x,1),alpha=0.15,size=0.75)+
    facet_grid(. ~ as.factor(Speed))+
    # Labels
    labs(title = "",
         x = "Trial Number",
         y = "Log Time to Peck",
         color = "Blade Type") # Specify the color palette
  
  p + theme_classic() +
    theme(legend.position = "none",  # Hide legend
          text = element_text(size = 12),
          line = element_line(size=0.6), 
          axis.title.y = element_text(margin = margin(t = 0, r = 10, b = 0, l = 0)),  # Adjust y-axis title margin
          axis.title.x = element_text(margin = margin(t = 10, r = 0, b = 0, l = 0)))  # Adjust x-axis title margin
  
  
      #UNUSED FIGURES
            
            # Create the plot
            p <- ggplot(df_train_sub, aes(x = FactorSpeed, y = TimeToPeck, 
                                 group = FactorSpeed)) +
              
              # Stats
              stat_summary(fun.data = mean_se, geom = "errorbar", aes(), 
                           width = 0.3,size=1, position = position_dodge(width = 0.3)) +  # Dodge error bars
          
              stat_summary(fun = mean, geom = "point", aes(), size = 4, shape = 21, fill="black", 
                           position = position_dodge(width = 0.3)) +  # Dodge mean points
              
              
              # Labels and plot customization
              labs(title = "",
                   x = "Wind Turbine Speed (rpm)",
                   y = "Time to Peck (Seconds)",
                   color = "Blade Type") +
              
              # Add significance comparisons
              geom_signif(
                comparisons = list(c("2.5", "0"), c("2.5", "5"), c("0", "5"),
                                   c("5", "10"),c("2.5", "10"), c("0", "10")),
                annotations = c("***","***","ns","ns","***","ns"),  # Significance codes
                y_position = c(19,22,25,28,31,34),  # Dynamic y-position
                textsize = 4, 
                vjust = 0,  
                hjust=0.2,
                tip_length = c(0.01, 0.01,0.01,0.01,0.01,0.01),
                map_signif_level = TRUE  # Automatically map significance levels to symbols
              ) +
              
              # Custom color scales
              scale_alpha_manual(values = c(1, 0.25)) + 
              scale_fill_manual(values = my_colors) + 
              scale_color_manual(values = my_colors) +
              
              # Adjust the y-axis to add more space between the bars
              scale_y_continuous(expand = expansion(mult = c(0.05, 0.2))) +  # More space at the top
              
              theme_classic() +
              theme(legend.position = "none",  # Hide legend
                    text = element_text(size = 12),
                    line = element_line(size = 0.6), 
                    axis.title.y = element_text(margin = margin(t = 0, r = 10, b = 0, l = 0)),
                    axis.title.x = element_text(margin = margin(t = 10, r = 0, b = 0, l = 0))) 
            
            # Display the plot
            print(p)
            
            
  
  
  
  
  
  
  
  
  
  # Create the plot
  p <- ggplot(df_train_sub, aes(x = DotType, y = TimeToPeck 
                                )) +
    
    # Stats
    stat_summary(fun.data = mean_se, geom = "errorbar", aes(color = DotType, fill=DotType), 
                 width = 0.3,size=1, position = position_dodge(width = 0.3)) +  # Dodge error bars
    
    stat_summary(fun = mean, geom = "point", aes(color = DotType, fill=DotType), size = 4, shape = 21, 
                 position = position_dodge(width = 0.3)) +  # Dodge mean points
    
    
    # Labels and plot customization
    labs(title = "",
         x = "Dot Type (in/out)",
         y = "Time to Peck (Seconds)",
         color = "Blade Type") +
    
    # Add significance comparisons
    geom_signif(
      comparisons = list(c("in", "out")),
      annotations = c("**"),  # Significance codes
      y_position = c(17),  # Dynamic y-position
      textsize = 4, 
      vjust = 0,  
      hjust=0.2,
      tip_length = c(0.01),
      map_signif_level = TRUE  # Automatically map significance levels to symbols
    ) +
    
    # Custom color scales
    scale_alpha_manual(values = c(1, 0.25)) + 
    # Adjust the y-axis to add more space between the bars
    scale_y_continuous(expand = expansion(mult = c(0.05, 0.2))) +  # More space at the top
    
    theme_classic() +
    theme(legend.position = "none",  # Hide legend
          text = element_text(size = 12),
          line = element_line(size = 0.6), 
          axis.title.y = element_text(margin = margin(t = 0, r = 10, b = 0, l = 0)),
          axis.title.x = element_text(margin = margin(t = 10, r = 0, b = 0, l = 0))) 
  
  # Display the plot
  print(p)
  
  
  

  
  ##2) Comparative Exp Analysis ----- 
  ##'''''''''''''''''''''''''''''''''''      
  
  ###i) Comp Timeouts ----
  ###~~~~~~~~~~~~~~~~~~~~
  
  ####. Percentage Timeouts----
  
  
  #WHITE BLADES
  T = subset(df_scaled,shownWindImage=="White")
  nrow(subset(T,TimeOut=="1" ))/nrow(T)*100
  #4.19
  
  #RED BLADES
  T = subset(df_scaled,shownWindImage=="Red")
  nrow(subset(T,TimeOut=="1" ))/nrow(T)*100
  #13.42
  
  #BLACK BLADES
  T = subset(df_scaled,shownWindImage=="Black")
  nrow(subset(T,TimeOut=="1" ))/nrow(T)*100
  #10.45
  
  #BIO BLADES
  T = subset(df_scaled,shownWindImage=="Bio")
  nrow(subset(T,TimeOut=="1" ))/nrow(T)*100
  #39.53488
  
  
    ####. Base models ----
  
    # complex model failure
      test_mod1 <- glmer(TimeOut ~shownWindImage*NovelBlade*TrialNumber+(1|BirdID)+(1|TrialID), data=subset(df_scaled), family=binomial)
      summary(test_mod1)
      # Model failed to converge!
    
    # Simplify by removing interactions.
      test_mod2 <- glmer(TimeOut ~shownWindImage+NovelBlade+TrialNumber+(1|BirdID)+(1|TrialID), data=subset(df_scaled), family=binomial)
      summary(test_mod2)
      # Model successfully converged!
  
    
    # define the base model
      base_model <- glmer(TimeOut ~shownWindImage+NovelBlade+TrialNumber+(1|BirdID)+(1|TrialID), data=subset(df_scaled), family=binomial)
      summary(base_model)
      qqPlot(residuals(base_model))
      hist(residuals(base_model))  
  
    ####. Likelihood ratio test ----
    
    #Single Variable deletion
    mod_null1 <- glmer(TimeOut ~NovelBlade+TrialNumber+(1|BirdID)+(1|TrialID), data=subset(df_scaled), family=binomial)
    mod_null2 <- glmer(TimeOut ~shownWindImage+TrialNumber+(1|BirdID)+(1|TrialID), data=subset(df_scaled), family=binomial)
    mod_null3 <- glmer(TimeOut ~shownWindImage+NovelBlade+(1|BirdID)+(1|TrialID), data=subset(df_scaled), family=binomial)
    
    print("null_1")
    anova(base_model, mod_null1)  # Wind turbine image is significant
    print("null_2") 
    anova(base_model, mod_null2)  # Novelty is significant
    print("null_3") 
    anova(base_model, mod_null3)  # Block number is significant
    
    ####. Reduced model ----
    
    # Final model is the original base model
    
    reduced_model <- glmer(TimeOut ~shownWindImage+NovelBlade+TrialNumber+(1|BirdID)+(1|TrialID), data=subset(df_scaled), family=binomial)
    summary(reduced_model)
    qqPlot(residuals(reduced_model))
    hist(residuals(reduced_model))  

    # Post-hoc comparisons for FactorSpeed
    emmeans(reduced_model, pairwise ~ shownWindImage, adjust = "tukey")
  

    ####. Figure 3 A ----
    
    # Calculate centre dashed line for white
    sdf= subset(df,shownWindImage=="White")
    white_avg = mean(as.numeric(sdf$TimeOut))-1
    white_avg
    
    # Create the plot
    p <- ggplot(subset(df), aes(x = shownWindImage, y = as.numeric(TimeOut)-1, 
                                fill = shownWindImage, group = shownWindImage)) +
      geom_hline(yintercept = white_avg, col = "gray", linetype = 2) +
      stat_summary(fun.data = mean_se, geom = "errorbar", aes(col = shownWindImage), 
                   width = 0.3,size=1, position = position_dodge(width = 0.3)) +  
      stat_summary(fun = mean, geom = "point", size = 4, shape = 21, aes(col = shownWindImage), 
                   fill = "white", position = position_dodge(width = 0.3)) +  
      stat_summary(fun = mean, geom = "point", aes(col = shownWindImage), size = 4, shape = 21, 
                   position = position_dodge(width = 0.3)) +  # Dodge mean points
    
      # Plot Labels
      labs(title = "",
           x = "Wind Turbine Pattern",
           y = "Likelihood of a TimeOut",
           color = "Blade Type") +
      
      
      # Significance Comparisons
      geom_signif(
        comparisons = list(c("Red", "White"), c("Red", "Black"), c("White", "Black"),
                           c("Black", "Bio"),c("Red", "Bio"), c("White", "Bio")),
        annotations = c("ns","ns","ns","***","***","***"),  # Significance codes
        y_position = c(0.15,0.20,0.28,0.45,0.55,0.65),  # Dynamic y-position
        textsize = 4, 
        vjust = 0,  
        hjust=0.2,
        tip_length = c(0.015,0.015,0.035,0.015,0.035,0.055),
        map_signif_level = TRUE  # Automatically map significance levels to symbols
      ) +
      
      # Custom color scales
      scale_alpha_manual(values = c(1, 0.25)) + 
      scale_fill_manual(values = my_colors) + 
      scale_color_manual(values = my_colors) +
      
      scale_y_continuous(expand = expansion(mult = c(0.05, 0.2))) +  # More space at the top
      theme_classic() +
      theme(legend.position = "none",  # Hide legend
            text = element_text(size = 12),
            line = element_line(size = 0.6), 
            axis.title.y = element_text(margin = margin(t = 0, r = 10, b = 0, l = 0)),
            axis.title.x = element_text(margin = margin(t = 10, r = 0, b = 0, l = 0))) 
    
    # Display the plot
    print(p)
    
  
  
  
  ###ii) Comp Capture Time ----
  ###~~~~~~~~~~~~~~~~~~~~
    
    ####. Average Capture Time----
    
      #WHITE BLADES
      T = subset(df,shownWindImage=="White")
      mean(as.numeric(T$TimeToPeck), na.rm = TRUE)
      #8.36 seconds
      
      #RED BLADES
      T = subset(df,shownWindImage=="Red")
      mean(as.numeric(T$TimeToPeck), na.rm = TRUE)
      #12.05 seconds
      
      #BLACK BLADES
      T = subset(df,shownWindImage=="Black")
      mean(as.numeric(T$TimeToPeck), na.rm = TRUE)
      #14.21 seconds
      
      #BIO BLADES
      T = subset(df,shownWindImage=="Bio")
      mean(as.numeric(T$TimeToPeck), na.rm = TRUE)
      #19.41
      
    
    ####. Base models ----
    
      # TimeToPeck = Capture time
      base_model <- lmer(Log_TimeToPeck ~shownWindImage*NovelBlade*TrialNumber*DotType+FirstPeck+(1|BirdID)+(1|TrialID), data=subset(df_scaled))
      summary(base_model)
      qqPlot(residuals(base_model))
      
      ####. Likelihood ratio test ----
      
      #Single Variable deletion
      mod_null1 <- lmer(Log_TimeToPeck ~ NovelBlade*TrialNumber*DotType+FirstPeck + (1|BirdID) + (1|TrialID), data=subset(df_scaled))
      mod_null2 <- lmer(Log_TimeToPeck ~ shownWindImage*TrialNumber*DotType+FirstPeck + (1|BirdID) + (1|TrialID), data=subset(df_scaled))
      mod_null3 <- lmer(Log_TimeToPeck ~ shownWindImage*NovelBlade*DotType+FirstPeck + (1|BirdID) + (1|TrialID), data=subset(df_scaled))
      mod_null4 <- lmer(Log_TimeToPeck ~ shownWindImage*NovelBlade*TrialNumber+FirstPeck + (1|BirdID) + (1|TrialID), data=subset(df_scaled))
      mod_null5 <- lmer(Log_TimeToPeck ~ shownWindImage*NovelBlade*TrialNumber*DotType + (1|BirdID) + (1|TrialID), data=subset(df_scaled))
      
      print("null_1")
      anova(base_model, mod_null1)  # Shown Turbine image is signficiant
      print("null_2") 
      anova(base_model, mod_null2)  # Novelblade is significant
      print("null_3") 
      anova(base_model, mod_null3)  # TrialNumber is signficiant
      print("null_4") 
      anova(base_model, mod_null4)  # DotType is not signficiant
      print("null_5") 
      anova(base_model, mod_null5)  # FirstPeck is not signficiant
      
    
      
      # Model 1: Only main effects
      mod1 <- lmer(Log_TimeToPeck ~ shownWindImage + NovelBlade + TrialNumber + (1|BirdID) + (1|TrialID), 
                   data=df_scaled)
      
      # Model 2: One two-way interaction at a time
      mod2a <- lmer(Log_TimeToPeck ~ shownWindImage * NovelBlade + TrialNumber + (1|BirdID) + (1|TrialID), 
                    data=df_scaled)
      
      mod2b <- lmer(Log_TimeToPeck ~ shownWindImage * TrialNumber + NovelBlade + (1|BirdID) + (1|TrialID), 
                    data=df_scaled)
      
      mod2c <- lmer(Log_TimeToPeck ~ NovelBlade * TrialNumber + shownWindImage + (1|BirdID) + (1|TrialID), 
                    data=df_scaled)
      
      # Model 3: Two two-way interactions at a time
      mod3a <- lmer(Log_TimeToPeck ~ shownWindImage * NovelBlade + shownWindImage * TrialNumber + (1|BirdID) + (1|TrialID), 
                    data=df_scaled)
      
      mod3b <- lmer(Log_TimeToPeck ~ shownWindImage * NovelBlade + NovelBlade * TrialNumber + (1|BirdID) + (1|TrialID), 
                    data=df_scaled)
      
      mod3c <- lmer(Log_TimeToPeck ~ shownWindImage * TrialNumber + NovelBlade * TrialNumber + (1|BirdID) + (1|TrialID), 
                    data=df_scaled)
      
      # Model 4: All two-way interactions
      mod4 <- lmer(Log_TimeToPeck ~ shownWindImage * NovelBlade + shownWindImage * TrialNumber + NovelBlade * TrialNumber + 
                     (1|BirdID) + (1|TrialID), data=df_scaled)
      
      # Model 5: Three-way interaction
      mod5 <- lmer(Log_TimeToPeck ~ shownWindImage * NovelBlade * TrialNumber + (1|BirdID) + (1|TrialID), 
                   data=df_scaled)
      
      # Likelihood ratio
      anova(mod1, mod2a)  # Does shownWindImage:NovelBlade help?
      anova(mod1, mod2b)  # Does shownWindImage:TrialNumber help?
      anova(mod1, mod2c)  # Does NovelBlade:TrialNumber help?
      
      anova(mod2a, mod3a) # Does adding shownWindImage:TrialNumber to mod2a help?
      anova(mod2b, mod3b) # Does adding NovelBlade:TrialNumber to mod2b help?
      anova(mod2c, mod3c) # Does adding shownWindImage:NovelBlade to mod2c help?
      
      anova(mod4, mod5)   # Does adding three-way interaction help?
      
      
      #mod 1 is the best
    
    ####. Reduced model ----
    
      reduced_model <- lmer(Log_TimeToPeck ~shownWindImage+NovelBlade+TrialNumber+(1|BirdID)+(1|TrialID), data=subset(df_scaled))
      summary(reduced_model)
      qqPlot(residuals(reduced_model))
      hist(residuals(reduced_model))  
    
      # Post-hoc comparisons for shownWindImage
      emmeans(reduced_model, pairwise ~ shownWindImage, adjust = "tukey")

      
    
    ####. Figure 3 B ----
    
      # calculate averages for white
      sdf_scaled= subset(df,shownWindImage=="White" & as.numeric(TimeToPeck)>0)
      white_avg = mean(sdf_scaled$TimeToPeck)
      white_avg
      
      
      # Create the plot
      p <- ggplot(df, aes(x = shownWindImage, y = TimeToPeck, 
                          fill = shownWindImage, group = shownWindImage)) +
        
        #horizontal line
        geom_hline(yintercept = white_avg, col = "gray", linetype = 2) +
        
        # Stats
        stat_summary(fun.data = mean_se, geom = "errorbar", aes(col = shownWindImage), 
                     width = 0.3,size=1, position = position_dodge(width = 0.3)) +  # Dodge error bars
        
        stat_summary(fun = mean, geom = "point", size = 4, shape = 21, aes(col = shownWindImage), 
                     fill = "white", position = position_dodge(width = 0.3)) +  # Dodge mean points
        
        stat_summary(fun = mean, geom = "point", aes(col = shownWindImage), size = 4, shape = 21, 
                     position = position_dodge(width = 0.3)) +  # Dodge mean points
        
        
        # Labels and plot customization
        labs(title = "",
             x = "Wind Turbine Pattern",
             y = "Time to Peck (Seconds)",
             color = "Blade Type") +
        
        # Add significance comparisons
        geom_signif(
          comparisons = list(c("Red", "White"), c("Red", "Black"), c("White", "Black"),
                             c("Black", "Bio"),c("Red", "Bio"), c("White", "Bio")),
          annotations = c("*","ns","***","***","***","***"),  # Significance codes
          y_position = c(10,12,15,17.5,19.5,21.5),  # Dynamic y-position
          textsize = 4, 
          vjust = 0,  
          hjust=0.2,
          #col= c("black","gray","black","black","black","black"),
          tip_length = c(0.01, 0.01,0.015,0.01,0.01,0.01),
          map_signif_level = TRUE  # Automatically map significance levels to symbols
        ) +
        
        # Custom color scales
        scale_alpha_manual(values = c(1, 0.25)) + 
        scale_fill_manual(values = my_colors) + 
        scale_color_manual(values = my_colors) +
        
        # Adjust the y-axis to add more space between the bars
        scale_y_continuous(expand = expansion(mult = c(0.05, 0.2))) +  # More space at the top
        
        theme_classic() +
        theme(legend.position = "none",  # Hide legend
              text = element_text(size = 12),
              line = element_line(size = 0.6), 
              axis.title.y = element_text(margin = margin(t = 0, r = 10, b = 0, l = 0)),
              axis.title.x = element_text(margin = margin(t = 10, r = 0, b = 0, l = 0))) 
      
      # Display the plot
      print(p)
      
      
      
      
    
  
  ###iii) Comp Dot Choice ----
  ###~~~~~~~~~~~~~~~~~~~~
      
    ####. Create Dot Type Subset ----
      
      dat_DotType = df_scaled
      dat_DotType = subset(dat_DotType,FirstPeck=="first")
      dat_DotType = subset(dat_DotType,DotType!="NA")
      dat_DotType$BinaryType = ifelse(dat_DotType$DotType =="in",0,1)
      
    ####. Percentage Outer Chosen First ----
      
      
      #White BLADES
      T = subset(dat_DotType,shownWindImage=="White")
      mean(as.numeric(T$BinaryType), na.rm = TRUE)*100
      #60.19% 
      
      
      #Red BLADES
      T = subset(dat_DotType,shownWindImage=="Red")
      mean(as.numeric(T$BinaryType), na.rm = TRUE)*100
      #70.10% 
      
      
      #BLACK BLADES
      T = subset(dat_DotType,shownWindImage=="Black")
      mean(as.numeric(T$BinaryType), na.rm = TRUE)*100
      #81.73% 
      
      
      #BIO BLADES
      T = subset(dat_DotType,shownWindImage=="Bio")
      mean(as.numeric(T$BinaryType), na.rm = TRUE)*100
      #81.08% 
      
      # Majority of the time the outer dot is chosen first.
      
    
    ####. Base models ----
      
      
      # Similar model to previous comparison.
      test_base_model1 <- glmer(BinaryType ~ shownWindImage*NovelBlade*TrialNumber+(1|BirdID)+(1|TrialID), 
                         data = dat_DotType, 
                         family = binomial)
      summary(test_base_model1)
      # Model failed to converge!
      
      # Interactions removed.
      test_base_model2 <- glmer(BinaryType ~ shownWindImage+NovelBlade+TrialNumber+(1|BirdID)+(1|TrialID), 
                         data = dat_DotType, 
                         family = binomial)
      summary(test_base_model2)
      # Model failed to converge!
      
      
      # TrialNumber Dropped.
      test_base_model3 <- glmer(BinaryType ~ shownWindImage+NovelBlade+(1|BirdID)+(1|TrialID), 
                         data = dat_DotType, 
                         family = binomial)
      summary(test_base_model3)
      # Model Converged! NovelBlade is not significant
      
      
      # NovelBlade Dropped.
      test_base_model3 <- glmer(BinaryType ~ shownWindImage+TrialNumber+(1|BirdID)+(1|TrialID), 
                         data = dat_DotType, 
                         family = binomial)
      summary(test_base_model3)
      # Model Converged! TrialNumber is not significant
      
    
    ####. Reduced model ----
      
      reduced_model <- glmer(BinaryType ~ shownWindImage+(1|BirdID)+(1|TrialID), 
                             data = dat_DotType, 
                             family = binomial)
      summary(reduced_model )
      
    
      # Post-hoc comparisons for turbine image
      emmeans( reduced_model, pairwise ~ shownWindImage, adjust = "tukey")
      
      
      
      
    
    ####. Figure 3 C ----    
    
        # Get average for white
        sdf= subset(dat_DotType,shownWindImage=="White")
        white_avg = mean(as.numeric(sdf$BinaryType))
        white_avg
        
        
        # Create the plot
        p <- ggplot(dat_DotType, aes(x = shownWindImage, y = as.numeric(BinaryType), 
                                     fill = shownWindImage, group = shownWindImage)) +
          
          # Add horizontal line for average of White
          geom_hline(yintercept = white_avg, col = "gray", linetype = 2) +
          
          # Add horizontal line for liklohood cap
          geom_hline(yintercept = 1, col = "blue", linetype = 3, size=0.5) +
          
          # Add error bars for mean and SE
          stat_summary(fun.data = mean_se, geom = "errorbar", aes(col = shownWindImage), 
                       width = 0.3,size=1, position = position_dodge(width = 0.3)) +  # Dodge error bars
          
          # Add mean points
          stat_summary(fun = mean, geom = "point", size = 4, shape = 21, aes(col = shownWindImage), 
                       fill = "white", position = position_dodge(width = 0.3)) +  # Dodge mean points
          
          stat_summary(fun = mean, geom = "point", aes(col = shownWindImage), size = 4, shape = 21, 
                       position = position_dodge(width = 0.3)) +  # Dodge mean points
          
          
          # Labels and plot customization
          labs(title = "",
               x = "Wind Turbine Pattern",
               y = "Likelihood of a TimeOut",
               color = "Blade Type") +
          
          
          # Add significance comparisons
          geom_signif(
            comparisons = list(c("Red", "White"), c("Red", "Black"), c("White", "Black"),
                               c("Black", "Bio"),c("Red", "Bio"), c("White", "Bio")),
            annotations = c("ns","ns","**","ns","ns","**"),  # Significance codes
            y_position = c(0.75,0.85,0.95,1.05,1.15,1.25),  # Dynamic y-position
            textsize = 4, 
            vjust = 0,  
            hjust=0.2,
            tip_length = c(0.02,0.02,0.03,0.02,0.03,0.04),
            map_signif_level = TRUE  # Automatically map significance levels to symbols
          ) +
          
          
          # Custom color scales
          scale_alpha_manual(values = c(1, 0.25)) + 
          scale_fill_manual(values = my_colors) + 
          scale_color_manual(values = my_colors) +
          
          # Adjust the y-axis to add more space between the bars
          coord_cartesian(ylim = c(0.5, 1.5))+
          
          theme_classic() +
          theme(legend.position = "none",  # Hide legend
                text = element_text(size = 12),
                line = element_line(size = 0.6), 
                axis.title.y = element_text(margin = margin(t = 0, r = 10, b = 0, l = 0)),
                axis.title.x = element_text(margin = margin(t = 10, r = 0, b = 0, l = 0))) 
        
        # Display the plot
        print(p)
        
        
        
        
# SUPPLEMENT ----- 
#.................................    
        
##1) Training Pattern & Peck Time ----- 
##'''''''''''''''''''''''''''''''''''      
#WHITE BLADES
T = subset(df_train_sub, Speed==0 & trainWindImage=="White")
mean(as.numeric(T$TimeToPeck), na.rm = TRUE)
#9.25 seconds

#RED BLADES
T = subset(df_train_sub, Speed==0 & trainWindImage=="Red")
mean(as.numeric(T$TimeToPeck), na.rm = TRUE)
#19.77 seconds

#BLACK BLADES
T = subset(df_train_sub, Speed==0 & trainWindImage=="Black")
mean(as.numeric(T$TimeToPeck), na.rm = TRUE)
#14.75 seconds

#BIO BLADES
T = subset(df_train_sub, Speed==0 & trainWindImage=="Bio")
mean(as.numeric(T$TimeToPeck), na.rm = TRUE)
#17.93 seconds


# In the absence of bird ID as a random effect there is a significant difference from white.
linear_model <- lm(Log_TimeToPeck ~trainWindImage+Number+DotType, data=subset(df_scaled_train, FactorSpeed=="0"))
summary(linear_model)
qqPlot(residuals(linear_model))
hist(residuals(linear_model))   

#Had we measured the initial peck times without a turbine the effect may have been significant.
emmeans(linear_model, pairwise ~ trainWindImage, adjust = "tukey")


  
        
  # Calculate centre dashed line for white
  sdf= subset(df_train_sub, Speed==0 & trainWindImage=="White" & TimeOut==0)
  white_avg = mean(as.numeric(sdf$TimeToPeck))
  white_avg
  
  # Create the plot
  
  p <- ggplot(subset(df_train_sub, TimeOut==0), aes(x=trainWindImage, y=TimeToPeck, col=trainWindImage)) +
    geom_hline(yintercept = white_avg, col = "gray", linetype = 2) + 
    stat_summary(fun = mean, geom = "point", size = 3) + 
    stat_summary(fun.data = mean_se, geom = "errorbar", size = 0.6, alpha=1) + 
    stat_summary(fun = mean, geom = "line", size = 0.8) + 
    #scale_y_log10()+
          scale_color_manual(values = my_colors) +
          scale_fill_manual(values = my_colors) + 
    labs(x="Wind Turbine Pattern",y="Log Time to Peck") +
          facet_grid(. ~ Speed) 
  p + theme_classic()+
    theme(legend.position = "none", text = element_text(size = 14),
          axis.ttle.y = element_text(margin = margin(t = 0, r = 20, b = 0, l = 0)), # Adjust margin for y-axis title
          axis.title.x = element_text(margin = margin(t = 20, r = 0, b = 0, l = 0))) # Adjust margin for x-axis title
  
 
  
  
  # In the absence of bird ID as a random effect there is a significant difference from white.
  base_model <- lm(Log_TimeToPeck ~trainWindImage+Number+DotType, data=subset(df_scaled_train, FactorSpeed=="0"))
  summary(base_model)
  qqPlot(residuals(base_model))
  hist(residuals(base_model))        
  
  
  
  #................................................
  
  
   
  p <- ggplot(subset(df_train_sub, Speed<5 & TimeOut==0), aes(x=trainWindImage, y=TimeToPeck, fill=trainWindImage)) +
    geom_hline(yintercept = white_avg, col = "gray", linetype = 2) + 
    geom_boxplot()+
    #scale_y_log10()+
    scale_color_manual(values = my_colors) +
    scale_fill_manual(values = my_colors) + 
    labs(x="Block Number",y="Time to Peck (Seconds)") +
    facet_grid(. ~ Speed) 
  p + theme_classic()+
    theme(legend.position = "none", text = element_text(size = 14),
          axis.ttle.y = element_text(margin = margin(t = 0, r = 20, b = 0, l = 0)), # Adjust margin for y-axis title
          axis.title.x = element_text(margin = margin(t = 20, r = 0, b = 0, l = 0))) # Adjust margin for x-axis title
  
  
  
        


p = ggplot(subset(df,TimeOut==0), aes(x = BlockNumber, y =TimeToPeck,col=shownWindImage, fill=shownWindImage)) +
  stat_summary(fun = mean, geom = "point", size = 2) + 
  stat_summary(fun.data = mean_se, geom = "errorbar", size = 0.5, alpha=0.2) + 
  stat_summary(fun = mean, geom = "line", size = 0.8) + 
  scale_y_log10()+
  labs(x="Block Number",y="Log Time to Peck") +facet_grid(. ~ as.factor(trainWindImage)) +
  scale_color_manual(values = my_colors) +
  scale_fill_manual(values = my_colors)   # Specify the color palette

p + theme_classic()+
  theme(legend.position = "none", text = element_text(size = 14),
        axis.title.y = element_text(margin = margin(t = 0, r = 20, b = 0, l = 0)), # Adjust margin for y-axis title
        axis.title.x = element_text(margin = margin(t = 20, r = 0, b = 0, l = 0))) # Adjust margin for x-axis title


p = ggplot(subset(df), aes(x = BlockNumber, y =as.numeric(TimeOut)-1,col=shownWindImage, fill=shownWindImage)) +
  stat_summary(fun = mean, geom = "point", size = 2) + 
  stat_summary(fun.data = mean_se, geom = "errorbar", size = 0.5, alpha=0.2) + 
  stat_summary(fun = mean, geom = "line", size = 0.8) + 
  labs(x="Block Number",y="Proportion of Timeouts") +facet_grid(. ~ as.factor(trainWindImage)) +
  scale_color_manual(values = my_colors) +
  scale_fill_manual(values = my_colors)   # Specify the color palette

p + theme_classic()+
  theme(legend.position = "none", text = element_text(size = 14),
        axis.title.y = element_text(margin = margin(t = 0, r = 20, b = 0, l = 0)), # Adjust margin for y-axis title
        axis.title.x = element_text(margin = margin(t = 20, r = 0, b = 0, l = 0))) # Adjust margin for x-axis title


df$FirstPeck

p = ggplot(subset(df, FirstPeck != "NA"), aes(x = BlockNumber, y =TimeToPeck,col=shownWindImage, fill=shownWindImage)) +
  stat_summary(fun = mean, geom = "point", size = 2) + 
  stat_summary(fun.data = mean_se, geom = "errorbar", size = 0.5, alpha=0.2) + 
  stat_summary(fun = mean, geom = "line", size = 0.8) + 
  scale_y_log10()+
  labs(x="Block Number",y="Log Time to Peck") +
  facet_grid(FirstPeck ~ as.factor(trainWindImage)) +
  scale_color_manual(values = my_colors) +
  scale_fill_manual(values = my_colors)   # Specify the color palette

p + theme_classic()+
  theme(legend.position = "none", text = element_text(size = 14),
        axis.title.y = element_text(margin = margin(t = 0, r = 20, b = 0, l = 0)), # Adjust margin for y-axis title
        axis.title.x = element_text(margin = margin(t = 20, r = 0, b = 0, l = 0))) # Adjust margin for x-axis title
```
