## Supplementary material for "Biologically inspired warning patterns deter birds from wind turbines": a_AddBird.m

```
trialInfoDlg = inputdlg({'Bird ID'}, 'Enter trial info');
trialInfo.bird = trialInfoDlg{1}; % Use index 1 instead of 0


trainList =  ["White","Red", "Black", "Bio"];


% Specify the directory path where you want to create the folder
folderPath =  fullfile('Birds/', trialInfo.bird ,'/'); % Replace this with your desired directory path

% Check if the directory exists
if ~exist(folderPath, 'dir')
    % If the directory doesn't exist, create it
    mkdir(folderPath);
    disp(['Folder created at ' folderPath]);
else
    disp(['Folder already exists at ' folderPath]);
end


% Display a dialog box to select a training type
trainIndex = listdlg('PromptString', 'Training_Pattern:', ... % Choose a training pattern
    'SelectionMode', 'single', 'ListString', trainList );

trainType = trainList{trainIndex};

% Define your variable
myVariable = trainType;


% Open a file for writing (create if it doesn't exist, overwrite if it does)
fileID = fopen(fullfile(folderPath, 'TrainPattern.txt'), 'w');

% Check if file is successfully opened
if fileID == -1
    error('Could not open file for writing');
end

% If trainType is a single string
if ischar(trainType)
    fprintf(fileID, '%s\n', trainType);
    
% If trainType is a cell array of strings
elseif iscellstr(trainType)
    for i = 1:numel(trainType)
        fprintf(fileID, '%s\n', trainType{i});
    end
    
else
    error('trainType must be a string or a cell array of strings.');
end

% Close the file
fclose(fileID);
```
