## Supplementary material for "Biologically inspired warning patterns deter birds from wind turbines": a_SetupScreens.m

```
!DisplaySwitch.exe /extend


% Use Bigger Screen
screenNumbers=Screen('Screens')
[width1, height1]=Screen('DisplaySize', 1);
[width2, height2]=Screen('DisplaySize', 2);

if(width2>width1)

primaryMonitorN = 2;
secondaryMonitorN = 1; 

else

primaryMonitorN = 1;
secondaryMonitorN = 2; 

end

% Define the full path to the nircmd.exe executable
nircmdPath = 'C:\Users\Localadmin_hangeorg\Documents\nircmd\nircmd.exe';

% Define the command to set monitor 1 as the primary display
command = [nircmdPath ' setprimarydisplay ' num2str(primaryMonitorN)];

% Execute the command
status = system(command);

% Check if the command executed successfully
if status == 0
    disp(['Primary display set']);
else
    disp('Error setting primary display.');
end


% 

PsychStartup

% Clear the workspace and the screen
sca;
close all;
clear;

% Here we call some default settings for setting up Psychtoolbox
PsychDefaultSetup(2);

% Get the screen numbers. This gives us a number for each of the screens
% attached to our computer.
screens = Screen('Screens')


    disp("Would you like to restart?")

    % Prompt the user for input
    response = input('"yes" or "no": ', 's'); % 's' flag specifies input as a string
    
    % Convert the input to lowercase for case-insensitive comparison
    response = lower(response);
    
    % Check the user's response
    if strcmp(response, 'yes')
        % Restart MATLAB using the system command
    system('matlab &');

    % Close MATLAB
    quit;
    end
```
