## Supplementary material for "Biologically inspired warning patterns deter birds from wind turbines": b_WindmillRandDotTrain.m

```
% Clear the workspace and the screen
sca;
close all;
clear;


primaryMonitorN = 2;
secondaryMonitorN = 1; 

secondaryMonitorN = 2; 


% Define the full path to the nircmd.exe executable
nircmdPath = 'C:\Users\Localadmin_hangeorg\Documents\nircmdd\nircmd.exe';


% Define the command to set monitor 1 as the primary display
command = [nircmdPath ' setprimarydisplay ' num2str(primaryMonitorN)];

% Execute the command
status = system(command);

% Check if the command executed successfully
if status == 0
    disp(['Primary display set']);
else
    disp('Error setting primary display.');
end


%Settings
%....................................................................
frameRate =165;

dotColor = [sqrt(0.2) sqrt(0.2) sqrt(0.2)];
dotSizePix = 30;

rng('shuffle');

timeLimit = 2*60; % Seconds

mouse = 'off';


screens = Screen('Screens')
if(length(screens)>1)
!DisplaySwitch.exe /extend
else
DisplayMonitorNumber = 0;
end


% Images and Layers
%......................................................................
% Load the background image
backgroundImage = imread('Backgrounds/Blank.png'); % Provide the path to your background image

dotColor = repelem(sqrt(0.2),3); % 20% grey (with sqrt nonlinearization)


% Set up Psychtoolbox
%......................................................................
PsychDefaultSetup(2);

Screen('Preference', 'SkipSyncTests', 1);
Screen('Preference', 'VisualDebugLevel', 01); 

% Open a window with a gray background
window = PsychImaging('OpenWindow', DisplayMonitorNumber, dotColor);

% Enable alpha blending for transparency
Screen('BlendFunction', window, GL_SRC_ALPHA, GL_ONE_MINUS_SRC_ALPHA);


%Setup Touch Screen Device
touchE = 0;
numTouchDevices = length(GetTouchDeviceIndices([], 1));

if(numTouchDevices>0)
  dev = min(GetTouchDeviceIndices([], 1));
  fprintf('Touch device properties:\n');
  info = GetTouchDeviceInfo(dev);
  disp(info);
  touchE = 1;
end

% Background Image
%...........................................................................


% Get the screen dimensions
[screenX, screenY] = Screen('WindowSize', window);

% Resize the background image to fit the screen dimensions
resizedBackgroundImage = imresize(backgroundImage, [screenY, screenX]);

if(touchE==1)
TouchQueueCreate(window, dev);
TouchQueueStart(dev);
end


% Trial Loops
%...........................................................................

endLoopGate=0;
while endLoopGate==0

Screen('FillRect', window, dotColor);
% Flip the screen
Screen('Flip', window);

pause(0.25);

% Create a texture for the background image
backgroundTexture = Screen('MakeTexture', window, resizedBackgroundImage);


    if strcmp(mouse,'off')
    HideCursor(window);
    end

gate=0;

dotType = "NA";

while (gate == 0)

% Start with a blank gray window
Screen('FillRect', window, dotColor);
Screen('Flip', window);

 % Check for key press
    [~, ~, keyCode] = KbCheck;
    
    if keyCode(KbName('1!')) % Check if '1'
        disp('You pressed "1" key.');
        dotType = 1;
        gate = 1;
    elseif keyCode(KbName('2@')) % Check if '2' 
        disp('You pressed "2" key.');
        dotType = 2;
        gate = 1;
    
    elseif keyCode(KbName('3#')) % Check if '3' or '#' key is pressed
            disp('You pressed "3" key.');
            dotType = 3;
            gate = 1;

    elseif keyCode(KbName('4$')) % Check if '4' or '$' key is pressed
        disp('You pressed "4" key.');
        dotType = 4;
        gate = 1;

    elseif keyCode(KbName('ESCAPE')) % Check for 'Escape' key
        endLoopGate = 1; % Exit loop if 'Escape' is pressed
        sca;
        endLoopGate=1;
        break;
    end

end

while KbCheck
% Do nothing, just wait for the key to be released
end

if(endLoopGate==0)

% Dot
%...........................................................................

% Generate a random angle in radians


% Set dot size and color
dotSize = round(screenY * 0.04);  % Radius of the dot in pixels
dotColor = [sqrt(0.2), sqrt(0.2), sqrt(0.2)];  % Dark gray color


% Calculate the coordinates of the dot center

radiusX = screenX/5;
radiusY = screenY/5;

if(dotType==1)
dotCenterX = round((screenX * 1/ 4) + (screenX * 1/ 20) + rand*(radiusX) * cos(rand*2*pi));
dotCenterY = round((screenY * 1/ 4) + (screenY * 1/ 20) + rand*(radiusY) * sin(rand*2*pi));

elseif(dotType==2)

dotCenterX = round((screenX * 3/ 4) - (screenX * 1/ 20) + rand*(radiusX) * cos(rand*2*pi));
dotCenterY = round((screenY * 1/ 4) + (screenY * 1/ 20) + rand*(radiusY) * sin(rand*2*pi));
elseif(dotType==3)

dotCenterX = round((screenX * 3/ 4) - (screenX * 1/ 20) + rand*(radiusX) * cos(rand*2*pi));
dotCenterY = round((screenY * 3/ 4) - (screenY * 1/ 20) + rand*(radiusY) * sin(rand*2*pi));
elseif(dotType==4)

dotCenterX = round((screenX * 1/ 4) + (screenX * 1/ 20) + rand*(radiusX) * cos(rand*2*pi));
dotCenterY = round((screenY * 3/ 4) - (screenY * 1/ 20) + rand*(radiusY) * sin(rand*2*pi));
end


% Create an array with the dot coordinates
dotCoords = [dotCenterX, dotCenterY];


newBackgroundTexture = backgroundTexture;


% Draw the dot onto the background texture
Screen('DrawDots', newBackgroundTexture, dotCoords, dotSize, dotColor, [], 2);


% Get the starting time
startTime = GetSecs;
tic
% Loop until either 360 frames have passed or the escape key is pressed

f=1;


% Trial
%...........................................................................


% Loop through frames
for frame = 1:10
    % Draw a gray screen
     Screen('FillRect', window, dotColor);
    
    % Flip the screen
    Screen('Flip', window);


end


if(touchE==1)
TouchQueueCreate(window, dev);
TouchQueueStart(dev);
HideCursor;
end


timeToClick = "NA";

tic
while toc < timeLimit % Run for time limit

    % Draw the background image
    Screen('DrawTexture', window, newBackgroundTexture);
    
    % Flip the screen to display the changes
    Screen('Flip', window);
   

    if(touchE==1)
          while TouchEventAvail(dev)
        % Process next touch event 'evt':
        evt = TouchEventGet(dev, window);

        % Touch blob id - Unique in the session at least as
        % long as the finger stays on the screen:
        id = evt.Keycode;

        % Keep the id's low, so we have to iterate over less blobcol slots
        % to save computation time:

           if (evt.Type == 2 ||  evt.Type == 3 ||  evt.Type == 4)
          % New touch point -> New blob!
          blobcol{id}.col = rand(3, 1);
          blobcol{id}.mul = 1.0;
          blobcol{id}.x = evt.MappedX;
          blobcol{id}.y = evt.MappedY;
          blobcol{id}.t = evt.Time;
          % Track time delta in msecs between touch point updates:
          blobcol{id}.dt = 0;


          if(blobcol{id}.y<screenY && blobcol{id}.x<screenX)
3
             disp([blobcol{id}.x,blobcol{id}.y]);
             disp([dotCenterX,dotCenterY]);

            if(((blobcol{id}.x-dotCenterX)^2 + (blobcol{id}.y-dotCenterY)^2)^0.5<dotSize*2)
            timeToClick = toc;

   

            disp("Touch Hit");
            break;
            
          end
          end

        end
    end
    end

     if ~strcmp(timeToClick,'NA')
        break
     end


 % Check for a click on the window
    [~, ~, buttons] = GetMouse(window);
    if any(buttons)
        % Get the clicked coordinates
        [clickX, clickY] = GetMouse(window);

        % Check if the click is inside dotInMask
        if(clickY<screenY && clickX<screenX)
        if( ((clickX-dotCenterX)^2 + (clickY-dotCenterY)^2)^0.5<dotSize*2)
            timeToClick = toc;

             disp("Click");

            break;
        end
        end

    end

    [~, ~, keyCode] = KbCheck;


    % Button Triggers
    if keyCode(KbName('1!')) % Check if '1' 
        timeToClick = toc;
        break;
    elseif keyCode(KbName('2@')) % Check if '2' 
        timeToClick = toc;
        break;
    
    elseif keyCode(KbName('3#')) % Check if '3' or '#' key is pressed
        timeToClick = toc;
        break;

    elseif keyCode(KbName('4$')) % Check if '4' or '$' key is pressed
        timeToClick = toc;
        break;

    elseif any(keyCode([KbName('space')]))
        timeToClick = toc;
        break;
    end

    % Check for the 'Escape' key to end the script
    if keyCode(KbName('Escape'))
        ShowCursor;
        sca;
        endLoopGate=1;
        break;
    end

f=f+1;
end

toc

while KbCheck
% Do nothing, just wait for the key to be released
end


% Get the current date and time as a datetime object
currentDateTime = datetime('now');

% Convert datetime object to date string
currentDate = datestr(currentDateTime, 'yyyy-mm-dd');

% Extract the time component as a string
currentTime = datestr(currentDateTime, 'HH:MM:SS');


disp(['Current date: ', currentDate]);
disp(['Current time: ', currentTime]);


disp(['Duration ' num2str(toc) ' seconds']);
disp(['Speed ' num2str(f/toc) ' fps']);

disp(['Type ' num2str(dotType)]);


if keyCode(KbName('Escape'))
    ShowCursor;
    endLoopGate=1;
    sca;
    
end
 

end
end


% Define the command to set monitor 1 as the primary display
command = [nircmdPath ' setprimarydisplay ' num2str(secondaryMonitorN)];

% Execute the command
status = system(command);

% Check if the command executed successfully
if status == 0
    disp(['Primary display set']);
else
    disp('Error setting primary display.');
end


% Close the window
sca;
```
