## Supplementary material for "Biologically inspired warning patterns deter birds from wind turbines": c_WindmillTrainingExperiment.m

```
% Clear the workspace and the screen
sca;
close all;
clear;

DisplayMonitorNumber=2;

primaryMonitorN = 1;
secondaryMonitorN = 2; 


% Define the full path to the nircmd.exe executable
nircmdPath = 'C:\Users\Localadmin_hangeorg\Documents\nircmd\nircmd.exe';


% Define the command to set monitor 1 as the primary display
command = [nircmdPath ' setprimarydisplay ' num2str(secondaryMonitorN)];

% Execute the command
status = system(command);

% Check if the command executed successfully
if status == 0
    disp(['Primary display set']);
else
    disp('Error setting primary display.');
end


%.....................................................................

DemoMode = 0; % Setting to 1 skips the save screen

WorkMonitor = "internal"; % Is the work monitor internal or external

screens = Screen('Screens');

screens = Screen('Screens');
if(length(screens)==1)
    DisplayMonitorNumber = 0;
end


%Bird
%....................................................................


% Specify the folder path
folderPath = 'Birds/';

% Get the list of items in the folder
itemList = dir(folderPath);

% Filter out only the folders
folderList = itemList([itemList.isdir]);

% Extract the folder names
folderNames = {folderList.name};

% Remove '.' and '..' entries
folderNames = folderNames(~ismember(folderNames, {'.', '..'}));

% Display a dialog box to select a bird
selectedFolderIndex = listdlg('PromptString', 'Choose a Bird:', ... % Choose a Bird
    'SelectionMode', 'single', 'ListString', folderNames);


trialList =  ["0_Static","1_Slow", "2_Medium", "3_Fast"];


% Display a dialog box to select a training type
trialIndex = listdlg('PromptString', 'Training_Trial:', ... % Choose a Bird
    'SelectionMode', 'single', 'ListString', trialList);

trialType = trialList{trialIndex};

selectedFolderName = folderNames{selectedFolderIndex};


% Specify the directory path where you want to create the folder
folderPath =  fullfile('Birds/', selectedFolderName,'/',trialType,'/'); % Replace this with your desired directory path

% Check if the directory exists
if ~exist(folderPath, 'dir')
    % If the directory doesn't exist, create it
    mkdir(folderPath);
    disp(['Folder created at ' folderPath]);
else
    disp(['Folder already exists at ' folderPath]);
end


   
% Get the list of items in the folder
itemList = dir(folderPath);

% Filter out only the CSV files
csvFiles = itemList(~[itemList.isdir]); % Select non-directory items
csvFiles = csvFiles(endsWith({csvFiles.name}, '.csv')); % Filter out CSV files

% Count the number of CSV files
existingTrials = numel(csvFiles);


% Specify the file path
filePath = fullfile(fullfile('Birds/', selectedFolderName,'/', 'TrainPattern.txt'));
fileID = fopen(filePath, 'r');

% Read the content of the file
content = fscanf(fileID, '%s');

% Close the file
fclose(fileID);

trainType = content;


% Define the command to set monitor 1 as the primary display
command = [nircmdPath ' setprimarydisplay ' num2str(primaryMonitorN)];

% Execute the command
status = system(command);

% Check if the command executed successfully
if status == 0
    disp(['Primary display set']);
else
    disp('Error setting primary display.');
end


%Settings
%....................................................................
frameRate =165;
speed = 20; % RPM

if strcmp(trialType,'0_Static')
speed = 0;
end

if strcmp(trialType,'1_Slow')
speed = 2.5;
end

if strcmp(trialType,'2_Medium')
speed = 5;
end

if strcmp(trialType,'3_Fast')
speed = 10;
end

if strcmp(trialType,'4_ExpSpeed')
speed = 20;
end


dotColor = [sqrt(0.2) sqrt(0.2) sqrt(0.2)];
dotSizeRatio = 0.04;

numFrames = frameRate * 60/speed;
rotationIncrement = 360/numFrames; % Rotation increment in degrees

expandR = 1; % Size relative to the target to expand


rng('shuffle');


innerDist = 1/5; % Distance of inner relative to screen width
outerDist = 1/3.5; % Distance of outer relative to screen width

timeLimit = 2*60; % Seconds

mouse = 'off';

touchMove =  0.05; % Touch Radius, how much does a touch have to move to count relative to the target size
touchTime = 0.15; % Time delay before a touch is registed.


overlap = 0; %


% Images and Layers
%......................................................................
% Load the background image
backgroundImage = imread('Backgrounds/Turbine.png'); % Provide the path to your background image

% Load the spinning image with alpha channel

if strcmp(trainType,'White')
[TurbineBlades, ~, alphaChannel] = imread('Blades/Spiral_White.png', 'png');
end

if strcmp(trainType,'Red')
[TurbineBlades, ~, alphaChannel] = imread('Blades/Spiral_Red.png', 'png');
end

if strcmp(trainType,'Black')
[TurbineBlades, ~, alphaChannel] = imread('Blades/Spiral_Black.png', 'png');
end

if strcmp(trainType,'Bio')
[TurbineBlades, ~, alphaChannel] = imread('Blades/Spiral_Bio.png', 'png');
end

dotColor = repelem(sqrt(0.2),3); % 20% grey (with sqrt nonlinearization)


% Pause for 3 seconds
pause(3);


% Set up Psychtoolbox
%......................................................................
PsychDefaultSetup(2);


Screen('Preference', 'SkipSyncTests', 1);
Screen('Preference', 'VisualDebugLevel', 0); 

% Open a window with a gray background
window = PsychImaging('OpenWindow', DisplayMonitorNumber, dotColor);

% Enable alpha blending for transparency
Screen('BlendFunction', window, GL_SRC_ALPHA, GL_ONE_MINUS_SRC_ALPHA);


%Setup Touch Screen Device
touchE = 0;
numTouchDevices = length(GetTouchDeviceIndices([], 1));

if(numTouchDevices>0)
  dev = min(GetTouchDeviceIndices([], 1));
  fprintf('Touch device properties:\n');
  info = GetTouchDeviceInfo(dev);
  disp(info);
  touchE = 1;
end

% Background Image
%...........................................................................


% Get the screen dimensions
[screenX, screenY] = Screen('WindowSize', window);

% Resize the background image to fit the screen dimensions
resizedBackgroundImage = imresize(backgroundImage, [screenY, screenX]);


% Create a texture for the spinning image / turbine
%...........................................................................

TurbineBlades = imresize(TurbineBlades, [screenY*0.85, screenY*0.85]); %Resize
alphaChannel = imresize(alphaChannel, [screenY*0.85, screenY*0.85]);

spinningTexture = Screen('MakeTexture', window, cat(3, TurbineBlades, alphaChannel));

% Get the size of the spinning image
[spinHeight, spinWidth, ~] = size(TurbineBlades);

% Calculate the position to draw the spinning image at the center of the screen
xPos = (screenX - spinWidth) / 2; % Adjust the x position to center the spinning image on the background
yPos = (screenY - spinHeight) / 2; % Adjust the y position to center the spinning image on the background


if(touchE==1)
TouchQueueCreate(window, dev);
TouchQueueStart(dev);
SetMouse(0,0,window);
HideCursor(window);
end

% Trial Loops
%...........................................................................

endLoopGate=0;
while endLoopGate==0

% Define initial rotation angle
rotationAngle = round((rand * 360)); % Initial rotation angle in degrees

 rotationAngle=90;


Screen('FillRect', window, dotColor);
% Flip the screen
Screen('Flip', window);

pause(0.25);

% Create a texture for the background image
backgroundTexture = Screen('MakeTexture', window, resizedBackgroundImage);


    if strcmp(mouse,'off')
    HideCursor(window);
    end

gate=0;

dotType = "NA";

while (gate == 0)

% Start with a blank gray window
Screen('FillRect', window, dotColor);
Screen('Flip', window);

 [~, ~, keyCode] = KbCheck;
    if any(keyCode([KbName('i')]))
        dotType = 'in';
        gate =1 ;

    end


    if any(keyCode([KbName('o')]))
        dotType = 'out';
        gate =1 ;
    end


    if any(keyCode([KbName('r')]))
    dotType = 'random';
    gate =1 ;
    end


        % Check for the 'Escape' key to end the script
    [~, ~, keyCode] = KbCheck;
    if keyCode(KbName('Escape'))
        ShowCursor;
        sca;
        endLoopGate=1;
        break;
    end

end

while KbCheck
% Do nothing, just wait for the key to be released
end


if strcmp(dotType,'random')
    if(rand<0.5)
    dotType = 'in';
    else
    dotType = 'out';
    end
end


disp(dotType);


leftRight = rand;

if(leftRight<0.5)
    leftRight="left";

else
    leftRight="right";
end

disp(leftRight);

if(endLoopGate==0)

% Dot
%...........................................................................

% Generate a random angle in radians

if strcmp(leftRight,'right') 
    randomAngle = (-35 + 70*rand) *pi / 180;
end


if strcmp(leftRight,'left') 
    randomAngle = (-35 +180+ 70*rand) *pi / 180;
end


% Set dot size and color
dotSize = round(screenY*dotSizeRatio);  % Radius of the dot in pixels
dotColor = [sqrt(0.2), sqrt(0.2), sqrt(0.2)];  % Dark gray color
expandDot = dotSize*expandR;

if strcmp(dotType,'in') 
    radius = screenX*innerDist;
end


if strcmp(dotType,'out') 
    radius = screenX*outerDist;
end

% Calculate the coordinates of the dot center
dotCenterX = round(screenX / 2 + (radius) * cos(randomAngle));
dotCenterY = round(screenY / 2 + (radius) * sin(randomAngle));


% Create an array with the dot coordinates
dotCoords = [dotCenterX, dotCenterY];


newBackgroundTexture = backgroundTexture;


% Draw the dot onto the background texture
Screen('DrawDots', newBackgroundTexture, dotCoords, dotSize, dotColor, [], 2);


% Get the starting time
startTime = GetSecs;
tic
% Loop until either 360 frames have passed or the escape key is pressed

f=1;


% Trial
%...........................................................................


% Loop through frames
for frame = 1:10
    % Draw a gray screen
     Screen('FillRect', window, dotColor);
    
    % Flip the screen
    Screen('Flip', window);


end


ba1="NA";
ba2="NA";
ba3="NA";


timeToClick = "NA";

peckX = "NA";
peckY = "NA";
peckD = "NA";

clicks = [];

pTX = 0; % Record prior touch coordinates
pTY = 0; % Record prior touch coordinates
pTT = 0; % Record prior touch time


pecks = [];

randomAngle = randomAngle*180/pi;
if(randomAngle<0)
randomAngle = 360+randomAngle;
end

if(overlap==0)

ba1 = rotationAngle+85;
ba2 = rotationAngle+85+120;
ba3 = rotationAngle+85+240;

if(ba1>360)ba1=ba1-360; end
if(ba2>360)ba2=ba2-360; end
if(ba3>360)ba3=ba3-360; end

bd1 = abs(ba1 - randomAngle);
if bd1 > 180  
bd1 = 360 - bd1;
end

bd2 = abs(ba2 - randomAngle);
if bd2 > 180  
bd2 = 360 - bd2;
end

bd3 = abs(ba3 - randomAngle);
if bd3 > 180  
bd3 = 360 - bd3;
end


% Define a 1D array
array = [bd1,bd2,bd3];

% Calculate the minimum value
minValue = min(array);


if(minValue<10)
    rotationAngle = rotationAngle+10;
    if rotationAngle>360
        rotationAngle = rotationAngle-360;
    end
end
end

disp("Trial Start");
disp("...................");

if(touchE==1)
TouchQueueCreate(window, dev);
TouchQueueStart(dev);
HideCursor(window);
SetMouse(0,0,window)
end


tic
while toc < timeLimit % Run for time limit

    peck=0;

    % Draw the background image
    Screen('DrawTexture', window, newBackgroundTexture);
    
    % Draw the spinning image onto the background texture at the center of the screen with the current rotation angle
    Screen('DrawTexture', window, spinningTexture, [], ...
        [xPos, yPos, xPos + spinWidth, yPos + spinHeight], rotationAngle);
    
    % Flip the screen to display the changes
    Screen('Flip', window);
    
    % Increment the rotation angle
    rotationAngle = rotationAngle + rotationIncrement;
    if(rotationAngle>360)
        rotationAngle=rotationAngle-360;
    end


    if(touchE==1)
          while TouchEventAvail(dev)
        % Process next touch event 'evt':
        evt = TouchEventGet(dev, window);

        % Touch blob id - Unique in the session at least as
        % long as the finger stays on the screen:
        id = evt.Keycode;

        % Keep the id's low, so we have to iterate over less blobcol slots
        % to save computation time:

          if (evt.Type == 2 ||  evt.Type == 3 ||  evt.Type == 4)
          % New touch point -> New blob!
          blobcol{id}.col = rand(3, 1);
          blobcol{id}.mul = 1.0;
          blobcol{id}.x = evt.MappedX;
          blobcol{id}.y = evt.MappedY;
          blobcol{id}.t = evt.Time;
          % Track time delta in msecs between touch point updates:
          blobcol{id}.dt = 0;

          if(((blobcol{id}.x-pTX)^2 + (blobcol{id}.y-pTY)^2)^0.5>dotSize*touchMove || toc-pTT > touchTime)

            pTT = toc;
            pTX= blobcol{id}.x;
            pTY = blobcol{id}.y;

          peck=1;
          disp("Touch");

          clicks(end+1, :) = [blobcol{id}.x, blobcol{id}.y, toc];

          if(blobcol{id}.y<screenY && blobcol{id}.x<screenX)

             disp([blobcol{id}.x,blobcol{id}.y]);
             disp([dotCenterX,dotCenterY,((blobcol{id}.x-dotCenterX)^2 + (blobcol{id}.y-dotCenterY)^2)^0.5]);

            if(((blobcol{id}.x-dotCenterX)^2 + (blobcol{id}.y-dotCenterY)^2)^0.5<dotSize+expandDot)
            timeToClick = toc;
            peckX = blobcol{id}.x;
            peckY = blobcol{id}.y;
            peckD = ((blobcol{id}.x-dotCenterX)^2 + (blobcol{id}.y-dotCenterY)^2)^0.5;
            break;
            
          end
          end
          end
        end
    end
    end


 % Check for a click on the window
    [~, ~, buttons] = GetMouse(window);
    if any(buttons)
        % Get the clicked coordinates
        [clickX, clickY] = GetMouse(window);

        if(((clickX-pTX)^2 + (clickY-pTY)^2)^0.5>dotSize*touchMove || toc-pTT > touchTime)
        
            pTT = toc;
            pTX= clickX;
            pTY = clickY;
        
        clicks(end+1, :) = [clickX, clickY, toc];

        disp("Click");

        % Check if the click is inside dotInMask
        if(clickY<screenY && clickX<screenX)
        if( ((clickX-dotCenterX)^2 + (clickY-dotCenterY)^2)^0.5<dotSize+expandDot)
            timeToClick = toc;
            peckX = clickX;
            peckY = clickY;
            peckD = ((clickX-dotCenterX)^2 + (clickY-dotCenterY)^2)^0.5;

            break;
        end
        end
        end
    end

    [~, ~, keyCode] = KbCheck;


    % Button Triggers
    if any(keyCode([KbName('i')]))
            timeToClick = toc;
            break;
    end

    if any(keyCode([KbName('o')]))
            timeToClick = toc;
            break;
    end

    if any(keyCode([KbName('r')]))
            timeToClick = toc;
            break;
    end

    if any(keyCode([KbName('space')]))
        timeToClick = toc;
        break;
    end


     if ~strcmp(timeToClick,'NA')
        break
     end


    % Check for the 'Escape' key to end the script
    if keyCode(KbName('Escape'))
        ShowCursor;
        sca;
        endLoopGate=1;
        break;
    end

f=f+1;
end

toc


        while KbCheck
        % Do nothing, just wait for the key to be released
        end

% Measures
%...........................................................................
ba1 = rotationAngle+95;
ba2 = rotationAngle+95+120;
ba3 = rotationAngle+95+240;

if(ba1>360)ba1=ba1-360; end
if(ba2>360)ba2=ba2-360; end
if(ba3>360)ba3=ba3-360; end


bd1 = abs(ba1 - randomAngle);
if bd1 > 180  
bd1 = 360 - bd1;
end

bd2 = abs(ba2 - randomAngle);
if bd2 > 180  
bd2 = 360 - bd2;
end

bd3 = abs(ba3 - randomAngle);
if bd3 > 180  
bd3 = 360 - bd3;
end


% Define a 1D array
array = [bd1,bd2,bd3];

% Calculate the minimum value
minValue = min(array);


disp("...................");
disp("Trial End");


%Output
%...........................................................................

state = "Fail";

if~strcmp(timeToClick,"NA")
state = "Succ";
end


% Get the current date and time as a datetime object
currentDateTime = datetime('now');

% Convert datetime object to date string
currentDate = datestr(currentDateTime, 'yyyy-mm-dd');

% Extract the time component as a string
currentTime = datestr(currentDateTime, 'HH:MM:SS');


% Define your header and data
header = {'BirdID','TrialType','trainWindImage', 'Speed', 'Number','Date','Time', 'InOut', 'LeftRight', 'TimeToPeck', 'SuccFail', 'PeckX', 'PeckY','PeckD', 'clicks','TargetX', 'TargetY', 'TargetSize', 'TargetAngle', 'RadiusCentre', 'TurbineAngle', 'Blade1Angle', 'Blade2Angle', 'Blade3Angle', 'Blade1Dist', 'Blade2Dist', 'Blade3Dist', 'MinBladeDist','expandTarget','frameRate','dotCol','timeOutTime','frames','screenWidth','screenHeight'}; % Example header


speedStr = num2str(speed);
existingTrialsStr = num2str(existingTrials+1);
dotCenterXStr = num2str(dotCenterX);
dotCenterYStr = num2str(dotCenterY);
dotSizeStr = num2str(dotSize);
randomAngleStr = num2str(randomAngle);
radiusStr = num2str(radius);
rotationAngleStr = num2str(rotationAngle);
ba1Str = num2str(ba1);
ba2Str = num2str(ba2);
ba3Str = num2str(ba3);
bd1Str = num2str(bd1);
bd2Str = num2str(bd2);
bd3Str = num2str(bd3);
minValueStr = num2str(minValue);

peckX = num2str(peckX);
peckY = num2str(peckY);

% Convert the array to a format with semicolons to separate columns and colons to separate rows
formattedClicks = '';

if(length(clicks)>0)
% Loop through each row of the array
for i = 1:size(clicks, 1)
    % Convert the current row to a string with semicolons
    rowString = sprintf('%d %d %d', clicks(i, 1), clicks(i, 2), clicks(i, 3));
    
    % Append a colon to separate rows
    formattedClicks = [formattedClicks, rowString, ';'];
end

% Remove the last colon
formattedClicks(end) = [];
else

formattedClicks = 'NA';

end


colorString = sprintf('%f %f %f', dotColor);


% Combine header and data into a single cell array
rowData = {selectedFolderName,trialType,trainType, speedStr, existingTrialsStr,currentDate,currentTime, dotType, leftRight, timeToClick, state, peckX, peckY,num2str(peckD), formattedClicks, dotCenterXStr, dotCenterYStr, dotSizeStr, randomAngleStr, radiusStr, rotationAngleStr, ba1Str, ba2Str, ba3Str, bd1Str, bd2Str, bd3Str, minValueStr,num2str(expandDot),num2str(f/toc),colorString,num2str(timeLimit),num2str(f),num2str(screenX),num2str(screenY)}; % Example data
data = [header; rowData];

% Create the filename based on the loop index
if ~strcmp(timeToClick, "NA")
    filename = [num2str(existingTrials+1), '_', dotType, '_S', '.csv'];
else
    filename = [num2str(existingTrials+1), '_', dotType, '_F', '.csv'];
end

% Open the file for writing
fileID = fopen(fullfile(folderPath, filename), 'w');


% Write the data rows
for i = 1:size(data, 1)
    fprintf(fileID, '%s,', data{i,1:end-1});
    fprintf(fileID, '%s\n', data{i,end});
end

% Close the file
fclose(fileID);


existingTrials = existingTrials + 1; % Corrected variable name

disp(['Type: ', dotType]);
disp(['Current date: ', currentDate]);
disp(['Current time: ', currentTime]);


disp(['Duration ' num2str(timeToClick) ' seconds']);
disp(['Speed ' num2str(f/toc) ' fps']);
disp(dotType);


if keyCode(KbName('Escape'))
    ShowCursor;
    endLoopGate=1;
    sca;
    
end
 

end
end


% Define the command to set monitor 1 as the primary display
command = [nircmdPath ' setprimarydisplay ' num2str(secondaryMonitorN)];

% Execute the command
status = system(command);

% Check if the command executed successfully
if status == 0
    disp(['Primary display set']);
else
    disp('Error setting primary display.');
end


% Close the window
sca;
```
