## Supplementary material for "Biologically inspired warning patterns deter birds from wind turbines": d_WindmillExperiment.m

```
% Clear the workspace and the screen
sca;
close all;
clear;

DisplayMonitorNumber=1;

primaryMonitorN = 1;
secondaryMonitorN = 2; 

% Define the full path to the nircmd.exe executable
nircmdPath = 'C:\Users\Localadmin_hangeorg\Documents\nircmd\nircmd.exe';


% Define the command to set monitor 1 as the primary display
command = [nircmdPath ' setprimarydisplay ' num2str(secondaryMonitorN)];

% Execute the command
status = system(command);

% Check if the command executed successfully
if status == 0
    disp(['Primary display set']);
else
    disp('Error setting primary display.');
end


%.....................................................................

DemoMode = 0; % Setting to 1 skips the save screen

WorkMonitor = "internal"; % Is the work monitor internal or external

screens = Screen('Screens');

screens = Screen('Screens');
if(length(screens)==1)
    DisplayMonitorNumber = 0;
end


%Bird
%....................................................................


% Specify the folder path
folderPath = 'Birds/';

% Get the list of items in the folder
itemList = dir(folderPath);

% Filter out only the folders
folderList = itemList([itemList.isdir]);

% Extract the folder names
folderNames = {folderList.name};

% Remove '.' and '..' entries
folderNames = folderNames(~ismember(folderNames, {'.', '..'}));

% Display a dialog box to select a bird
selectedFolderIndex = listdlg('PromptString', 'Choose a Bird:', ... % Choose a Bird
    'SelectionMode', 'single', 'ListString', folderNames);


trialList =  ["4_MultiDot_ExpSpeed","5_Comparative_Experiment"];


% Display a dialog box to select a training type
trialIndex = listdlg('PromptString', 'Experiment:', ... % Choose a Bird
    'SelectionMode', 'single', 'ListString', trialList);

trialType = trialList{trialIndex};

selectedFolderName = folderNames{selectedFolderIndex};


% Specify the directory path where you want to create the folder
folderPath =  fullfile('Birds/', selectedFolderName,'/',trialType,'/'); % Replace this with your desired directory path

% Check if the directory exists
if ~exist(folderPath, 'dir')
    % If the directory doesn't exist, create it
    mkdir(folderPath);
    disp(['Folder created at ' folderPath]);
else
    disp(['Folder already exists at ' folderPath]);
end


   
% Get the list of items in the folder
itemList = dir(folderPath);

% Filter out only the CSV files
csvFiles = itemList(~[itemList.isdir]); % Select non-directory items
csvFiles = csvFiles(endsWith({csvFiles.name}, '.csv')); % Filter out CSV files

% Count the number of CSV files
existingTrials = numel(csvFiles);


% Specify the file path
filePath = fullfile(fullfile('Birds/', selectedFolderName,'/', 'TrainPattern.txt'));
fileID = fopen(filePath, 'r');

% Read the content of the file
content = fscanf(fileID, '%s');

% Close the file
fclose(fileID);

trainType = content;


% Define the command to set monitor 1 as the primary display
command = [nircmdPath ' setprimarydisplay ' num2str(primaryMonitorN)];

% Execute the command
status = system(command);

% Check if the command executed successfully
if status == 0
    disp(['Primary display set']);
else
    disp('Error setting primary display.');
end


%Settings
%....................................................................
frameRate =165;
speed = 20; % RPM

if strcmp(trialType,'0_Static')
speed = 0;
end

if strcmp(trialType,'1_Slow')
speed = 2.5;
end

if strcmp(trialType,'2_Medium')
speed = 5;
end

if strcmp(trialType,'3_Fast')
speed = 10;
end

if strcmp(trialType,'4_ExpSpeed')
speed = 20;
end


dotColor = [sqrt(0.2) sqrt(0.2) sqrt(0.2)];
dotSizeRatio = 0.04;

numFrames = frameRate * 60/speed;
rotationIncrement = 360/numFrames; % Rotation increment in degrees

expandR = 0; % Size relative to the target to expand


rng('shuffle');


innerDist = 1/5; % Distance of inner relative to screen width
outerDist = 1/3.5; % Distance of outer relative to screen width

timeLimit = 2*60; % Seconds


mouse = 'off';

touchMove =  0.25; % Touch Radius, how much does a touch have to move to count relative to the target size
touchTime = 0.3; % Time delay before a touch is registed.


overlap = 0; %


% Images and Layers
%......................................................................
% Load the background image
backgroundImage = imread('Backgrounds/Turbine.png'); % Provide the path to your background image

% Load the spinning image with alpha channel


[WhiteBlades, ~, alphaChannel] = imread('Blades/Spiral_White.png', 'png');

[BlackBlades, ~, alphaChannel] = imread('Blades/Spiral_Black.png', 'png');

[RedBlades, ~, alphaChannel] = imread('Blades/Spiral_Red.png', 'png');

[BioBlades, ~, alphaChannel] = imread('Blades/Spiral_Bio.png', 'png');


dotColor = repelem(sqrt(0.2),3); % 20% grey (with sqrt nonlinearization)


% Pause for 3 seconds
pause(3);


% Set up Psychtoolbox
%......................................................................
PsychDefaultSetup(2);

Screen('Preference', 'SkipSyncTests', 1);
Screen('Preference', 'VisualDebugLevel', 0); 

% Open a window with a gray background
window = PsychImaging('OpenWindow', DisplayMonitorNumber, dotColor);

% Enable alpha blending for transparency
Screen('BlendFunction', window, GL_SRC_ALPHA, GL_ONE_MINUS_SRC_ALPHA);


%Setup Touch Screen Device
touchE = 0;
numTouchDevices = length(GetTouchDeviceIndices([], 1));

if(numTouchDevices>0)
  dev = min(GetTouchDeviceIndices([], 1));
  fprintf('Touch device properties:\n');
  info = GetTouchDeviceInfo(dev);
  disp(info);
  touchE = 1;
end

% Background Image
%...........................................................................


% Get the screen dimensions
[screenX, screenY] = Screen('WindowSize', window);

% Resize the background image to fit the screen dimensions
resizedBackgroundImage = imresize(backgroundImage, [screenY, screenX]);


% Create a texture for the spinning image / turbine
%...........................................................................

WhiteBlades = imresize(WhiteBlades, [screenY*0.85, screenY*0.85]); %Resize
BlackBlades = imresize(BlackBlades, [screenY*0.85, screenY*0.85]); %Resize
RedBlades = imresize(RedBlades, [screenY*0.85, screenY*0.85]); %Resize
BioBlades = imresize(BioBlades, [screenY*0.85, screenY*0.85]); %Resize

alphaChannel = imresize(alphaChannel, [screenY*0.85, screenY*0.85]);


if strcmp(trainType,'White')
TurbineBlades = WhiteBlades;
end


if strcmp(trainType,'Black')
TurbineBlades = BlackBlades;
end


if strcmp(trainType,'Red')
TurbineBlades = RedBlades;
end


if strcmp(trainType,'Bio')
TurbineBlades = BioBlades;
end


% Get the size of the spinning image
[spinHeight, spinWidth, ~] = size(TurbineBlades);

% Calculate the position to draw the spinning image at the center of the screen
xPos = (screenX - spinWidth) / 2; % Adjust the x position to center the spinning image on the background
yPos = (screenY - spinHeight) / 2; % Adjust the y position to center the spinning image on the background


% Trial Loops
%...........................................................................

if(touchE==1)
TouchQueueCreate(window, dev);
TouchQueueStart(dev);
SetMouse(0,0,window);
HideCursor(window);
end


endLoopGate=0;
while endLoopGate==0


    endTime=0;

    % Define initial rotation angle
rotationAngle = round(rand * 360); % Initial rotation angle in degrees


Screen('FillRect', window, dotColor);
% Flip the screen
Screen('Flip', window);

pause(0.25);

% Create a texture for the background image
backgroundTexture = Screen('MakeTexture', window, resizedBackgroundImage);


    if strcmp(mouse,'off')
    HideCursor(window);
    end

gate=0;

bladeType = "NA";

while (gate == 0)

% Start with a blank gray window
Screen('FillRect', window, dotColor);
Screen('Flip', window);

 [~, ~, keyCode] = KbCheck;

    if keyCode(KbName('1!')) % Check if '1'
        disp('You pressed "1" key.');
        bladeType = 'White';
        gate = 1;
    elseif keyCode(KbName('2@')) % Check if '2' 
        disp('You pressed "2" key.');
        bladeType = 'Black';
        gate = 1;
    
    elseif keyCode(KbName('3#')) % Check if '3' or '#' key is pressed
        disp('You pressed "3" key.');
        bladeType = 'Red';
        gate = 1;

    elseif keyCode(KbName('4$')) % Check if '4' or '$' key is pressed
        disp('You pressed "4" key.');
        bladeType = 'Bio';
        gate = 1;

    end

        % Check for the 'Escape' key to end the script
    [~, ~, keyCode] = KbCheck;
    if keyCode(KbName('Escape'))
        ShowCursor;
        sca;
        endLoopGate=1;
        break;
    end

end

        while KbCheck
        % Do nothing, just wait for the key to be released
        end


if(endLoopGate==0)


% Change Spinning
%...........................................................................

if strcmp(trialType,'5_Comparative_Experiment')

if strcmp(bladeType,'White')
TurbineBlades = WhiteBlades;
end


if strcmp(bladeType,'Black')
TurbineBlades = BlackBlades;
end


if strcmp(bladeType,'Red')
TurbineBlades = RedBlades;
end


if strcmp(bladeType,'Bio')
TurbineBlades = BioBlades;
end

else


bladeType = trainType;

end

spinningTexture = Screen('MakeTexture', window, cat(3, TurbineBlades, alphaChannel));


% Create Dots
%...........................................................................

% Set dot size and color
dotSize = round(screenY*dotSizeRatio);  % Radius of the dot in pixels
dotColor = [sqrt(0.2), sqrt(0.2), sqrt(0.2)];  % Dark gray color
expandDot = dotSize*expandR;

% Dot I

if(rand<0.5)
    leftRightI="left";

else
    leftRightI="right";
end


if strcmp(leftRightI,'right') 
    randomAngleI = (-35 + 70*rand) *pi / 180;
end


if strcmp(leftRightI,'left') 
    randomAngleI = (-35 +180+ 70*rand) *pi / 180;
end


radiusI = screenX*innerDist;

% Calculate the coordinates of the dot center
dotCenterIX = round(screenX / 2 + (radiusI) * cos(randomAngleI));
dotCenterIY = round(screenY / 2 + (radiusI) * sin(randomAngleI));

% Create an array with the dot coordinates
dotCoordsI = [dotCenterIX, dotCenterIY];


% Dot O


if(rand<0.5)
    leftRightO="left";

else
    leftRightO="right";
end


if strcmp(leftRightO,'right') 
    randomAngleO = (-35 + 70*rand) *pi / 180;
end


if strcmp(leftRightO,'left') 
    randomAngleO = (-35 +180+ 70*rand) *pi / 180;
end


radiusO = screenX*outerDist;


% Calculate the coordinates of the dot center
dotCenterOX = round(screenX / 2 + (radiusO) * cos(randomAngleO));
dotCenterOY = round(screenY / 2 + (radiusO) * sin(randomAngleO));

% Create an array with the dot coordinates
dotCoordsO = [dotCenterOX, dotCenterOY];


randomAngleI = randomAngleI*180/pi;

randomAngleO = randomAngleO*180/pi;

if(randomAngleI<0)
randomAngleI = 360+randomAngleI;
end

if(randomAngleO<0)
randomAngleO = 360+randomAngleO;
end


% Create Backgrounds
%...........................................................................


% Draw the dot onto the background textures
Screen('DrawDots', backgroundTexture, dotCoordsI, dotSize, dotColor, [], 2);
Screen('DrawDots', backgroundTexture, dotCoordsO, dotSize, dotColor, [], 2);


% Get the starting time
startTime = GetSecs;
tic
% Loop until either 360 frames have passed or the escape key is pressed

f=1;


% Trial 1
%...........................................................................


% Loop through frames
for frame = 1:10
    % Draw a gray screen
     Screen('FillRect', window, dotColor);
    
    % Flip the screen
    Screen('Flip', window);


end


ba1_1="NA";
ba2_1="NA";
ba3_1="NA";

ba1_2="NA";
ba2_2="NA";
ba3_2="NA";


timeToClick = "NA";
timeToClick1 = "NA";
timeToClick2 = "NA";

timeToClickI = "NA";
timeToClickO = "NA";

peckX = "NA";
peckY = "NA";

peckX1 = "NA";
peckX2 = "NA";

peckY1 = "NA";
peckY2= "NA";

peckD = "NA";

peckD1 = "NA";
peckD2 = "NA";

clicks1 = [];

pTX = 0; % Record prior touch coordinates
pTY = 0; % Record prior touch coordinates
pTT = 0; % Record prior touch time


disp("Trial 1 Start");
disp("...................");


firstClick='NA';


if(touchE==1)
TouchQueueCreate(window, dev);
TouchQueueStart(dev);
SetMouse(0,0,window);
HideCursor(window);
end

tic
while toc < timeLimit % Run for time limit

    
    
    peck=0;

    % Draw the background image
    Screen('DrawTexture', window, backgroundTexture);
    
    % Draw the spinning image onto the background texture at the center of the screen with the current rotation angle
    Screen('DrawTexture', window, spinningTexture, [], ...
        [xPos, yPos, xPos + spinWidth, yPos + spinHeight], rotationAngle);
    
    % Flip the screen to display the changes
    Screen('Flip', window);
    
    % Increment the rotation angle
    rotationAngle = rotationAngle + rotationIncrement;
    if(rotationAngle>360)
        rotationAngle=rotationAngle-360;
    end


    if(touchE==1)
          while TouchEventAvail(dev)
        % Process next touch event 'evt':
        evt = TouchEventGet(dev, window);

        % Touch blob id - Unique in the session at least as
        % long as the finger stays on the screen:
        id = evt.Keycode;

        % Keep the id's low, so we have to iterate over less blobcol slots
        % to save computation time:

          if (evt.Type == 2 ||  evt.Type == 3 ||  evt.Type == 4)
          % New touch point -> New blob!
          blobcol{id}.col = rand(3, 1);
          blobcol{id}.mul = 1.0;
          blobcol{id}.x = evt.MappedX;
          blobcol{id}.y = evt.MappedY;
          blobcol{id}.t = evt.Time;
          % Track time delta in msecs between touch point updates:
          blobcol{id}.dt = 0;

          if(((blobcol{id}.x-pTX)^2 + (blobcol{id}.y-pTY)^2)^0.5>dotSize*touchMove || toc-pTT > touchTime)


            pTT = toc;
            pTX= blobcol{id}.x;
            pTY = blobcol{id}.y;

          peck=1;
          disp("Touch");

          clicks1(end+1, :) = [blobcol{id}.x, blobcol{id}.y, toc];

          if(blobcol{id}.y<screenY && blobcol{id}.x<screenX)

            if(((blobcol{id}.x-dotCenterIX)^2 + (blobcol{id}.y-dotCenterIY)^2)^0.5<dotSize+expandDot)
            timeToClick = toc;
            firstClick = 'in';
            peckX = blobcol{id}.x;
            peckY = blobcol{id}.y;
            peckD =((blobcol{id}.x-dotCenterIX)^2 + (blobcol{id}.y-dotCenterIY)^2)^0.5;
disp(['first click = ', num2str(firstClick)]);
            elseif (((blobcol{id}.x-dotCenterOX)^2 + (blobcol{id}.y-dotCenterOY)^2)^0.5<dotSize+expandDot)
            timeToClick = toc;
            firstClick = 'out';
            peckX = blobcol{id}.x;
            peckY = blobcol{id}.y;
            peckD =((blobcol{id}.x-dotCenterOX)^2 + (blobcol{id}.y-dotCenterOY)^2)^0.5;
disp(['first click = ', num2str(firstClick)]);
            break;
            
          end
          end
          end
        end
    end
    end

     if ~strcmp(timeToClick,'NA')
        break
     end


 % Check for a click on the window
    [~, ~, buttons] = GetMouse(window);
    if any(buttons)
        % Get the clicked coordinates
        [clickX, clickY] = GetMouse(window);

        if(((clickX-pTX)^2 + (clickY-pTY)^2)^0.5>dotSize*touchMove || toc-pTT > touchTime)
        
            pTT = toc;
            pTX= clickX;
            pTY = clickY;
        
        clicks1(end+1, :) = [clickX, clickY, toc];

        disp("Click");

        % Check if the click is inside dotInMask
        if(clickY<screenY && clickX<screenX)
        if( ((clickX-dotCenterIX)^2 + (clickY-dotCenterIY)^2)^0.5<dotSize+expandDot)
            timeToClick = toc;
            peckX = clickX;
            peckY = clickY;
            peckD = ((clickX-dotCenterIX)^2 + (clickY-dotCenterIY)^2)^0.5;
            firstClick = 'in';
disp(['first click = ', num2str(firstClick)]);
        elseif( ((clickX-dotCenterOX)^2 + (clickY-dotCenterOY)^2)^0.5<dotSize+expandDot)
            timeToClick = toc;
            peckX = clickX;
            peckY = clickY;
            peckD = ((clickX-dotCenterOX)^2 + (clickY-dotCenterOY)^2)^0.5;
            firstClick = 'out';
disp(['first click = ', num2str(firstClick)]);
        break;
        end
        end
        end
    end

    [~, ~, keyCode] = KbCheck;


    % Button Triggers
    if any(keyCode([KbName('i')]))
            timeToClick = toc;
            firstClick = 'in';
disp(['first click = ', num2str(firstClick)]);
            break;
    end

    if any(keyCode([KbName('o')]))
            timeToClick = toc;
            firstClick = 'out';
disp(['first click = ', num2str(firstClick)]);
            break;
    end


    % Check for the 'Escape' key to end the script
    if keyCode(KbName('Escape'))
        ShowCursor;
        sca;
        endLoopGate=1;
        break;
    end

f=f+1;
end

toc

endTime = toc;


while KbCheck
% Do nothing, just wait for the key to be released
end


Screen('FillRect', window, dotColor);
Screen('Flip', window);


% Measures
%...........................................................................


gate2=0;
gate=0;

timeToClick1 = timeToClick;
peckX1 = peckX;
peckY1 = peckY;
peckD1 = peckD;

clicks2 = [];


while(gate2 == 0)

    if(gate==1)
        gate2=1
    end


f1 = f; 
rotationAngle1 = rotationAngle;


ba1_1 = rotationAngle+85;
ba2_1 = rotationAngle+85+120;
ba3_1 = rotationAngle+85+240;

if(ba1_1>360)ba1_1=ba1_1-360; end
if(ba2_1>360)ba2_1=ba2_1-360; end
if(ba3_1>360)ba3_1=ba3_1-360; end

bd1_1 = 'NA';
bd2_1 = 'NA';
bd3_1 = 'NA';
minValue1= 'NA';

if strcmp(firstClick,'in')

bd1_1 = abs(ba1_1 - randomAngleI);
if bd1_1 > 180  
bd1_1 = 360 - bd1_1;
end

bd2_1 = abs(ba2_1 - randomAngleI);
if bd2_1 > 180  
bd2_1 = 360 - bd2_1;
end

bd3_1 = abs(ba3_1 - randomAngleI);
if bd3_1 > 180  
bd3_1 = 360 - bd3_1;
end

elseif  strcmp(firstClick,'out')

bd1_1 = abs(ba1_1 - randomAngleO);
if bd1_1 > 180  
bd1_1 = 360 - bd1_1;
end

bd2_1 = abs(ba2_1 - randomAngleO);
if bd2_1 > 180  
bd2_1 = 360 - bd2_1;
end

bd3_1 = abs(ba3_1 - randomAngleO);
if bd3_1 > 180  
bd3_1 = 360 - bd3_1;
end

end


% Define a 1D array
array = [bd1_1,bd2_1,bd3_1];

% Calculate the minimum value
minValue1 = min(array);


% Get the current date and time as a datetime object
currentDateTime = datetime('now');

% Convert datetime object to date string
currentDate = datestr(currentDateTime, 'yyyy-mm-dd');

% Extract the time component as a string
currentTime1 = datestr(currentDateTime, 'HH:MM:SS');

disp(['Current date: ', currentDate]);
disp(['Current time: ', currentTime1]);


% Trial 2
%...........................................................................


% Define initial rotation angle
rotationAngle = round(rand * 360); % Initial rotation angle in degrees


if strcmp(firstClick,'in')

backgroundTexture = Screen('MakeTexture', window, resizedBackgroundImage);
Screen('DrawDots', backgroundTexture, dotCoordsO, dotSize, dotColor, [], 2);

dotCenterRX = dotCenterOX;
dotCenterRY = dotCenterOY;

else


backgroundTexture = Screen('MakeTexture', window, resizedBackgroundImage);
Screen('DrawDots', backgroundTexture, dotCoordsI, dotSize, dotColor, [], 2);

dotCenterRX = dotCenterIX;
dotCenterRY = dotCenterIY;

end


ba1I="NA";
ba2I="NA";
ba3I="NA";


timeToClick = "NA";


peckX = "NA";
peckY = "NA";
peckD = "NA";

pTX = 0; % Record prior touch coordinates
pTY = 0; % Record prior touch coordinates
pTT = 0; % Record prior touch time


if(endTime<timeLimit)

while(gate == 0)

% Start with a blank gray window
Screen('FillRect', window, dotColor);
Screen('Flip', window);

 [~, ~, keyCode] = KbCheck;

    if keyCode(KbName('1!')) % Check if '1'
        disp('You pressed "1" key.');
        gate = 1;
    elseif keyCode(KbName('2@')) % Check if '2' 
        disp('You pressed "2" key.');
        gate = 1;
    elseif keyCode(KbName('3#')) % Check if '3' or '#' key is pressed
        disp('You pressed "3" key.');
        gate = 1;
    elseif keyCode(KbName('4$')) % Check if '4' or '$' key is pressed
        disp('You pressed "4" key.');
        gate = 1;
    elseif keyCode(KbName('k')) % Check if '4' or '$' key is pressed
        disp('You pressed "i" key.');
        gate = 1;
        firstClick = 'in';

    elseif keyCode(KbName('l')) % Check if '4' or '$' key is pressed
        disp('You pressed "o" key.');
        gate = 1;
        firstClick = 'out';

    end

        % Check for the 'Escape' key to end the script
    [~, ~, keyCode] = KbCheck;
    if keyCode(KbName('Escape'))
        ShowCursor;
        sca;
        endLoopGate=1;
        break;
    end

while KbCheck
% Do nothing, just wait for the key to be released
end

end

end

end


if(endTime<timeLimit)


% Loop through frames
for frame = 1:10
    % Draw a gray screen
     Screen('FillRect', window, dotColor);
    
    % Flip the screen
    Screen('Flip', window);

end


disp("Trial 2 Start");
disp("...................");

if(touchE==1)
TouchQueueCreate(window, dev);
TouchQueueStart(dev);
SetMouse(0,0,window);
HideCursor(window);
end


f=0;


tic
while toc < timeLimit % Run for time limit

    peck=0;

    % Draw the background image
    Screen('DrawTexture', window, backgroundTexture);
    
    % Draw the spinning image onto the background texture at the center of the screen with the current rotation angle
    Screen('DrawTexture', window, spinningTexture, [], ...
        [xPos, yPos, xPos + spinWidth, yPos + spinHeight], rotationAngle);
    
    % Flip the screen to display the changes
    Screen('Flip', window);
    
    % Increment the rotation angle
    rotationAngle = rotationAngle + rotationIncrement;
    if(rotationAngle>360)
        rotationAngle=rotationAngle-360;
    end


    if(touchE==1)
          while TouchEventAvail(dev)
        % Process next touch event 'evt':
        evt = TouchEventGet(dev, window);

        % Touch blob id - Unique in the session at least as
        % long as the finger stays on the screen:
        id = evt.Keycode;

        % Keep the id's low, so we have to iterate over less blobcol slots
        % to save computation time:

          if (evt.Type == 2 ||  evt.Type == 3 ||  evt.Type == 4)
          % New touch point -> New blob!
          blobcol{id}.col = rand(3, 1);
          blobcol{id}.mul = 1.0;
          blobcol{id}.x = evt.MappedX;
          blobcol{id}.y = evt.MappedY;
          blobcol{id}.t = evt.Time;
          % Track time delta in msecs between touch point updates:
          blobcol{id}.dt = 0;

          if(((blobcol{id}.x-pTX)^2 + (blobcol{id}.y-pTY)^2)^0.5>dotSize*touchMove || toc-pTT > touchTime)

            pTT = toc;
            pTX= blobcol{id}.x;
            pTY = blobcol{id}.y;

          peck=1;
          disp("Touch");

          clicks2(end+1, :) = [blobcol{id}.x, blobcol{id}.y, toc];

          if(blobcol{id}.y<screenY && blobcol{id}.x<screenX)

            if(((blobcol{id}.x-dotCenterRX)^2 + (blobcol{id}.y-dotCenterRY)^2)^0.5)
            peckD = ((blobcol{id}.x-dotCenterRX)^2 + (blobcol{id}.y-dotCenterRY)^2)^0.5;
            timeToClick = toc;
            break;
            
          end
          end
          end
        end
    end
    end

     if ~strcmp(timeToClick,'NA')
        break
     end


 % Check for a click on the window
    [~, ~, buttons] = GetMouse(window);
    if any(buttons)
        % Get the clicked coordinates
        [clickX, clickY] = GetMouse(window);

        if(((clickX-pTX)^2 + (clickY-pTY)^2)^0.5>dotSize*touchMove || toc-pTT > touchTime)
        
            pTT = toc;
            pTX= clickX;
            pTY = clickY;
        
        clicks2(end+1, :) = [clickX, clickY, toc];

        disp("Click");

        % Check if the click is inside dotInMask
        if(clickY<screenY && clickX<screenX)
        if( ((clickX-dotCenterRX)^2 + (clickY-dotCenterRY)^2)^0.5 <dotSize+expandDot)
            timeToClick = toc;
            peckX = clickX;
            peckY = clickY;
            peckD = ((clickX-pTX)^2 + (clickY-pTY)^2)^0.5>dotSize*touchMove || toc-pTT > touchTime;

        break;
        end
        end
        end
    end

    [~, ~, keyCode] = KbCheck;


    % Button Triggers
    if any(keyCode([KbName('i')]))
            timeToClick = toc;
            break;
    end

    if any(keyCode([KbName('o')]))
            timeToClick = toc;
            break;
    end

    if any(keyCode([KbName('space')]))
        timeToClick = toc;
        break;
    end

    % Check for the 'Escape' key to end the script
    if keyCode(KbName('Escape'))
        ShowCursor;
        sca;
        endLoopGate=1;
        break;
    end

f=f+1;
end

toc


while KbCheck
% Do nothing, just wait for the key to be released
end

end


% Measures
%...........................................................................
timeToClick2 = timeToClick;
peckX2 = peckX;
peckY2 = peckY;
peckD2 = peckD;

f2 = f; 
rotationAngle2 = rotationAngle;


ba1_2 = rotationAngle+85;
ba2_2 = rotationAngle+85+120;
ba3_2 = rotationAngle+85+240;

if(ba1_2>360)ba1_2=ba1_2-360; end
if(ba2_2>360)ba2_2=ba2_2-360; end
if(ba3_2>360)ba3_2=ba3_2-360; end

bd1_2 = 'NA';
bd2_2 = 'NA';
bd3_2 = 'NA';
minValue2= 'NA';

if strcmp(firstClick,'in')

bd1_2 = abs(ba1_2 - randomAngleI);
if bd1_2 > 180  
bd1_2 = 360 - bd1_2;
end

bd2_2 = abs(ba2_2 - randomAngleI);
if bd2_2 > 180  
bd2_2 = 360 - bd2_2;
end

bd3_2 = abs(ba3_2 - randomAngleI);
if bd3_2 > 180  
bd3_2 = 360 - bd3_2;
end

elseif  strcmp(firstClick,'out')

bd1_2 = abs(ba1_2 - randomAngleO);
if bd1_2 > 180  
bd1_2 = 360 - bd1_2;
end

bd2_2 = abs(ba2_2 - randomAngleO);
if bd2_2 > 180  
bd2_2 = 360 - bd2_2;
end

bd3_2 = abs(ba3_2 - randomAngleO);
if bd3_2 > 180  
bd3_2 = 360 - bd3_2;
end

end


% Define a 1D array
array = [bd1_2,bd2_2,bd3_2];

% Calculate the minimum value
minValue2 = min(array);


disp(['Duration ' num2str(toc) ' seconds']);
disp(['Speed ' num2str(f/toc) ' fps']);

disp("...................");
disp("Trial 2 End");


%Output
%...........................................................................

state1 = "Fail";
if~strcmp(timeToClick1,"NA")
state1 = "Succ";
end


state2 = "Fail";
if~strcmp(timeToClick2,"NA")
state2 = "Succ";
end


% Get the current date and time as a datetime object
currentDateTime = datetime('now');

% Convert datetime object to date string
currentDate = datestr(currentDateTime, 'yyyy-mm-dd');

% Extract the time component as a string
currentTime2 = datestr(currentDateTime, 'HH:MM:SS');

disp(['Current date: ', currentDate]);
disp(['Current time: ', currentTime2]);

dotType1 = 'NA';
dotType2 = 'NA';


if strcmp(firstClick,'in')

timeToClickI = timeToClick1;
timeToClickO = timeToClick2;

peckX_i = peckX1;
peckY_i = peckY1;

peckX_o = peckX2;
peckY_o = peckY2;

dotType1 = 'in';
dotType2 = 'out';

leftRight1= leftRightI

elseif strcmp(firstClick,'out')

timeToClickI = timeToClick2;
timeToClickO = timeToClick1;

peckX_i = peckX2;
peckY_i = peckY2;

peckX_o = peckX1;
peckY_o = peckY1;

dotType1 = 'out';
dotType2 = 'in';

end


% Convert the array to a format with semicolons to separate columns and
% colons to separate rows Clicks1
formattedClicks1 = '';

if(length(clicks1)>0)
% Loop through each row of the array
for i = 1:size(clicks1, 1)
    % Convert the current row to a string with semicolons
    rowString = sprintf('%d %d %d', clicks1(i, 1), clicks1(i, 2), clicks1(i, 3));
    
    % Append a colon to separate rows
    formattedClicks1 = [formattedClicks1, rowString, ';'];
end

% Remove the last colon
formattedClicks1(end) = [];
else

formattedClicks1 = 'NA';

end


% Convert the array to a format with semicolons to separate columns and
% colons to separate rows Clicks2
formattedClicks2 = '';

if(length(clicks2)>0)
% Loop through each row of the array
for i = 1:size(clicks2, 1)
    % Convert the current row to a string with semicolons
    rowString = sprintf('%d %d %d', clicks2(i, 1), clicks2(i, 2), clicks2(i, 3));
    
    % Append a colon to separate rows
    formattedClicks2 = [formattedClicks2, rowString, ';'];
end

% Remove the last colon
formattedClicks2(end) = [];
else

formattedClicks2 = 'NA';

end


colorString = sprintf('%f %f %f', dotColor);


% Combine header and data into a single cell array
header = {'BirdID','TrialType','trainWindImage','shownWindImage' 'Speed', 'Number','Date','Time','FirstPeck', 'TimeToPeck', 'Clicks', 'SuccFail', 'DotType', 'PeckX', 'PeckY','PeckDist', 'leftRight_i','leftRight_o', 'TargetX_i', 'TargetY_i', 'TargetX_o', 'TargetY_o', 'TargetSize', 'TargetAngle_i', 'TargetAngle_o', 'RadiusCentre_i', 'RadiusCentre_o', 'TurbineAngle', 'Blade1Angle', 'Blade2Angle', 'Blade3Angle', 'Blade1Dist', 'Blade2Dist', 'Blade3Dist', 'MinBladeDist','expandTarget','frameRate','dotCol','timeOutTime','frames','screenWidth','screenHeight'}; % Example header

rowData1 = {selectedFolderName,trialType,trainType,bladeType, num2str(speed), num2str(existingTrials+1),currentDate,currentTime1,'first', num2str(timeToClick1), formattedClicks1, state1, dotType1, num2str(peckX1), num2str(peckY1),num2str(peckD1), leftRightI, leftRightO, num2str(dotCenterIX), num2str(dotCenterIY), num2str(dotCenterOX), num2str(dotCenterOY), num2str(dotSize), num2str(randomAngleI), num2str(randomAngleO), num2str(radiusI), num2str(radiusO), num2str(rotationAngle1), num2str(ba1_1), num2str(ba2_1), num2str(ba3_1), num2str(bd1_1), num2str(bd2_1), num2str(bd3_1), num2str(minValue1),num2str(expandDot),num2str(f1/toc),colorString,num2str(timeLimit),num2str(f1),num2str(screenX),num2str(screenY)}; % Example data

rowData2 = {selectedFolderName,trialType,trainType,bladeType, num2str(speed), num2str(existingTrials+1.5),currentDate,currentTime2,'second', num2str(timeToClick2), formattedClicks2, state2, dotType2, num2str(peckX2), num2str(peckY2),num2str(peckD2), leftRightI, leftRightO, num2str(dotCenterIX), num2str(dotCenterIY), num2str(dotCenterOX), num2str(dotCenterOY), num2str(dotSize), num2str(randomAngleI), num2str(randomAngleO), num2str(radiusI),num2str(radiusO), num2str(rotationAngle2), num2str(ba1_2), num2str(ba2_2), num2str(ba3_2), num2str(bd1_2), num2str(bd2_2), num2str(bd3_2), num2str(minValue2),num2str(expandDot),num2str(f2/toc),colorString,num2str(timeLimit),num2str(f2),num2str(screenX),num2str(screenY)}; % Example data

data = [header; rowData1; rowData2];

% Create the filename based on the loop index
if ~strcmp(timeToClick, "NA")
    filename = [num2str(existingTrials+1), '_S', '.csv'];
else
    filename = [num2str(existingTrials+1), '_F', '.csv'];
end

% Open the file for writing
fileID = fopen(fullfile(folderPath, filename), 'w');


% Write the data rows
for i = 1:size(data, 1)
    fprintf(fileID, '%s,', data{i,1:end-1});
    fprintf(fileID, '%s\n', data{i,end});
end

% Close the file
fclose(fileID);


existingTrials = existingTrials + 1; % Corrected variable name


if keyCode(KbName('Escape'))
    ShowCursor;
    endLoopGate=1;
    sca;
    
end
 

end
end

% Define the command to set monitor 1 as the primary display
command = [nircmdPath ' setprimarydisplay ' num2str(secondaryMonitorN)];

% Execute the command
status = system(command);

% Check if the command executed successfully
if status == 0
    disp(['Primary display set']);
else
    disp('Error setting primary display.');
end


% Close the window
sca;
```
