## Supplementary material for "Biologically inspired warning patterns deter birds from wind turbines": LiveSpinTest.m

```
% Clear the workspace and the screen
sca;
close all;
clear;

% Set up Psychtoolbox
PsychDefaultSetup(2);

Screen('Preference', 'SkipSyncTests', 1);
Screen('Preference', 'VisualDebugLevel', 0); 

screenNumber = 1;

% Open a window with a gray background
window = PsychImaging('OpenWindow', screenNumber, [0.5, 0.5, 0.5]);

% Enable alpha blending for transparency
Screen('BlendFunction', window, GL_SRC_ALPHA, GL_ONE_MINUS_SRC_ALPHA);

% Load the background image
backgroundImage = imread('Backgrounds/BlankGray.png'); % Provide the path to your background image

% Get the screen dimensions
[screenX, screenY] = Screen('WindowSize', window);

% Resize the background image to fit the screen dimensions
resizedBackgroundImage = imresize(backgroundImage, [screenY, screenX]);

% Create a texture for the background image
backgroundTexture = Screen('MakeTexture', window, resizedBackgroundImage);

% Load the spinning image with alpha channel
[TurbineBlades, ~, alphaChannel] = imread('Blades/Spiral_Test.png', 'png');

TurbineBlades = imresize(TurbineBlades, [screenY*0.85, screenY*0.85]);
alphaChannel = imresize(alphaChannel, [screenY*0.85, screenY*0.85]);

% Create a texture for the spinning image
spinningTexture = Screen('MakeTexture', window, cat(3, TurbineBlades, alphaChannel));

% Get the size of the spinning image
[spinHeight, spinWidth, ~] = size(TurbineBlades);

% Calculate the position to draw the spinning image at the center of the screen
xPos = (screenX - spinWidth) / 2; % Adjust the x position to center the spinning image on the background
yPos = (screenY - spinHeight) / 2; % Adjust the y position to center the spinning image on the background

% Define initial rotation angle
rotationAngle = 0; % Initial rotation angle in degrees

frameRate = 165;
speed = 20; % RPM

numFrames = frameRate * 60/speed;

rotationIncrement = 360/numFrames; % Rotation increment in degrees

% Get the starting time
startTime = GetSecs;
tic
% Loop until either 360 frames have passed or the escape key is pressed

while rotationAngle < 360 % Run for 2 minutes
    % Draw the background image
    Screen('DrawTexture', window, backgroundTexture);
    
    % Draw the spinning image onto the background texture at the center of the screen with the current rotation angle
    Screen('DrawTexture', window, spinningTexture, [], ...
        [xPos, yPos, xPos + spinWidth, yPos + spinHeight], rotationAngle);
    
    % Flip the screen to display the changes
    Screen('Flip', window);
    
    % Increment the rotation angle
    rotationAngle = rotationAngle + rotationIncrement;
    
    % Check for the escape key press
    [~, ~, keyCode] = KbCheck;
    if keyCode(KbName('Escape'))
        break; % Break the loop if escape key is pressed
    end
end
toc

disp(['Duration ' num2str(toc) ' seconds']);

disp(['Speed ' num2str(numFrames/toc) ' fps']);

% Close the window
sca;
```
