## Supplementary material for "Biologically inspired warning patterns deter birds from wind turbines": MergeCSV.m

```
% Select folder containing CSV files
folder = uigetdir('Select folder containing CSV files');
if folder == 0
    error('Folder not selected.');
end


parts = strsplit(folder, filesep);
parent_path = strjoin(parts(1:end-1), filesep);

[~, folderName, ~] = fileparts(folder);

% Remove forward slashes
folderName = strrep(folderName, '/', '');

% Remove backslashes
folderName = strrep(folderName, '\', '');


disp(parent_path);

% Get list of CSV files
files = dir(fullfile(folder, '*.csv'));

% Initialize cell array to store file numbers
fileNumbers = cell(length(files), 1);

% Extract numbers from file names
for i = 1:length(files)
    [~, name, ~] = fileparts(files(i).name);
    number = sscanf(name, '%d_');
    if ~isempty(number)
        fileNumbers{i} = number;
    end
end

% Remove empty cells
fileNumbers = fileNumbers(~cellfun('isempty', fileNumbers));

% Sort file numbers
[~, idx] = sort(cell2mat(fileNumbers));

% Reorder files
files = files(idx);

% Display the list of CSV files
disp('CSV files in selected folder:');
disp({files.name}');

% Initialize file ID for writing
outputFileName = fullfile(parent_path , [folderName '.csv']);
disp(outputFileName);
fid = fopen(outputFileName, 'w');
if fid == -1
    error('Failed to open file for writing.');
end

% Initialize flag to indicate whether it's the first file
firstFile = true;

% Read the content of each CSV file
for i = 1:length(files)
    % Read the content of the current file as a string
    fileContents = fileread(fullfile(folder, files(i).name));
    
    % Split the string using newline as delimiter
    fileRows = strsplit(fileContents, '\n');
    
    % Remove the first row (header) if it's not the first file
    if ~firstFile
        fileRows(1) = [];
    else
        firstFile = false;
    end
    
    % Remove the last row
    if ~isempty(fileRows)
        fileRows(end) = [];
    end
    
    % Write the rows to the combined CSV file
    for j = 1:length(fileRows)
        fprintf(fid, '%s\n', fileRows{j});
    end
end

% Close the file
fclose(fid);

disp(['Combined data saved to ' outputFileName]);
```
