## Supplementary material for "Biologically inspired warning patterns deter birds from wind turbines": MergeCSV_Birds.m

```
% Select folder containing CSV files
birdsfolder = uigetdir('Select Birds Folders');
if birdsfolder == 0
    error('Folder not selected.');
end

% Get the list of all files and folders in the specified directory
listing = dir(birdsfolder);

% Initialize an empty cell array to store folder names
birdNames = {};

% Iterate through each item in the directory listing
for i = 1:numel(listing)
    % Check if the current item is a directory, starts with 'G24', and not '.' or '..'
    if listing(i).isdir && startsWith(listing(i).name, '24G') && ~strcmp(listing(i).name, '.') && ~strcmp(listing(i).name, '..')
        % If it meets conditions, append its name to the birdNames cell array
        birdNames{end+1} = listing(i).name;
    end
end

% Display the names of folders
disp(birdsfolder);
disp(birdNames);

for B = 1:length(birdNames)

    birdName = birdNames{B};
        birdName  = strrep(birdName , '/', '');
        birdName  = strrep(birdName , '\', '');
    char(birdName);

    TrialFolder = fullfile(birdsfolder,birdNames{B},"/Trials/");

    F0 = ("/0_Static/");
     F1 = ("/1_Slow/");
      F2 = ("/2_Medium/");
       F3 = ("/3_Fast/");
        F4 = ("/4_MultiDot_ExpSpeed/");
         F5 = ("/5_Comparative_Experiment/");

      trialDirectories = {F0,F1,F2,F3,F4,F5};
        

      %Merge Trials
      %........................................................

      for F = 1:length(trialDirectories)

          folder = fullfile(TrialFolder,trialDirectories{F});

                folderName= trialDirectories{F};
                
                % Remove forward slashes
                folderName = strrep(folderName, '/', '');
                
                % Remove backslashes
                folderName = strrep(folderName, '\', '');
                
                
                % Get list of CSV files
                files = dir(fullfile(folder, '*.csv'));
                
                % Initialize cell array to store file numbers
                fileNumbers = cell(length(files), 1);
                
                % Extract numbers from file names
                for i = 1:length(files)
                    [~, name, ~] = fileparts(files(i).name);
                    number = sscanf(name, '%d_');
                    if ~isempty(number)
                        fileNumbers{i} = number;
                    end
                end
                
                % Remove empty cells
                fileNumbers = fileNumbers(~cellfun('isempty', fileNumbers));
                
                % Sort file numbers
                [~, idx] = sort(cell2mat(fileNumbers));
                
                % Reorder files
                files = files(idx);
                
                % Display the list of CSV files
                disp('CSV files in selected folder:');
                disp({files.name}');
                
                
                % Initialize file ID for writing
                outputFileName = [char(TrialFolder) char(folderName) '.csv'];
                
                disp("OutputFileName")
                disp(outputFileName);
                disp("TrialFolder")
                disp(TrialFolder);
                disp("FolderName");
                disp(folderName);
                
                
                fid = fopen(outputFileName, 'w');
                if fid == -1
                    error('Failed to open file for writing.');
                end
                
                % Initialize flag to indicate whether it's the first file
                firstFile = true;
                
                % Read the content of each CSV file
                for i = 1:length(files)
                    % Read the content of the current file as a string
                    fileContents = fileread(fullfile(folder, files(i).name));
                    
                    % Split the string using newline as delimiter
                    fileRows = strsplit(fileContents, '\n');
                    
                    % Remove the first row (header) if it's not the first file
                    if ~firstFile
                        fileRows(1) = [];
                    else
                        firstFile = false;
                    end
                    
                    % Remove the last row
                    if ~isempty(fileRows)
                        fileRows(end) = [];
                    end
                    
                    % Write the rows to the combined CSV file
                    for j = 1:length(fileRows)
                        fprintf(fid, '%s\n', fileRows{j});
                    end
                end
                
                % Close the file
                fclose(fid);
                
                disp(['Combined data saved to ' outputFileName]);
                
                
                      end


      %Merge Training and Experiment Data
      %........................................................

            % Select Training
        
                files = {"0_Static.csv", "1_Slow.csv", "2_Medium.csv", "3_Fast.csv"};
    
                outputFileName = [char(TrialFolder) char(birdName) char("_Training.csv")];
    
                fid = fopen(outputFileName, 'w');
                
                % Initialize flag to indicate whether it's the first file
                firstFile = true;
                
                % Read the content of each CSV file and write to combined CSV file
                for i = 1:length(files)
                % Read the content of the current file as a string
                fileContents = fileread(fullfile(TrialFolder, files{i}));
                
                % Split the string using newline as delimiter
                fileRows = strsplit(fileContents, '\n');
                
                % Remove the first row (header) if it's not the first file
                if ~firstFile
                    fileRows(1) = [];
                else
                    firstFile = false;
                end
                
                % Remove the last row
                if ~isempty(fileRows)
                    fileRows(end) = [];
                end
                
                % Write the rows to the combined CSV file
                for j = 1:length(fileRows)
                    fprintf(fid, '%s\n', fileRows{j});
                end
                end
                
                % Close the file
                fclose(fid);
                
                disp(['Combined data saved to ' outputFileName]);
    
            
            % Select Experiment
        
                files = {"4_MultiDot_ExpSpeed.csv", "5_Comparative_Experiment.csv"};
    
                outputFileName = [char(TrialFolder) char(birdName) char("_Experiment.csv")];
    
                fid = fopen(outputFileName, 'w');
                
                % Initialize flag to indicate whether it's the first file
                firstFile = true;
                
                % Read the content of each CSV file and write to combined CSV file
                for i = 1:length(files)
                % Read the content of the current file as a string
                fileContents = fileread(fullfile(TrialFolder, files{i}));
                
                % Split the string using newline as delimiter
                fileRows = strsplit(fileContents, '\n');
                
                % Remove the first row (header) if it's not the first file
                if ~firstFile
                    fileRows(1) = [];
                else
                    firstFile = false;
                end
                
                % Remove the last row
                if ~isempty(fileRows)
                    fileRows(end) = [];
                end
                
                % Write the rows to the combined CSV file
                for j = 1:length(fileRows)
                    fprintf(fid, '%s\n', fileRows{j});
                end
                end
                
                % Close the file
                fclose(fid);
                
                disp(['Combined data saved to ' outputFileName]);
    
           

end


%Merge Birds
%........................................................

      %Merge Training
      %........................................................

                outputFileName = [char(birdsfolder) char("/Combined_Training.csv")];
    
                fid = fopen(outputFileName, 'w');
                
                % Initialize flag to indicate whether it's the first file
                firstFile = true;
                
                % Read the content of each CSV file and write to combined CSV file
                for i = 1:length(birdNames)
                % Read the content of the current file as a string

                birdName = birdNames{i};
                birdName  = strrep(birdName , '/', '');
                birdName  = strrep(birdName , '\', '');
                char(birdName);

                fileContents = fileread(fullfile(birdsfolder,birdNames{i},[char("/Trials/") char(birdName) char("_Training.csv")]));
                
                % Split the string using newline as delimiter
                fileRows = strsplit(fileContents, '\n');
                
                % Remove the first row (header) if it's not the first file
                if ~firstFile
                    fileRows(1) = [];
                else
                    firstFile = false;
                end
                
                % Remove the last row
                if ~isempty(fileRows)
                    fileRows(end) = [];
                end
                
                % Write the rows to the combined CSV file
                for j = 1:length(fileRows)
                    fprintf(fid, '%s\n', fileRows{j});
                end
                end
                
                % Close the file
                fclose(fid);
                
                disp(['Combined data saved to ' outputFileName]);
    
     %Merge Experiment
      %........................................................

                outputFileName = [char(birdsfolder) char("/Combined_Experiment.csv")];
    
                fid = fopen(outputFileName, 'w');
                
                % Initialize flag to indicate whether it's the first file
                firstFile = true;
                
                % Read the content of each CSV file and write to combined CSV file
                for i = 1:length(birdNames)
                % Read the content of the current file as a string

                birdName = birdNames{i};
                birdName  = strrep(birdName , '/', '');
                birdName  = strrep(birdName , '\', '');
                char(birdName);

                fileContents = fileread(fullfile(birdsfolder,birdNames{i},[char("/Trials/") char(birdName) char("_Experiment.csv")]));
                
                % Split the string using newline as delimiter
                fileRows = strsplit(fileContents, '\n');
                
                % Remove the first row (header) if it's not the first file
                if ~firstFile
                    fileRows(1) = [];
                else
                    firstFile = false;
                end
                
                % Remove the last row
                if ~isempty(fileRows)
                    fileRows(end) = [];
                end
                
                % Write the rows to the combined CSV file
                for j = 1:length(fileRows)
                    fprintf(fid, '%s\n', fileRows{j});
                end
                end
                
                % Close the file
                fclose(fid);
                
                disp(['Combined data saved to ' outputFileName]);
    
            


  
```
