## Supplementary material for "Biologically inspired warning patterns deter birds from wind turbines": MergeCSV_Multi.m

```
% Select multiple CSV files
[files, folder] = uigetfile('*.csv', 'Select CSV files', 'MultiSelect', 'on');
if isequal(files, 0)
    error('No files selected.');
end

% Prompt user for custom name for combined CSV file
prompt = {'Enter custom name for combined CSV file:'};
dlgtitle = 'Custom Name';
dims = [1 50];
definput = {'combined_data'};
name = inputdlg(prompt, dlgtitle, dims, definput);

if isempty(name)
    error('No custom name provided.');
end

customName = char(name);

% Initialize file ID for writing
outputFileName = fullfile(folder, [customName '.csv']);
fid = fopen(outputFileName, 'w');
if fid == -1
    error('Failed to open file for writing.');
end

% Initialize flag to indicate whether it's the first file
firstFile = true;

% Read the content of each CSV file and write to combined CSV file
for i = 1:length(files)
    % Read the content of the current file as a string
    fileContents = fileread(fullfile(folder, files{i}));
    
    % Split the string using newline as delimiter
    fileRows = strsplit(fileContents, '\n');
    
    % Remove the first row (header) if it's not the first file
    if ~firstFile
        fileRows(1) = [];
    else
        firstFile = false;
    end
    
    % Remove the last row
    if ~isempty(fileRows)
        fileRows(end) = [];
    end
    
    % Write the rows to the combined CSV file
    for j = 1:length(fileRows)
        fprintf(fid, '%s\n', fileRows{j});
    end
end

% Close the file
fclose(fid);

disp(['Combined data saved to ' outputFileName]);
```
