## Supplementary figures and images for "Biologically inspired warning patterns deter birds from wind turbines"

### Blank.png

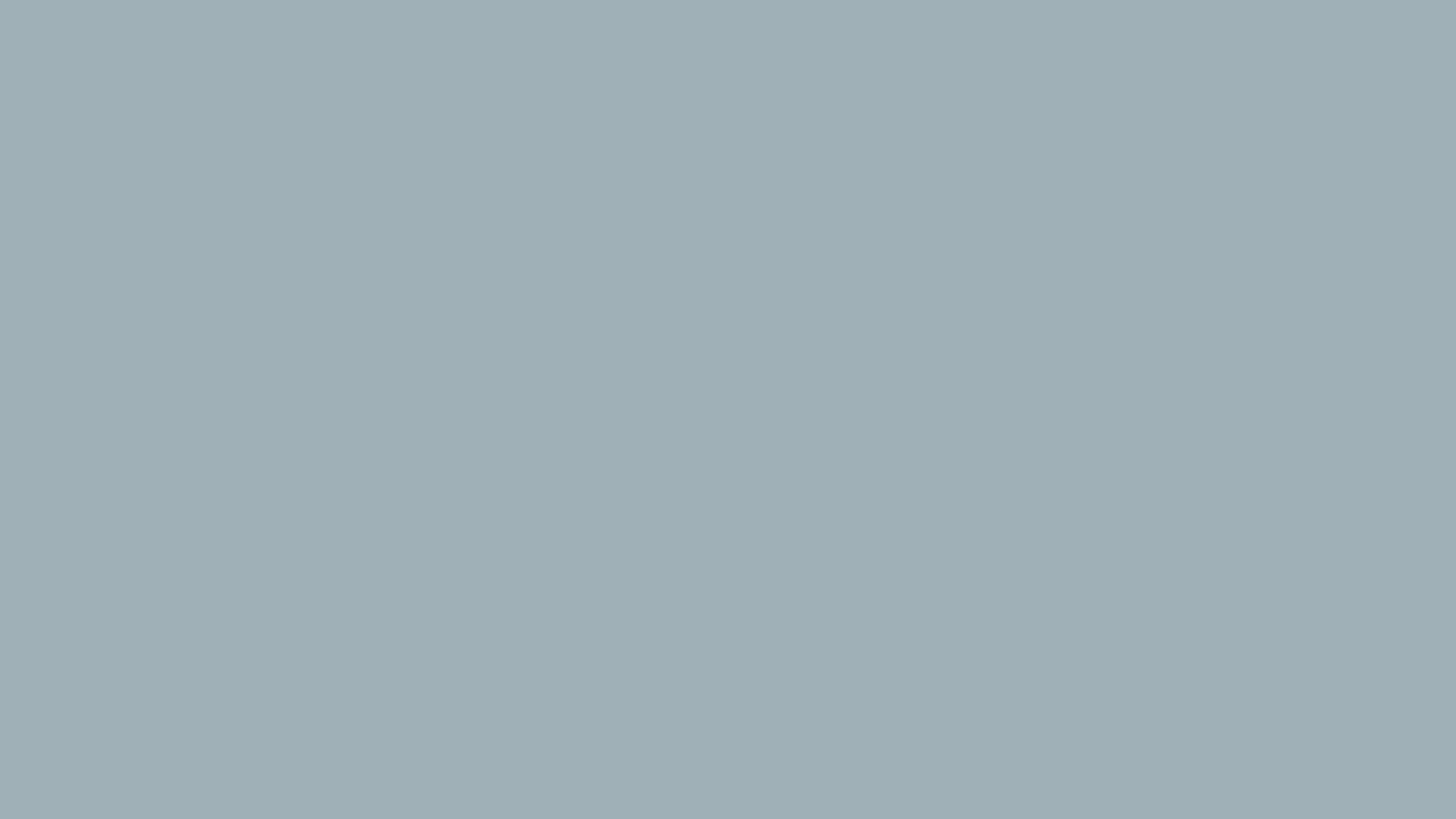

### BlankGray.png

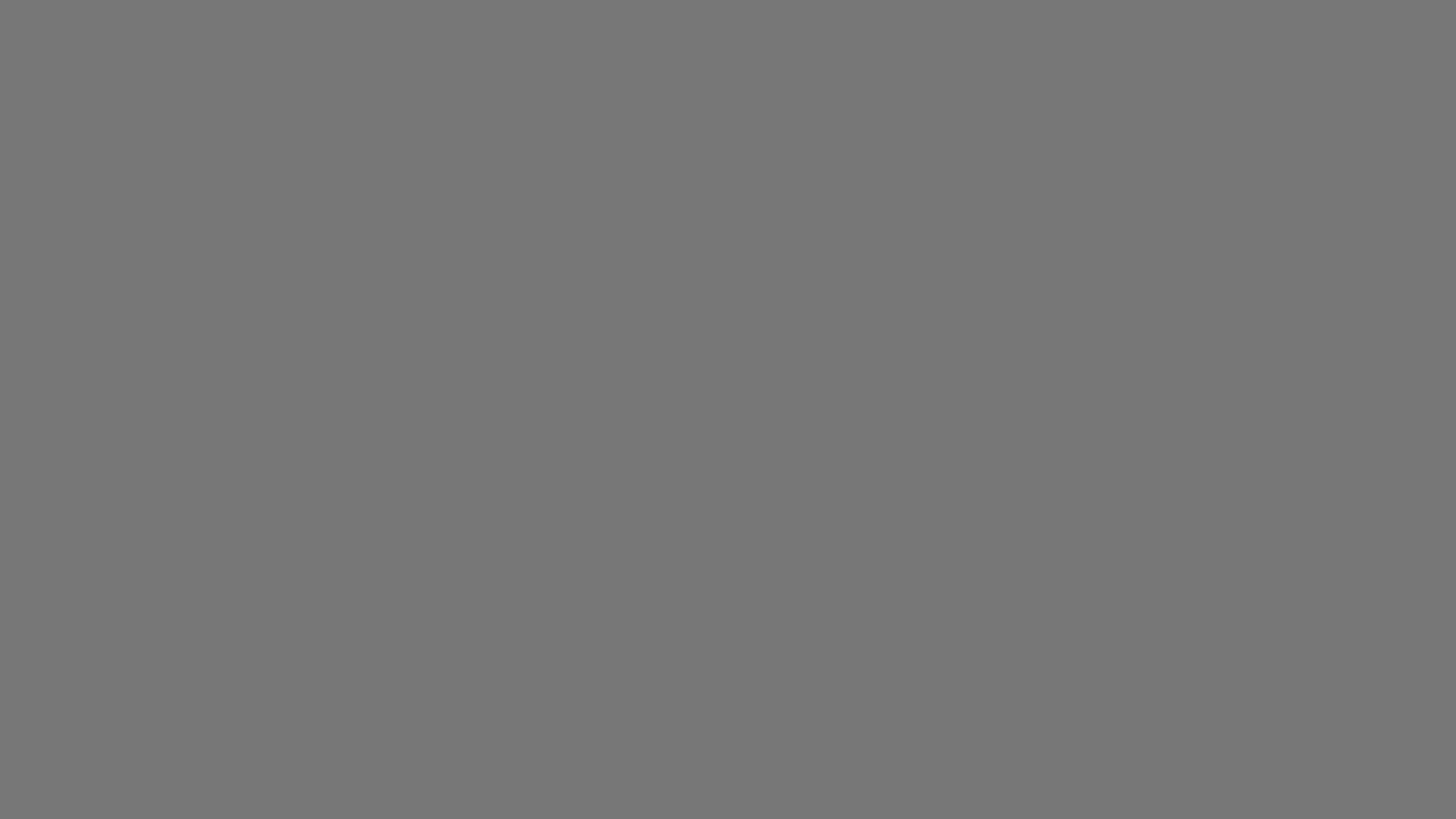

### Spiral_Bio.png

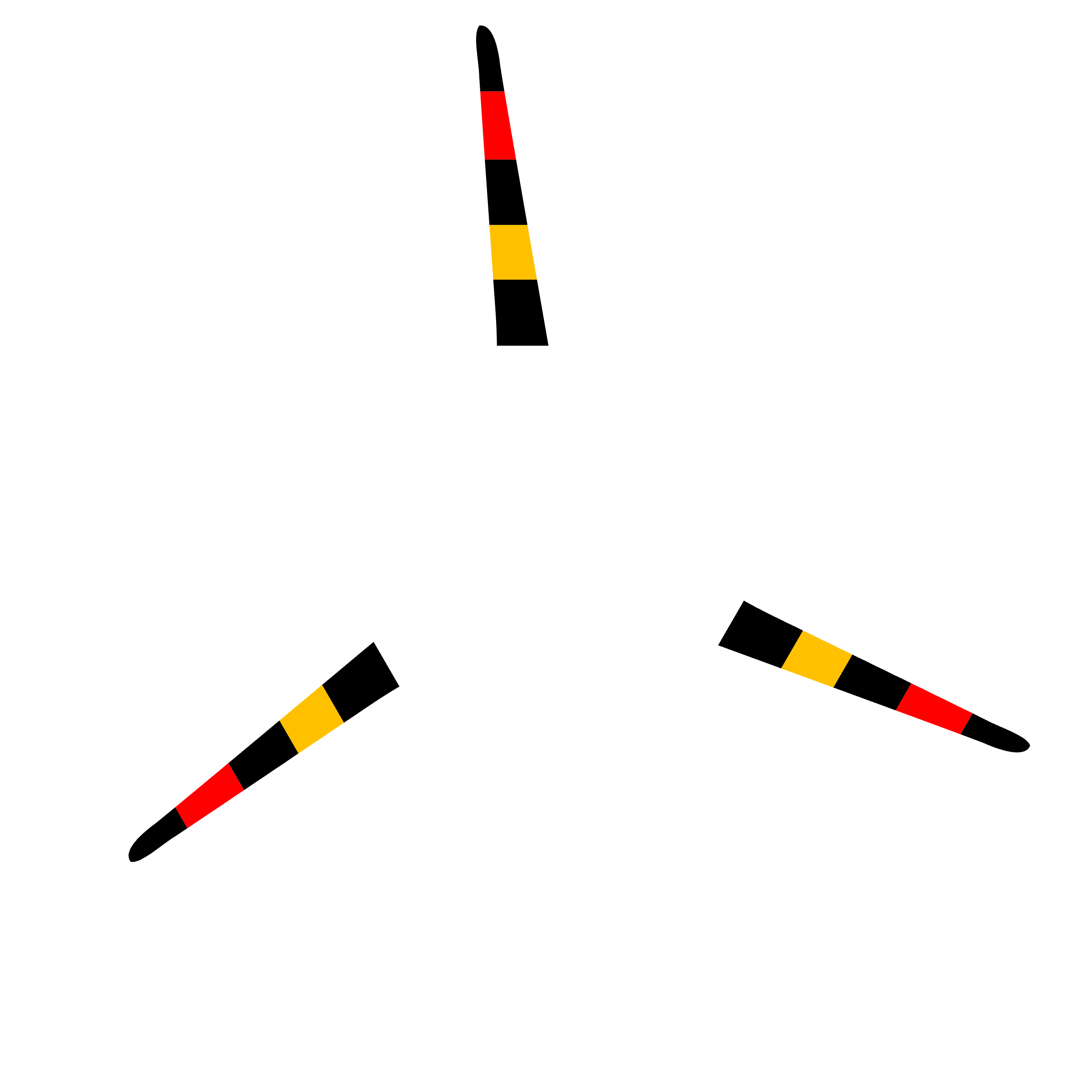

### Spiral_Black.png

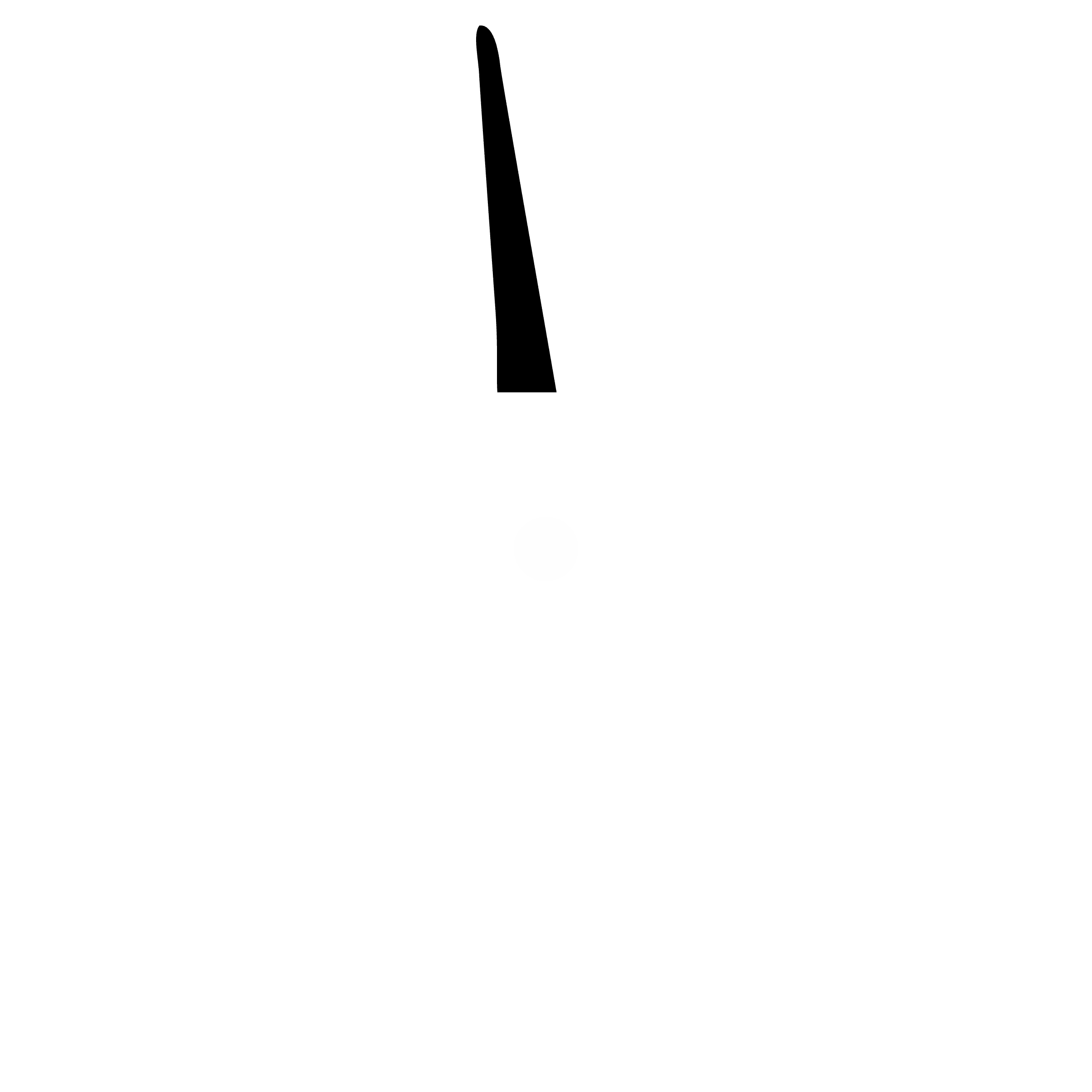

### Spiral_Red.png

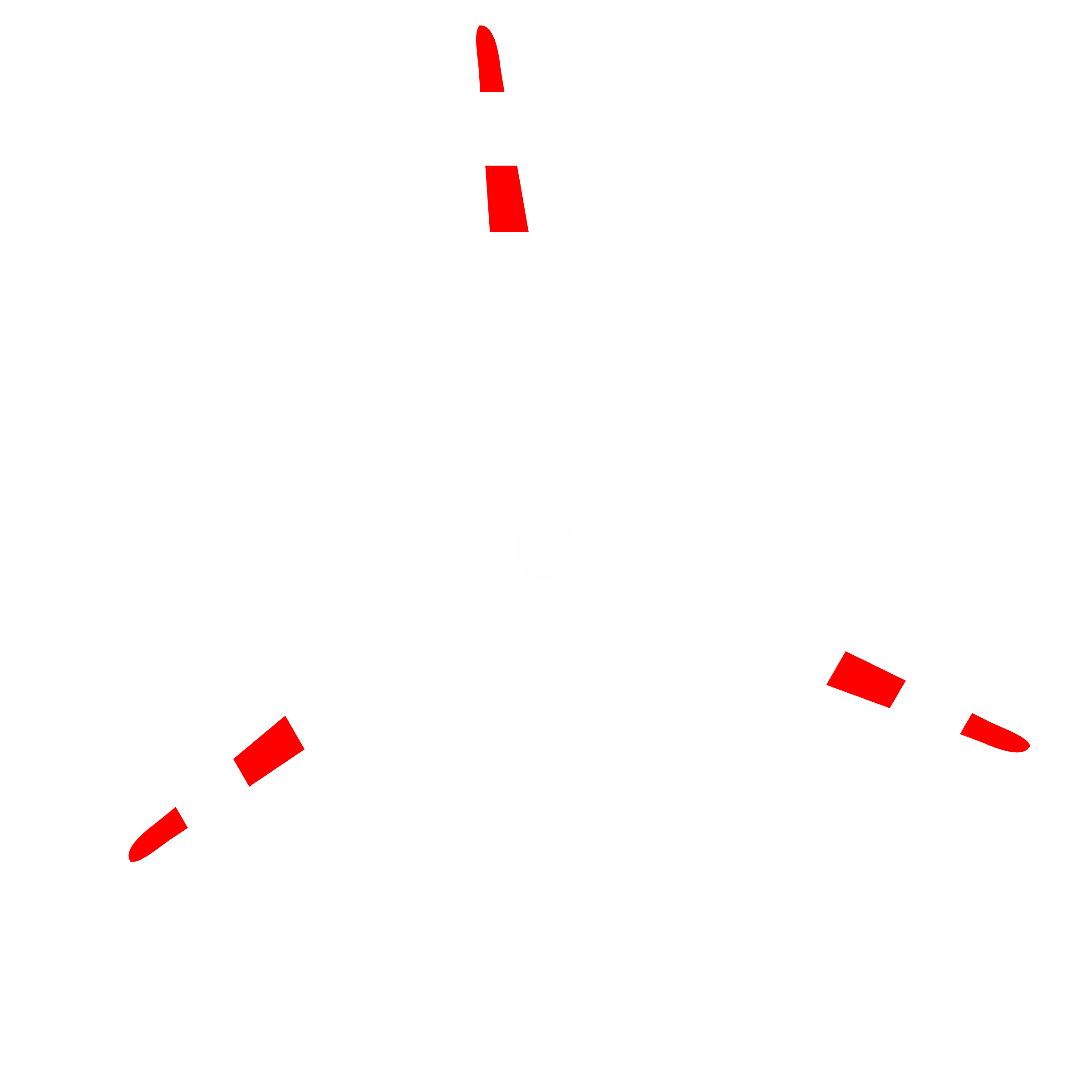

### Spiral_White.png

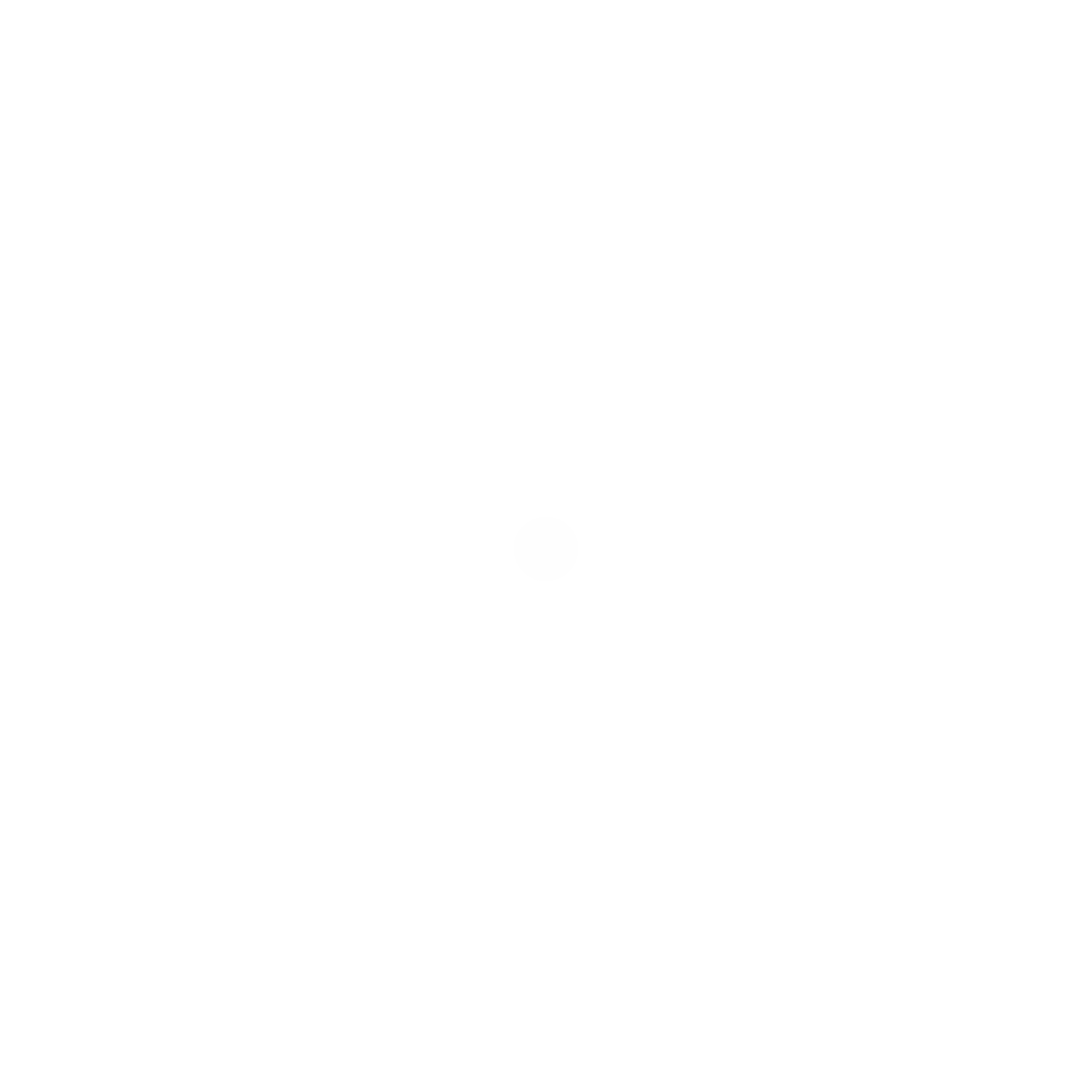

### Turbine.png

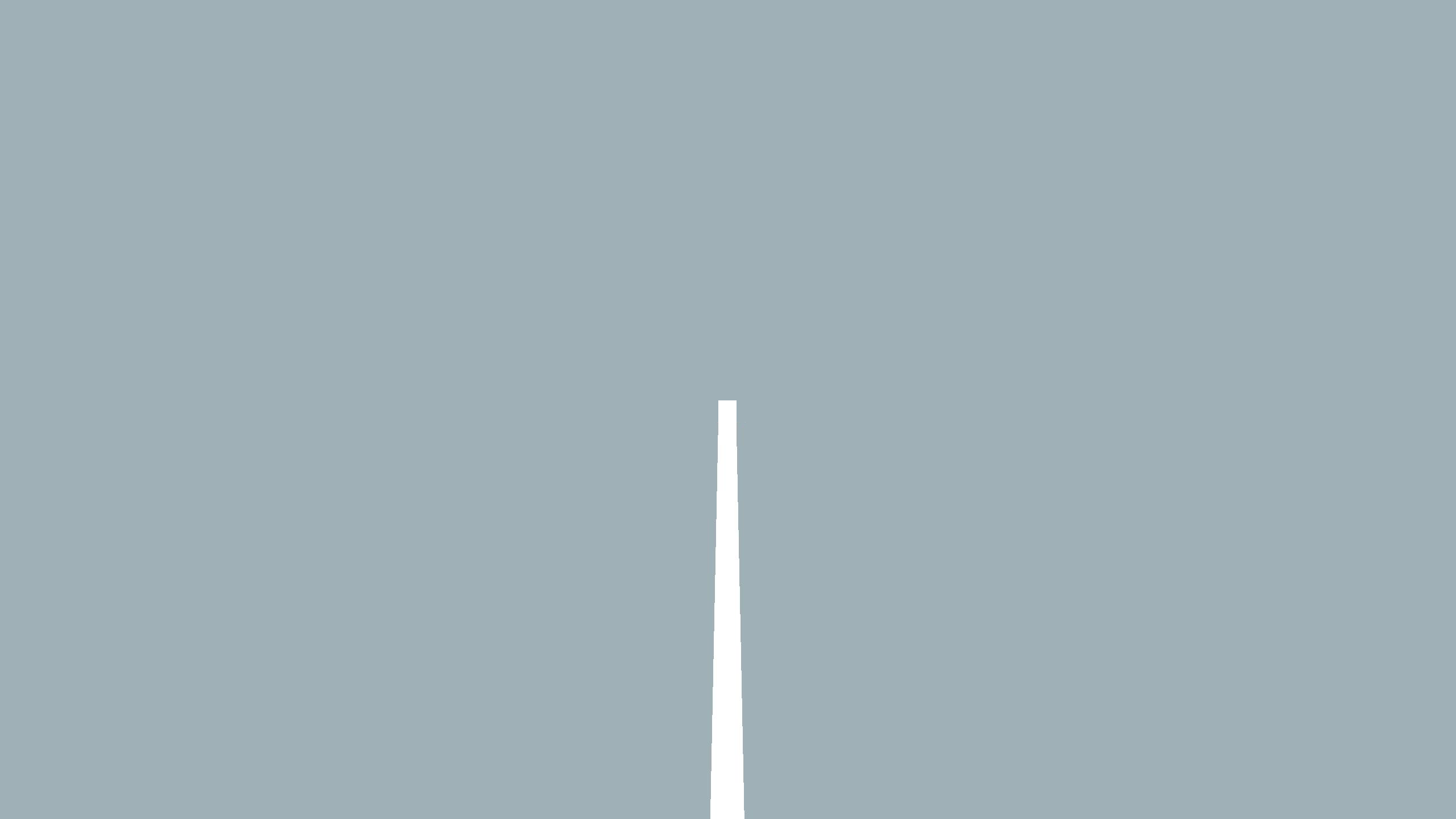
